# Metabolic precision labeling enables selective probing of O-linked *N*-acetylgalactosamine glycosylation

**DOI:** 10.1101/2020.04.23.057208

**Authors:** Marjoke F. Debets, Omur Y. Tastan, Simon P. Wisnovsky, Stacy A. Malaker, Nikolaos Angelis, Leonhard K. R. Moeckl, Junwon Choi, Helen Flynn, Lauren J. S. Wagner, Ganka Bineva-Todd, Aristotelis Antononopoulos, Anna Cioce, William M. Browne, Zhen Li, David C. Briggs, Holly L. Douglas, Gaelen T. Hess, Anthony J. Agbay, Chloe Roustan, Svend Kjaer, Stuart M. Haslam, Ambrosius P. Snijders, Michael C. Bassik, W. E. Moerner, Vivian S. W. Li, Carolyn R. Bertozzi, Benjamin Schumann

## Abstract

Protein glycosylation events that happen early in the secretory pathway are often dysregulated during tumorigenesis. These events can be probed, in principle, by monosaccharides with bioorthogonal tags that would ideally be specific for distinct glycan subtypes. However, metabolic interconversion into other monosaccharides drastically reduces such specificity in the living cell. Here, we use a structure-based design process to develop the monosaccharide probe GalNAzMe that is specific for cancer-relevant Ser/Thr-*N*-acetylgalactosamine (O-GalNAc) glycosylation. By virtue of a branched N-acylamide side chain, GalNAzMe is not interconverted by epimerization to the corresponding N-acetylglucosamine analog like conventional GalNAc-based probes. GalNAzMe enters O-GalNAc glycosylation but does not enter other major cell surface glycan types including Asn (N)-linked glycans. We equip cells with the capacity to biosynthesize the nucleotide-sugar donor UDP-GalNAzMe from a caged precursor. Tagged with a bioorthogonal azide group, GalNAzMe serves as an O-glycan specific reporter in superresolution microscopy, chemical glycoproteomics, a genome-wide CRISPR knock-out (KO) screen, and imaging of intestinal organoids. GalNAzMe is a precision tool that allows a detailed view into the biology of a major type of cancer-relevant protein glycosylation.

**Significance statement:** A large portion of all secreted and cell surface proteins in humans are modified by Ser/Thr(O)-linked glycosylation with *N*-acetylgalactosamine (GalNAc). While of fundamental importance in health and disease, O-GalNAc glycosylation is technically challenging to study because of a lack of specific tools to be used in biological assays. Here, we design an O-GalNAc specific reporter molecule termed GalNAzMe to selectively label O-GalNAc glycoproteins in living human cells. GalNAzMe is compatible with a range of experiments in quantitative biology to broaden our understanding of glycosylation. We further demonstrate that labeling is genetically programmable by expression of a mutant glycosyltransferase, allowing application even to experiments with low inherent sensitivity.

## Introduction

The many facets of cellular glycosylation in health and disease demand methods of visualizing and characterizing glycoconjugates. These methods are essential for the development of next-generation therapeutics and diagnostics that depend on understanding glycosylation of target biomolecules(1, 2). A number of modern experimental techniques have shaped our understanding of biology, such as advanced microscopy, mass spectrometry (MS)-(glyco)proteomics and genome-wide CRISPR-KO screens. Application of these techniques to glycobiology relies on the suitability of detection reagents. Antibodies, lectins, and engineered glycosidases have been instrumental, but are somewhat restricted to sterically accessible epitopes(3–6). Monosaccharides with bioorthogonal, chemically-editable functionalities have allowed a complementary view into glycobiology by entering early glycosylation events (7–11). For instance, the first azide-containing *N*-acetylgalactosamine (GalNAc) analog, GalNAz, and subsequent renditions made it possible to probe core glycosylation that is difficult to reach with protein-based reagents (8). GalNAc analogs are fed to cells as esterase-sensitive precursors and converted into the corresponding uridine diphosphate (UDP) sugar donors by the kinase GALK2 and the pyrophosphorylase AGX1 (Fig. 1A). The cellular glycosylation machinery then incorporates tagged monosaccharides into glycoconjugates where they are reacted with reporter moieties such as fluorophores or biotin by either copper(I)-catalyzed or strain-promoted azide-alkyne cycloaddition (CuAAC or SPAAC, respectively) (12, 13).

**Fig 1:**
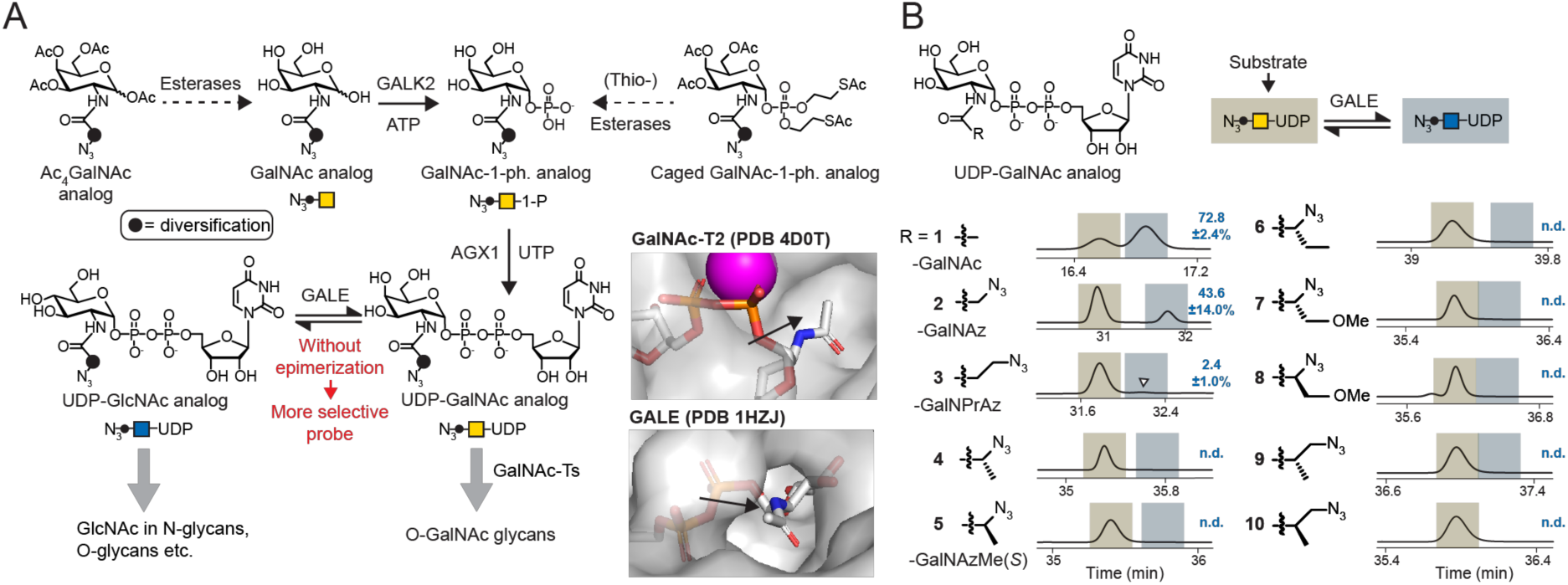
Design of an O-GalNAc specific metabolic labeling reagent. *A*, rationale of probe design. UDP-GalNAc analogs that are not epimerized to the corresponding UDP-GlcNAc derivatives are O-GalNAc specific by design. Derivatives are delivered to the living cell by virtue of per-acetylated or phosphotriester-caged precursors. Compounds with a sterically congested diversification may be resistant to GALE-mediated epimerization but are accepted by GalNAc-Ts. Insert shows UDP-GlcNAc and UDP-GalNAc binding by GalNAc-T2 and GALE, respectively. *B, in vitro* epimerization as assessed by ion-pair HPLC. Retention times of UDP-GalNAc analogs (yellow) and UDP-GlcNAc analogs (blue) are highlighted based on retention times of standards or epimerization reactions with 50-fold higher GALE concentration (*see* Fig. S1B). Arrowhead depicts epimerization of compound **3**. Numbers are % epimerization as assessed by peak integration as means ± SD of three independent replicates or not detected (n.d.). Traces depict relative intensity of absorbance at 260 nm. Data are from one representative out of three independent experiments and were reproduced using lysates of wild type cells as a source of GALE or GALE-KO cells as a negative control in two independent replicates (*see* Fig. S1B).

A particular drawback of most current chemically modified monosaccharides is their low specificity: UDP-GalNAz enters mucin-type (O-GalNAc) glycans but is also converted to the corresponding *N*-acetylglucosamine (GlcNAc) derivative UDP-GlcNAz by the cellular UDP-GlcNAc/GalNAc-4-epimerase (GALE) (Fig. 1A) (14). UDP-GlcNAz then enters N-linked glycans as well as other GlcNAc-containing glycans (14, 15). Other GalNAc analogs are presumably interconverted into GlcNAc analogs in a similar fashion, but their metabolic fate can be variable (16).

Forays have been made into developing reagents that are specific for the structurally simple nucleocytoplasmic O-GlcNAc modification (17–19). However, no such reagents are available to specifically probe the complex cell surface O-GalNAc glycosylation that has fundamental relevance in many aspects of cancer (20, 21).

Studying O-GalNAc glycoproteins by MS-based glycoproteomics is complicated by glycan heterogeneity, the lack of O-glycosylation consensus sequences, and selective enrichment tools. Further complexity is added by the interplay of 20 GalNAc transferase (GalNAc-T1…T20) isoenzymes that mediate the first O-GalNAc biosynthesis step (22, 23). In a “bump-and-hole” (BH) approach, we have recently engineered GalNAc-Ts to carry a double mutation to preferentially accept UDP-GalNAc analogs with bulky chemical, editable tags (24, 25). Although this technique produced bioorthogonal reporters with great specificity for particular GalNAc-T isoenzymes, epimerization of GalNAc analogs by GALE was still a challenge and resulted in background N-glycan labeling (25).

A frequently-used strategy to visualize O-GalNAc glycans is the use of GalNAz in GALE-deficient cells that cannot epimerize UDP-GalNAz (14, 26). However, this strategy is of limited use as GALE deficiency heavily interferes with glycan metabolism and might therefore not be easily adaptable to multicellular model systems such as organoids (27, 28).

Here, we report the first GalNAc-specific, bioorthogonal metabolic labeling reagent *N*-(*S*)-azidoalaninyl galactosamine (GalNAzMe). Using a collection of synthetic azide-containing UDP-GalNAc analogs and structure-informed probe design, we find that branched acylamide side chains confer resistance to GALE-mediated epimerization. We use a caged precursor of the nucleotide-sugar UDP-GalNAzMe to probe O-GalNAc glycosylation in a range of experimental conditions, including superresolution microscopy, chemical MS-glycoproteomics, a genome-wide CRISPR-KO screen and intestinal organoid imaging. GalNAzMe labeling can be enhanced in the presence of a BH-GalNAc-T2 double mutant, further expanding the use of this monosaccharide in glycobiology experiments. Precision tools such as GalNAzMe are essential to uncover the fine details of cellular glycosylation.

## Results

### Probe design

We envisioned that a chemically modified UDP-GalNAc analog would be O-GalNAc-specific if it was i) not epimerized to the UDP-GlcNAc analog by GALE and ii) used by either wild type (WT) or BH-engineered GalNAc-Ts to be incorporated into cell surface O-GalNAc glycans (Fig. 1A). Investigation of the co-crystal structure of human GalNAc-T2 and UDP-GalNAc suggested that the GalNAc acetamide group is embedded in a pocket that allows for some three-dimensional freedom (Fig 1A). A similar binding site architecture was observed in the crystal structures of GalNAc-T10 and GalNAc-T7 with GalNAc or UDP-GalNAc in their active sites, respectively (Fig. S1A) (29, 30). In contrast to GalNAc-Ts, human GALE accommodates the acetamide in a long, narrow cavity, as evidenced in a co-crystal structure of GALE with UDP-GlcNAc (Fig. 1A). This difference in substrate recognition prompted us to explore the chemical determinants of GALE-mediated epimerization by *in vitro* assays. We expressed human GALE in insect cells and used a collection of UDP-GalNAc **1** as well as analogs **2**-**10** with azide-containing acylamide groups as substrates for epimerization (24, 31). Ion-pair high performance liquid chromatography (IP-HPLC) was used to separate the UDP-GalNAc analogs from their UDP-GlcNAc epimers (32). UDP-GalNAc **1**, UDP-GalNAz **2**, and UDP-*N*-3-azidopropionate **3** which we term UDP-GalNPrAz, were epimerized to the corresponding UDP-GlcNAc derivatives (Fig. 1B). In contrast, all compounds containing a branched acylamide moiety (**4-10**) were resistant towards epimerization under these conditions, evident by the absence of a peak with a later retention time in HPLC chromatograms. To rule out co-elution of both epimers, we used commercial and newly synthesized UDP-GlcNAc-derived epimers of **1** (UDP-GlcNAc), **2** (UDP-GlcNAz), **3** (UDP-GlcNPrAz) and **5** (UDP-GlcNAzMe) as standards and confirmed a marked difference in retention time (Fig. S1B). GALE-mediated epimerization of linear, but not branched UDP-GalNAc analogs was corroborated by performing the reactions in the presence of cytosolic extracts of K-562 cells with or without functional GALE (control-sgRNA or GALE-KO, respectively) (25). An extract containing GALE epimerized compounds **1**-**3**, but not **4**-**10**, whereas an extract from GALE-KO cells was devoid of epimerization in all cases (Fig. S1C). When assessing the scope of GALE reactivity, we succeeded in forcing branched analogs **4**-**9** to epimerize by increasing the concentration of purified GALE 50-fold *in vitro* (Fig. S1C). These data indicate that branched acylamide side chains confer resistance to epimerization unless the concentration of GALE is increased to unphysiologically high levels.

We then chose one of the structurally simplest branched UDP-GalNAc analogs in our collection to assess turnover by GalNAc-Ts. We had previously found UDP-GalNAzMe **5** to be a substrate of WT-GalNAc-T2 (24), and confirmed acceptance by WT-GalNAc-T1, T7, and T10 in *in vitro* glycosylation experiments of peptide substrates (Fig. S2A) (24). UDP-GalNAzMe **5** displayed a very similar activity profile to the well-known substrates UDP-GalNAc **1** and UDP-GalNAz **2**, albeit at lower incorporation levels. The azide-containing molecules **2** and **5** were used by WT-T1 and T2 to glycosylate proteins in a membrane protein preparation, as visualized by CuAAC with a biotin-alkyne and fluorescently labeled streptavidin by western blot (Fig. S2C). These data indicate that UDP-GalNAzMe is a viable substrate for GalNAc-Ts to generate azide-tagged O-GalNAc glycans.

### Labeling the cellular O-GalNAc glycome

We then opted to deliver UDP-GalNAzMe **5** into the living cell. Our initial attempts of using a per-acetylated precursor failed, as we did not observe **5** in cell lysates by high-performance anion exchange chromatography with pulsed amperometric detection (HPAEC-PAD). This was in line with previous findings on the low promiscuity of both endogeneous biosynthetic enzymes GALK2 and AGX1 towards chemically modified substrate analogs (16, 25, 33). Mutants of AGX1 with enlarged active sites have been used by us and Yu et al. to successfully transform analogs of GlcNAc-1-phosphate or GalNAc-1-phosphate into the corresponding UDP-sugars and bypass the GALK2 phosphorylation step (*see* Fig. 1A) (25, 34). We thus synthesized a caged, membrane-permissive version of GalNAzMe-1-phosphate **11** (Fig. 2A) (25) and equipped cells with the capacity to biosynthesize UDP-GalNAzMe **5** (*see* Fig. 1A). We transfected HEK293T cells with single and double mutants of the AGX1 active site residues Phe381 and Phe383 (Fig. 2A). Feeding these cells with caged GalNAzMe-1-phosphate **11** led to a peak corresponding to UDP-GalNAzMe **5** only when AGX1^F383A^ termed “mut-AGX1” (25) was present (Fig. 2A, Fig. S2B). This result was somewhat surprising in the context of our previous finding that both AGX1^F383A^ and AGX1^F383G^ accepted a different chemically modified GalNAc-1-phosphate analog (25). When UDP-GalNAzMe **5** was biosynthesized from precursor **11**, we never observed a peak with the retention time of UDP-GlcNAzMe in two different cell lines (Fig. S2B). As a control, Ac_4_GalNAz feeding generated an approximate 3:8 equilibrium between UDP-GalNAz and UDP-GlcNAz even without overexpression of AGX1 (Fig. 2A) (14). Collectively, these data indicate that UDP-GalNAzMe **5** can be biosynthesized by mut-AGX1 in living cells and is not epimerized by endogenous GALE.

**Fig 2:**
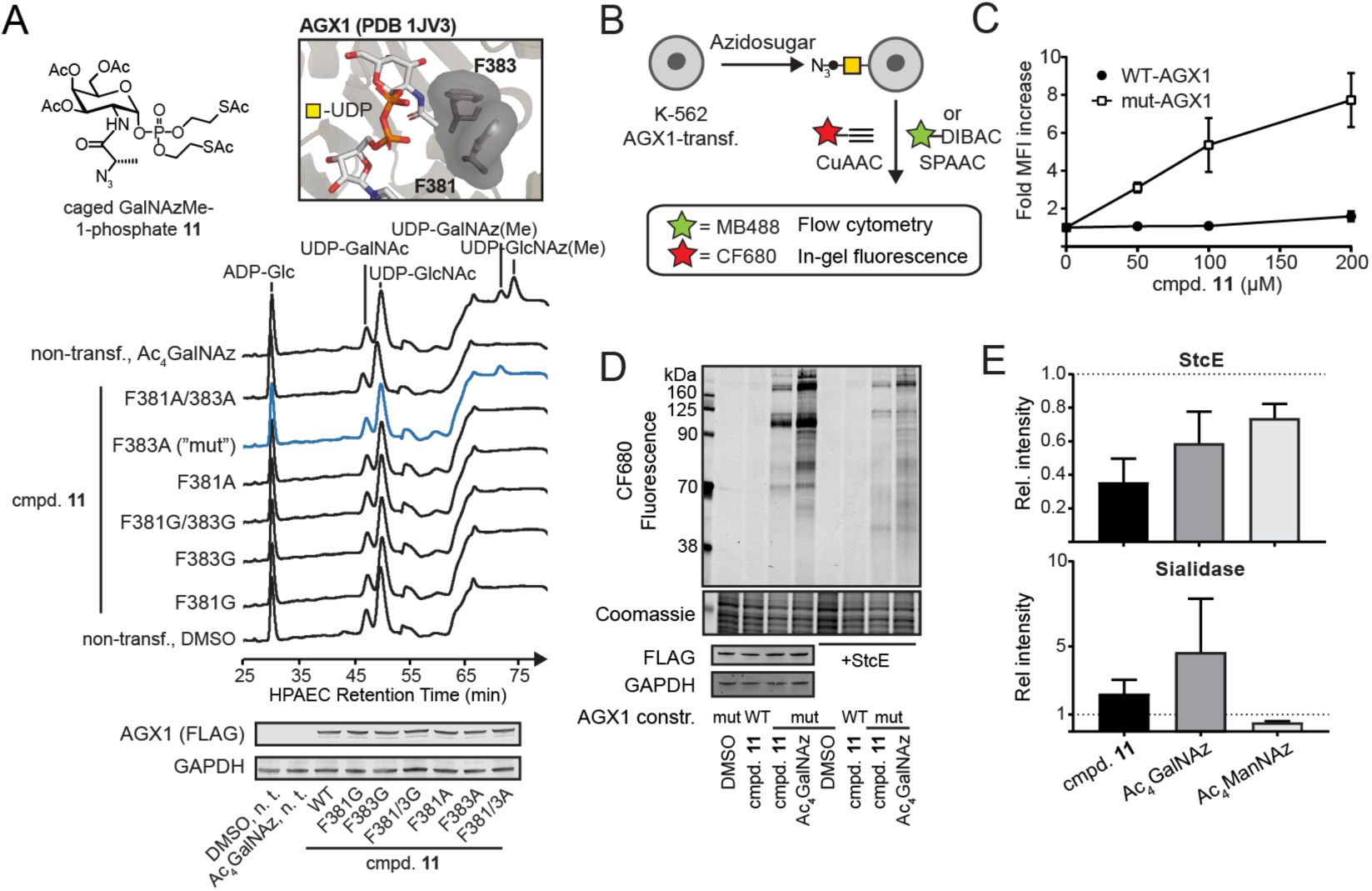
GalNAzMe can be used to label the cell surface glycoproteome. *A*, biosynthesis of UDP-GalNAzMe by mut-AGX1. HEK293T cells were transiently transfected with plasmids encoding for different AGX1 constructs or left non-transfected. Cells were fed with 200 µM compound **11** or Ac_4_GalNAz, and cell lysates were analyzed by HPAEC-PAD. *B*, cell surface labeling workflow using either CuAAC or SPAAC. *C*, dose dependence of GalNAzMe labeling by K-562 cells stably expressing WT-AGX1 or mut-AGX1, as assessed by flow cytometry. Data are mean ± SD from three independent replicates. *D*, cell surface mucin labeling by GalNAzMe and GalNAz. K-562 cells stably expressing WT-AGX1 or mut-AGX1 were fed with DMSO, 3 µM Ac_4_GalNAz, or 100 µM compound **11** and treated with CF680-alkyne as outlined in *B*. Cells were optionally treated with 50 nM StcE before the click reaction. Data are from one representative out of two independent experiments. *E*, cells were treated with either StcE or *V. cholerae* sialidase, then treated with MB488-DIBAC as outlined in *B*, and glycosylation was assessed by flow cytometry. Data are mean + SD of three independent experiments.

We next assessed incorporation of GalNAzMe into cell surface glycans. K-562 cells stably transfected with WT- or mut-AGX1 were treated with either caged GalNAzMe-1-phosphate **11**, Ac_4_GalNAz or DMSO. Azide-containing glycans on the surface of living cells were reacted with clickable (by CuAAC or SPAAC) fluorophores and visualized by flow cytometry or in-gel fluorescence imaging (Fig. 2C-E) (12, 13). Caged GalNAzMe-1-phosphate **11** exhibited dose- (Fig. 2C) and time-dependent (Fig. S3A) incorporation when cells expressed mut-AGX1, but not WT-AGX1. Our data confirmed that UDP-GalNAzMe **5** must be biosynthesized for fluorescent labeling to be detectable, thereby ruling out non-specific incorporation (35).

To elucidate the nature of azide-labeled cell surface glycans, we compared the glycoprotein patterns labeled with GalNAz or GalNAzMe by in-gel fluorescence. Feeding caged GalNAzMe-1-phosphate **11** labeled a subset of the glycoprotein bands of Ac_4_GalNAz (Fig. 2D), consistent with UDP-GalNAz **2** being epimerized and entering GlcNAc-containing glycans. The same behavior was observed in HepG2 cells (Fig. S3B). To assess labeling specificity, we also tested glycoprotein susceptibility towards hydrolytic enzymes. We treated samples with the mucinase StcE that specifically digests highly O-GalNAcylated mucin domains, or with sialidase that removes sialic acid from glycoconjugates (36). Following StcE treatment, the most intense bands labeled by both caged GalNAzMe-1-phosphate **11** and Ac_4_GalNAz feeding had disappeared. The remaining band pattern was much more complex in samples from Ac_4_GalNAz-than from **11**-fed cells (Fig. 2D). Flow cytometry confirmed that StcE treatment decreased the overall labeling intensity of cells fed with either caged GalNAzMe-1-phosphate **11**, Ac_4_GalNAz, or the azide-tagged sialic acid precursor Ac_4_ManNAz (Fig. 2E). In contrast, sialidase treatment led to an increase of labeling with both **11** and Ac_4_GalNAz, presumably due to better accessibility by the click reagents to the azide-tagged glycan structures without sialic acid. The labeling intensity after feeding Ac_4_ManNAz was reduced by sialidase treatment (Fig. 2E, Fig. S3C). These data suggest that GalNAzMe enters the mucin subset of GalNAz-modified glycoproteins, and neither GalNAc derivative substantially enters the sialic acid pool.

We next confirmed that GalNAzMe specifically enters O-GalNAc glycosylation in living cells. We used mut-AGX1-transfected GALE-KO K-562 cells or the corresponding control cells carrying a non-coding single guide (sg)RNA (25). In GALE-KO cells, GalNAz and GalNAzMe should enter the exact same subset of glycans. In cells expressing GALE, UDP-GalNAz **2** should be epimerized and label more cellular glycoproteins than UDP-GalNAzMe **5** (Fig. 3A). We first profiled UDP-sugar levels by HPAEC-PAD in azidosugar-fed cells. As predicted, UDP-GalNAz **2** and UDP-GlcNAz (from the precursor Ac_4_GlcNAz) were not epimerized in GALE-KO cells while epimerization occurred in GALE-expressing cells (Fig. 3B) (25). UDP-GalNAzMe **5** levels were equal in both cell lines fed with **11** and no epimerization was observed irrespective of the presence of GALE. To confirm that these azidosugars enter glycans, we performed a competition experiment in GALE-KO cells by flow cytometry. We used the free sugars GalNAc and GlcNAc to compete with metabolic labeling, and SPAAC to fluorescently detect azide-containing glycoproteins (Fig. 3C-D). Cells fed with both Ac_4_GalNAz and caged GalNAzMe-1-phosphate **11** lost fluorescence intensity in the presence of increasing concentrations of GalNAc, while only Ac_4_GlcNAz labeling was abrogated by an excess of GlcNAc (Fig. 3C).

**Fig 3:**
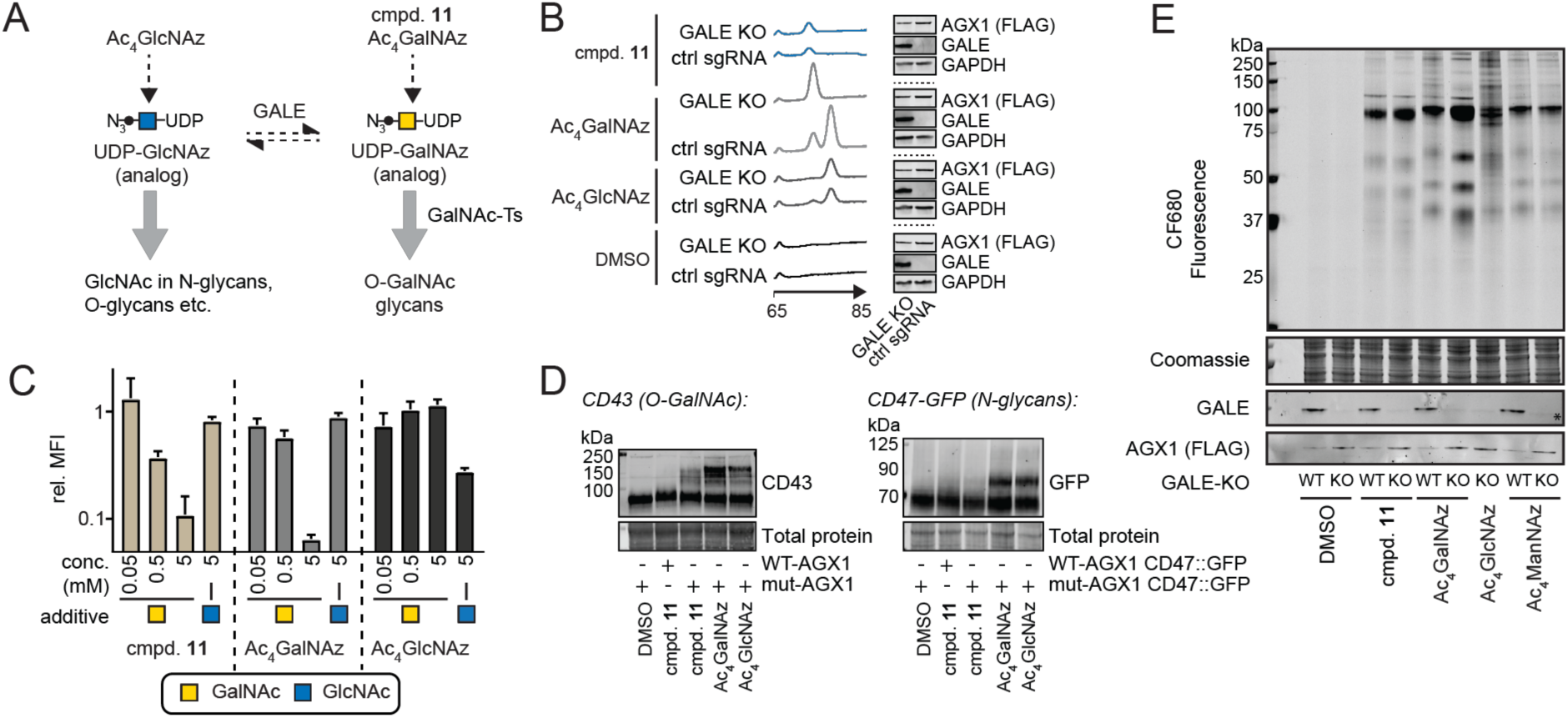
UDP-GalNAzMe is not epimerized and labeled a subset of the UDP-GalNAz-modified glycoproteome. *A*, schematic of the pathways probed herein. Both GalNAc-1-phosphate analog **11** and Ac_4_GalNAz are precursors for O-GalNAc glycosylation. *B*, UDP-GalNAzMe is not epimerized in the living cell while UDP-GalNAz and UDP-GlcNAz are epimerized. K-562 GALE-KO and control cells stably transfected with mut-AGX1 were treated with 200 µM compound **11**, Ac_4_GalNAz, DMSO, or Ac_4_GlcNAz, and UDP-sugar biosynthesis was assessed by HPAEC-PAD. *C*, GALE-KO cells were treated with 100 µM compound **11**, 10 µM Ac_4_GalNAz, or 10 µM Ac_4_GlcNAz, and supplemented with GalNAc or GlcNAc in the indicated concentrations. Cell surface labeling was assessed by flow cytometry after SPAAC using MB488-DIBAC, and fluorescence intensity was normalized to DMSO-treated cells. Data are mean + SD from three independent experiments. *D*, left: K-562 cells stably expressing WT- or mut-AGX1 were fed with DMSO, 100 µM compound **11**, 3 µM Ac_4_GalNAz, or 8 µM Ac_4_GlcNAz, and subjected to PEG mass tagging. K-562 cells stably expressing WT- or mut-AGX1 and GFP::CD47 were fed with DMSO, 100 µM compound **11**, 3 µM Ac_4_GalNAz, or 8 µM Ac_4_GlcNAz, and subjected to PEG mass tagging. *E*, cells were fed with compounds as in *D*, live cells were treated with CF680-alkyne under CuAAC conditions, and proteins in cell lysates were visualized by in-gel fluorescence. Ac_4_ManNAz (0.5 µM) was used as a positive control.

We then assessed glycosylation of discrete *bona fide* O-GalNAc-glycosylated or N-glycosylated proteins with azidosugars. CD43, the most abundant cell surface glycoprotein on K-562 cells, is heavily O-GalNAc glycosylated (37). In contrast, CD47 contains six potential N-glycosylation sites and no predicted O-GalNAc glycans (38). We fed normal or CD47-GFP-overexpressing K-562 cells with caged GalNAzMe-1-phosphate **11**, Ac_4_GalNAz, Ac_4_GlcNAz, or DMSO. Cell lysis and subsequent conjugation with an azide-reactive 10 kDa PEG chain by SPAAC led to a mass shift visible by western blot whenever the azido-sugar was incorporated (39, 40). We observed a clear mass shift in CD43 after feeding GalNAzMe-1-phosphate **11**, Ac_4_GalNAz, or Ac_4_GlcNAz to WT K-562 cells (Fig. 3D). The mass shift induced by GalNAzMe-1-phosphate **11** was only observed when mut-AGX1 was expressed. The Ac_4_GlcNAz-induced mass shift was lost in GALE-KO cells, confirming that these cells could not generate UDP-GalNAz from UDP-GlcNAz (Fig. S4A). A mass shift in overexpressed CD47-GFP was only seen in lysates of cells fed with Ac_4_GalNAz or Ac_4_GlcNAz, but not with caged GalNAzMe-1-phosphate **11** (Fig. 3D). CD43 was labeled by **11** in the same cell line (Fig. S4B).

In-gel fluorescence confirmed that caged GalNAzMe-1-phosphate **11** and Ac_4_GalNAz led to identical band patterns of glycoproteins in GALE-KO cells (Fig. 3E). Strikingly, Ac_4_GlcNAz feeding of GALE-KO cells led to a diffuse pattern of low-intensity glycoprotein bands that resembled the background bands of WT cells fed with Ac_4_GalNAz. Furthermore, the GalNAzMe labeling pattern was not influenced by the presence or absence of GALE. Taken together, these data indicate that UDP-GalNAzMe **5** exclusively enters O-GalNAc glycans, while UDP-GalNAz **2** is epimerized and additionally enters GlcNAc-containing glycans.

To further structurally confirm that UDP-GalNAzMe **5** is not accepted as a substrate by GALE, but is accepted by GalNAc-Ts such as GalNAc-T2, we computationally docked UDP-GalNAzMe into the active sites of both enzymes. We found that the energy-minimized conformation would place the 2-azidopropioamide side chain closer (2.7 Å and 2.9 Å) than the N-C van der Waals radius of 3.3 Å from nearby amino acid side chains in GALE (Fig. S4C). In contrast, UDP-GalNAzMe was accommodated in GalNAc-T2 without such steric clashes.

### GalNAzMe as an O-GalNAc specific reporter molecule

We obtained mass spectrometric evidence for incorporation of GalNAzMe into O-GalNAc glycans. We first confirmed that global cell surface N- and O-glycome profiles of K-562 cells fed with either caged GalNAzMe-1-phosphate **11** or Ac_4_GalNAz did not differ substantially (Fig. S5). We then used chemical MS-glycoproteomics to assess the incorporation of GalNAzMe into cell surface O-GalNAc glycans. Biotin-containing, acid-cleavable alkynyl probe **12** served to enrich azide-containing glycoproteins from the de-N-glycosylated secretome of HepG2 cells. Samples were digested with Lysyl endopeptidase (LysC) after enrichment on Lys-dimethylated Neutravidin beads with enhanced LysC resistance (41). Following glycopeptide release, tandem MS was used to sequence glycopeptides. Higher-energy collisional dissociation (HCD) served to characterize glycan-derived ions, and spectra containing the ions for GalNAzMe (343.1617 m/z) and GalNAz (329.1461 m/z) triggered corresponding electron-transfer dissociation (ETD) to sequence peptides (25). All spectra were manually validated. Both GalNAzMe and GalNAz were found as peptide-proximal residues in O-GalNAc glycans (Fig. 4A, Data S1) and were extended by the downstream glycosylation machinery. For instance, biosynthetic considerations allowed the assignment of the disaccharide β-Gal-(1-3)-α-GalNAzMe-(Thr*) on the glycopeptide TTPPT*TATPIR of human fibronectin, along with other glycoforms and even a di-glycosylated peptide TTPPT*T*ATPIR (Data S1). DMSO feeding did not lead to discernible signal. Taken together, GalNAzMe is a substitute of the peptide-proximal O-GalNAc residue.

**Fig. 4:**
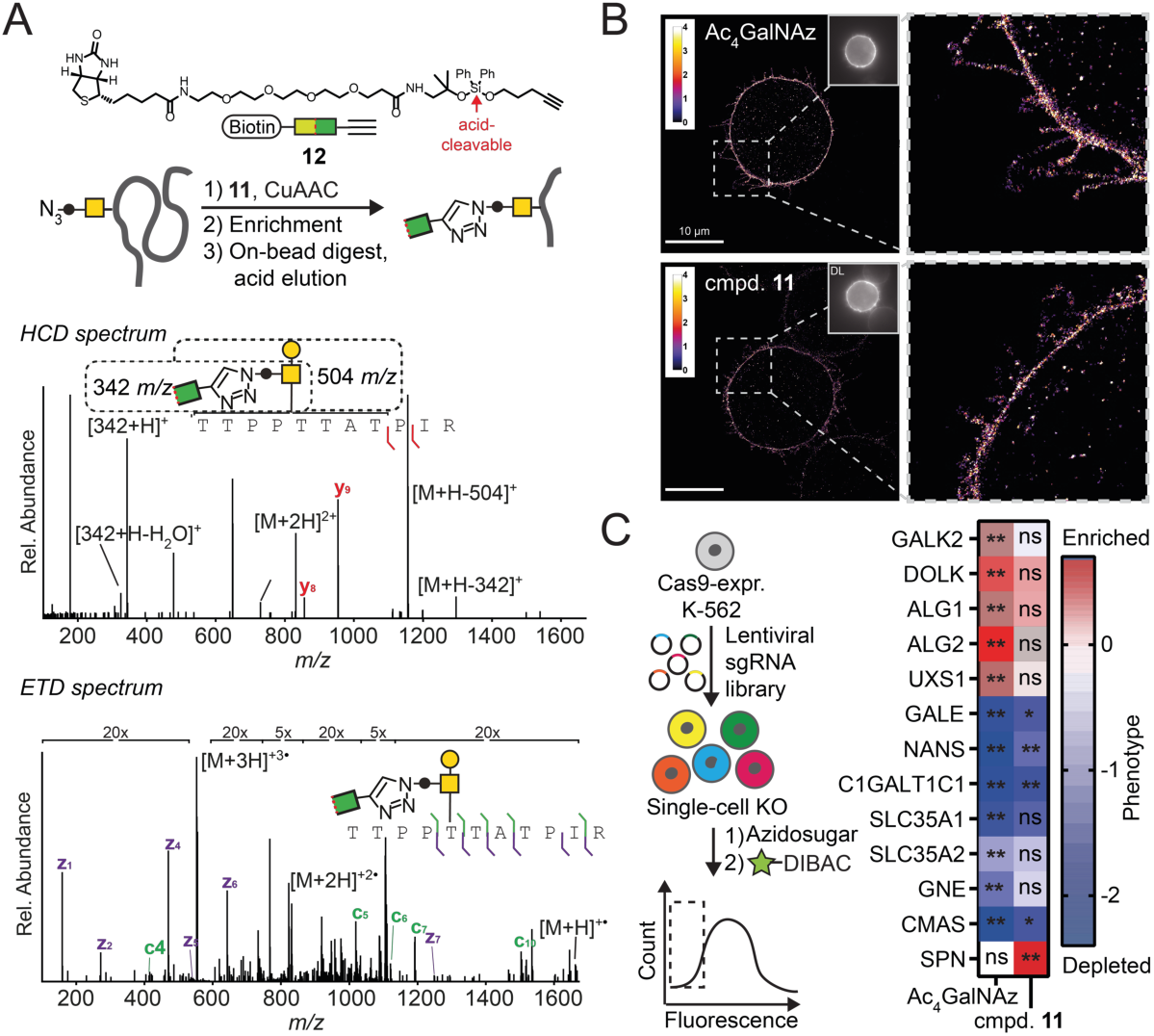
GalNAzMe is a reporter for the biology of O-GalNAc glycosylation. *A*, GalNAzMe as a reporter in mass-spectrometry based glycoproteomics of the HepG2 secretome. Exemplary mass spectra from GalNAzMe-containing glycopeptides. *B*, GalNAzMe as a reporter for superresolution microscopy using K562 cells for labeling with GalNAzMe or GalNAz and CuAAC with Alexa Fluor 647 alkyne as a visualization strategy. Scale bar, 10 µm. *C*, GalNAzMe as a reporter for a genome-wide CRISPR-KO screen in K-562 cells stably transduced with Cas9 and mut-AGX1 followed by feeding with Ac_4_GalNAz or compound **11**, labeled by MB488-DIBAC and subjected to FACS to sort the bottom 15% fluorescent cells and sequence sgRNAs. Effects on selected glyco-genes are shown – color depicts the relative phenotype (positive/red: enriched in the low fluorescence population; negative/blue: depleted in the low fluorescence bottom population), while asterisks depict false-discovery rate (FDR) as a measure of statistical significance from two independent experiments: ** FDR 2%; * FDR 5%; n.s. non-significant.

We then probed the potential of GalNAzMe as an O-GalNAc specific reporter molecule in methods of modern glycobiology. Superresolution microscopy was used to image the glycocalyx on mut-AGX1 transfected K-562 cells fed with caged GalNAzMe-1-phosphate **11** and Ac_4_GalNAz (Fig 4B). Recently-described mucin-covered tubules on these cells were clearly visible with both reagents, reflecting the fact that mucins are the most abundant glycoproteins in this cell line (42).

We next employed GalNAzMe-1-phosphate **11** as a reporter in a fluorescence-based genome-wide CRISPR-KO screen to investigate the genetic factors of glycan biosynthesis (Fig. 4C, Data S2, Data S3). Specifically, we hypothesized that GalNAzMe labeling would be sensitive to KO of genes that mediate cell surface O-glycan presentation, such as mucins. GalNAz labeling, conversely, is likely to be reduced by KO of a wider array of glyco-genes. We thus conducted paired genome-wide KO screens to reveal, in an unbiased manner, the key genes that are essential for cell-surface incorporation of the two metabolic labels. K-562 cells stably expressing *Streptococcus pyogenes* Cas9 and mut-AGX1 were transduced with a lentiviral plasmid library encoding 212821 single guide RNAs (sgRNAs) targeting 20549 genes (10 sgRNAs/gene) (43). Cells were subsequently fed with caged GalNAzMe-1-phosphate **11** or Ac_4_GalNAz and treated with the fluorophore MB488-DIBAC under SPAAC conditions. Cells with the 15% lowest fluorescence intensity were collected via FACS. Changes in sgRNA frequency were determined by deep sequencing and calculated relative to a non-treated control sample. Using the multiplicity of sgRNAs targeting the same gene, a statistical score and effect size could be derived for each gene using the casTLE scoring system (44). The gene encoding for the GalNAc 1-kinase GALK2 was essential for labeling with Ac_4_GalNAz, but not significant for labeling with caged GalNAzMe-1-phosphate **11** (Fig. 4C, Fig. S6D). This finding is consistent with the use of caged sugar-1-phosphates, such as **11**, to bypass the GALK2 step (25, 34). Strikingly, targeting the genes encoding for dolichol kinase DOLK and the mannosyltransferases ALG1 and ALG2 in the N-glycan biosynthesis pathway was detrimental for Ac_4_GalNAz labeling. In contrast, the same genes were not essential for labeling with caged GalNAzMe-1-phosphate **11**, consistent with our findings that GalNAzMe does not label N-glycans. KO of UDP-glucuronic acid decarboxylase (UXS1), an early enzyme in the biosynthesis of glycosaminoglycans such as heparin sulfate (HS), was also detrimental for GalNAz, but not GalNAzMe labeling (Fig. 4C, Fig. S6D). UDP-GalNAz **1** may enter HS after epimerization to UDP-GlcNAz that can be used as a substrate by the HS polymerases EXT1/EXT2. (45). Conversely, one of the top genes associated with GalNAzMe signal was *SPN* encoding for CD43, consistent with CD43 being glycosylated with GalNAzMe (*see* Fig. 3D). CD43 KO was not detrimental for GalNAz fluorescence, indicating that other glycans, including N-glycans, may compensate for the loss of CD43 under these conditions. Loss of several genes that encode for glycan biosynthetic determinants led to a net increase of fluorescence intensity. This was indicated by a depletion of sgRNAs in the pool of 15% cells with lowest fluorescent labeling. These genes were generally associated with the elaboration of glycans that, upon loss, probably led to better accessibility of azido-sugars to the click reagents. For instance, the chaperone C1GALTC1 is implicated in elaborating O-GalNAc glycans using UDP-galactose, a metabolite which is, in turn, shuttled into the Golgi compartment by the transporter SLC35A2. KO of both *C1GALTC1*and *SLC35A2* led to de-enrichment in the low-labeling pool (Fig. 4C). Loss of GALE generally leads to a decrease of cellular UDP-GalNAc levels (25). As a consequence, azide-tagged UDP-GalNAc analogs might be preferentially used as substrates by GalNAc-Ts, explaining the comcomitant increase in fluorescence labeling (25). Furthermore, impaired sialic acid biosynthesis by KO of the transporter SLC35A1 or the enzymes NANS, GNE, and CMAS led to an increase of labeling with both **11** and Ac_4_GalNAz. This finding is in line with our result that sialidase treatment of the cell surface increased the labeling intensity of a clickable fluorophore (*see* Fig. S3C). Taken together, these results validate GalNAzMe as a potent reporter tool for further genetic profiling of O-GalNAc glycan biosynthesis.

### Bump-and-hole mediated increase of GalNAzMe labeling by GalNAc-T2

Although UDP-GalNAzMe **5** can be biosynthesized by mut-AGX1 and enter O-GalNAc glycans, we consistently observed moderate glycoprotein labeling efficiency compared to UDP-GalNAz **2**. While it is not surprising that increasing specificity of a reagent impairs its efficiency, we tested whether GalNAzMe signal could be enhanced by a chemical genetics approach. One of the factors hampering signal was low acceptance by WT-GalNAc-Ts (*see* Fig. S2A). We therefore opted to develop a programmable labeling boost by making use of our bump-and-hole GalNAc-T technology (24, 25). We employed the GalNAc-T2^I253A/L310A^ double mutant (BH-T2) that exhibits a twofold increased activity with UDP-GalNAzMe **5** compared to the WT enzyme, but displays lower activity with UDP-GalNAc **1** and UDP-GalNAz **2** (Fig. 5A) (24, 25). Labeling of membrane proteins with UDP-GalNAzMe **5** by WT-T2 *in vitro* was competed out by increasing concentrations of UDP-GalNAc **1** (Fig. 5A). In contrast, labeling with **5** by BH-T2 could not be competed out with UDP-GalNAc **1**. Labeling with UDP-GalNAz **2** was competed out by an excess of UDP-GalNAc **1** in the presence of both WT- and BH-T2. The presence of BH-T2 also led to a marked increase of glycoprotein labeling with caged GalNAzMe-1-phosphate **11** compared to WT-T2 in the living cell, as observed by in-gel fluorescence experiments (Fig. 5B). In contrast, Ac_4_GalNAz labeling was unchanged. These data indicate that O-GalNAc labeling by GalNAzMe can be enhanced by bump-and-hole engineered BH-T2.

**Fig. 5:**
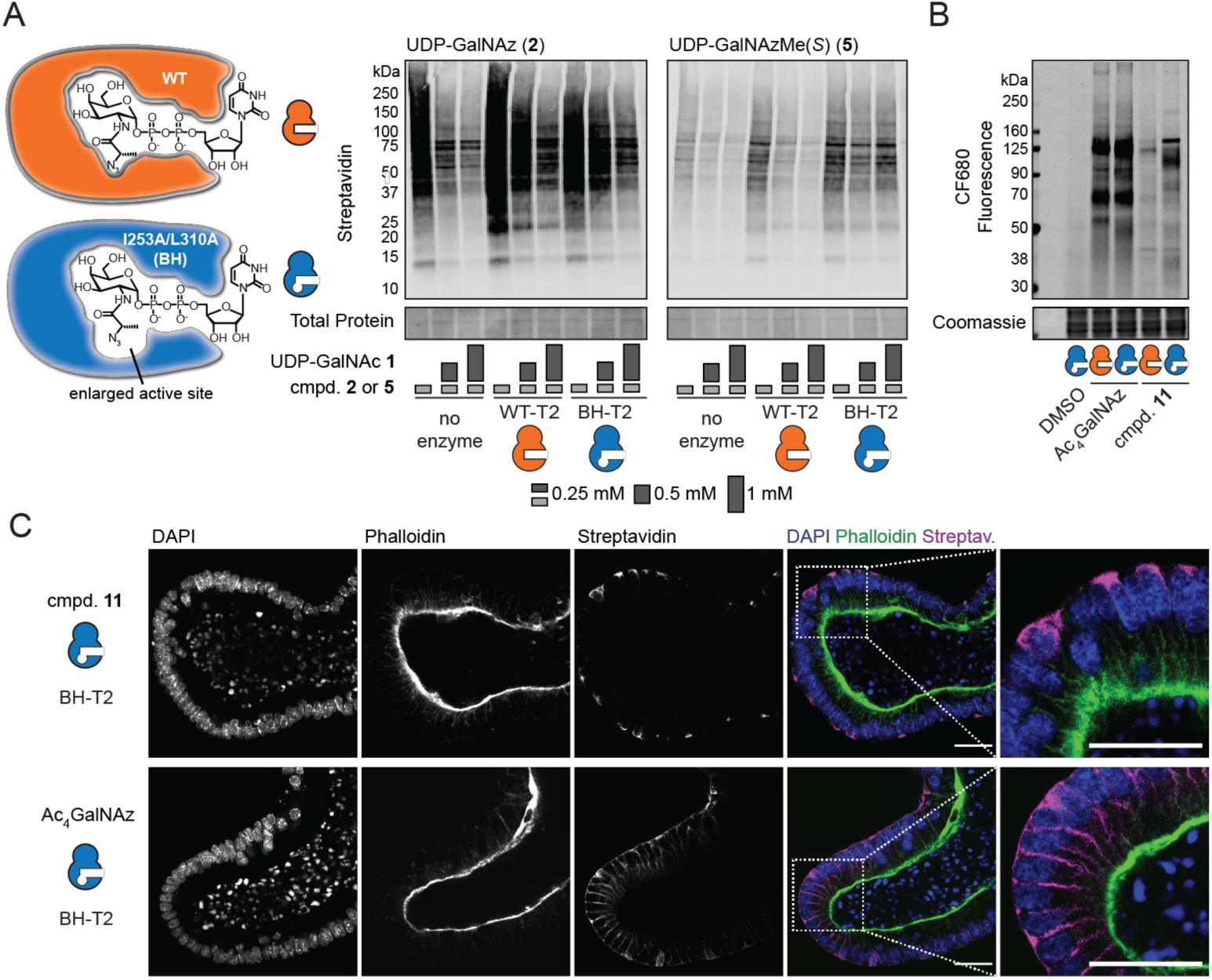
An engineered BH-T2 double mutant enhances GalNAzMe labeling. *A, in vitro* glycosylation using WT- or BH-T2 as enzyme sources. UDP-GalNAz **2** and UDP-GalNAzMe **5** were used as substrates, and UDP-GalNAc **1** was used as a competitor at different concentrations. Azide-labeled glycoproteins were visualized as in Fig. 2B. Data are from one representative out of two independent replicates. *B*, live cell surface glycosylation by K-562 cells stably transfected with mut-AGX1 and WT- or BH-T2 and fed with DMSO, 50 µM compound **11**, or 3 µM Ac_4_GalNAz. Data are from one representative out of two independent replicates. *C*, Glycosylation in intestinal organoids transfected with mut-AGX1 and BH-T2. Organoids were fed with 50 µM compound **11** or 1.5 µM Ac_4_GalNAz, fixed and treated with biotin alkyne under CuAAC conditions followed by Streptavidin Alexa Fluor 647 staining.

Data are from one representative out of two independent experiments and shown as grayscale images for each channel and a color merge image of all three channels. Scale bar, 100 µm.

### Labeling the O-GalNAc glycome in organoids

We then turned to investigating O-GalNAc glycosylation in a multicellular model system. Intestinal organoids are instrumental in understanding some of the key concepts of bowel cancer formation as well as normal gut development and homeostasis (46–50). Production of O-GalNAc glycans in such systems is often probed by either backbone-directed antibodies or lectins (51, 52). We used GalNAzMe as an O-GalNAc glycan detection tool that is independent of both protein backbone and glycan capping but reports on the peptide-proximal, invariant GalNAc moiety. We stably transfected murine intestinal organoids with both mut-AGX1 and BH-T2 and fed either caged GalNAzMe-1-phosphate **11** or Ac_4_GalNAz (53). Treatment with a clickable biotin-alkyne under CuAAC conditions and fluorescently-labeled streptavidin indicated a striking difference in labeling patterns between the two azidosugars by confocal microscopy (Fig. 5C). Ac_4_GalNAz labeling was generally found on all cell surfaces, including intercellular boundaries. In contrast, caged GalNAzMe-1-phosphate **11** labeling was focused on a subset of cells. Our labeling strategy was topologically restricted to the basolateral (non-luminal) side of the organoids, and GalNAzMe labeling was broadly localized to both cell surface and a subcortical space. Streptavidin signal was absent in both non-transfected, **11**-fed organoids as well as transfected, DMSO-fed organoids, excluding non-specific labeling (Fig. S7). We concluded that caged GalNAzMe-1-phosphate **11** is a valuable labeling tool with an O-GalNAc glycan precision that is not seen in the conventional reagent Ac_4_GalNAz.

## Conclusion

Efforts to map the systems biology of organisms, tissues, and single cells demand specific and curated reporter tools. The capacity to accurately report on the presence and dynamics of individual glycan types is essential to understanding how glycans impact biological processes. Protein-based reporter reagents have enabled the study of glycobiology, but rarely probe non-accessible glycan core structures. Thus far, the forays made into developing chemical tools have yielded an arsenal of monosaccharide analogs, for instance of ManNAc/Sia (7, 54–56), GlcNAc (17–19, 34), Fuc (57), Gal (58, 59), and GalNAc/GlcNAc (9, 14, 16, 60, 61). Probes are typically selected based on their labeling intensity, which, in turn, is often a function of poor glycan specificity. The usefulness of these probes in biological applications is therefore limited, especially in the case of GalNAc analogs that can be epimerized to the corresponding UDP-GlcNAc analogs. UDP-GlcNAc is not only thermodynamically more stable than UDP-GalNAc, but also used by a much more diverse set of glycosyltransferases (http://www.cazy.org) (14). The possibility to interconvert derivatives of both metabolites is therefore likely to create a GlcNAc-dependent labeling background if GalNAc is actually to be studied. Here, a panel of synthetic UDP-GalNAc analogs was essential to corroborate our structure-based design of the first GalNAc-specific metabolic labeling reagent. GalNAzMe is a useful monosaccharide in a range of biological applications, showcased here by superresolution microscopy, chemical glycoproteomics, a genome-wide CRISPR KO screen and imaging of intestinal organoids. Our finding that GalNAzMe incorporation can be elevated by simply expressing a bump-and-hole engineered GalNAc-T double mutant renders GalNAzMe a valuable tool even in experiments in which high glycan labeling intensity is desired. GalNAzMe is a precision tool that will prove to be invaluable in tackling important mucin-specific biological questions.

## Acknowledgements

The authors thank Douglas Fox for help with HPAEC-PAD experiments and Kayvon Pedram for providing StcE. We thank David Spiciarich and Yi-Chang Liu for helpful discussions. We thank Phil Walker for advice on vector choice, and Acely Garza-Garcia for helpful discussions on HPLC. The authors are grateful for support by the Francis Crick Institute Cell Services Science Technology Platform. The authors are grateful for generous funding by Stanford University, Stanford ChEM-H, University of California, Berkeley, and Howard Hughes Medical Institute. This work was supported by the National Institutes of Health (R01 CA200423 to C. R. B.) and the National Institute of General Medical Sciences (R35 GM118067 to W. E. M.). This work was supported by the Francis Crick Institute (to O. Y. T., N. A. G. B. T., A. C., Z. L., D. C. B., W. M. B., V. S. W. L. and B. S.) which receives its core funding from Cancer Research UK (FC001749, FC001105, FC001115), the UK Medical Research Council (FC001749, FC001105, FC001115,), and the Wellcome Trust (FC001749, FC001105, FC001115), and the the Biotechnology and Biological Sciences Research Council (BB/F008309/1 to S. M. H.). W. M. B was supported by a PhD studentship funded by the EPSRC Centre for Doctoral Training in Chemical Biology - Innovation for the Life Sciences EP/S023518/1 and GlaxoSmithKline (GSK). M. F. D. was supported by a NWO Rubicon Postdoctoral Fellowship. S. P. W. was supported by a Banting Postdoctoral Fellowship from the Canadian Institutes of Health Research. S.A.M. was supported by a National Institute of General Medical Sciences F32 Postdoctoral Fellowship (F32-GM126663-01). A. J. A. was supported by a Stanford ChEM-H undergraduate scholarship. H. L. D. acknowledges funds from Wellcome Trust New Investigator Award 104785/B/14/Z.

---

**Fig. S1:**
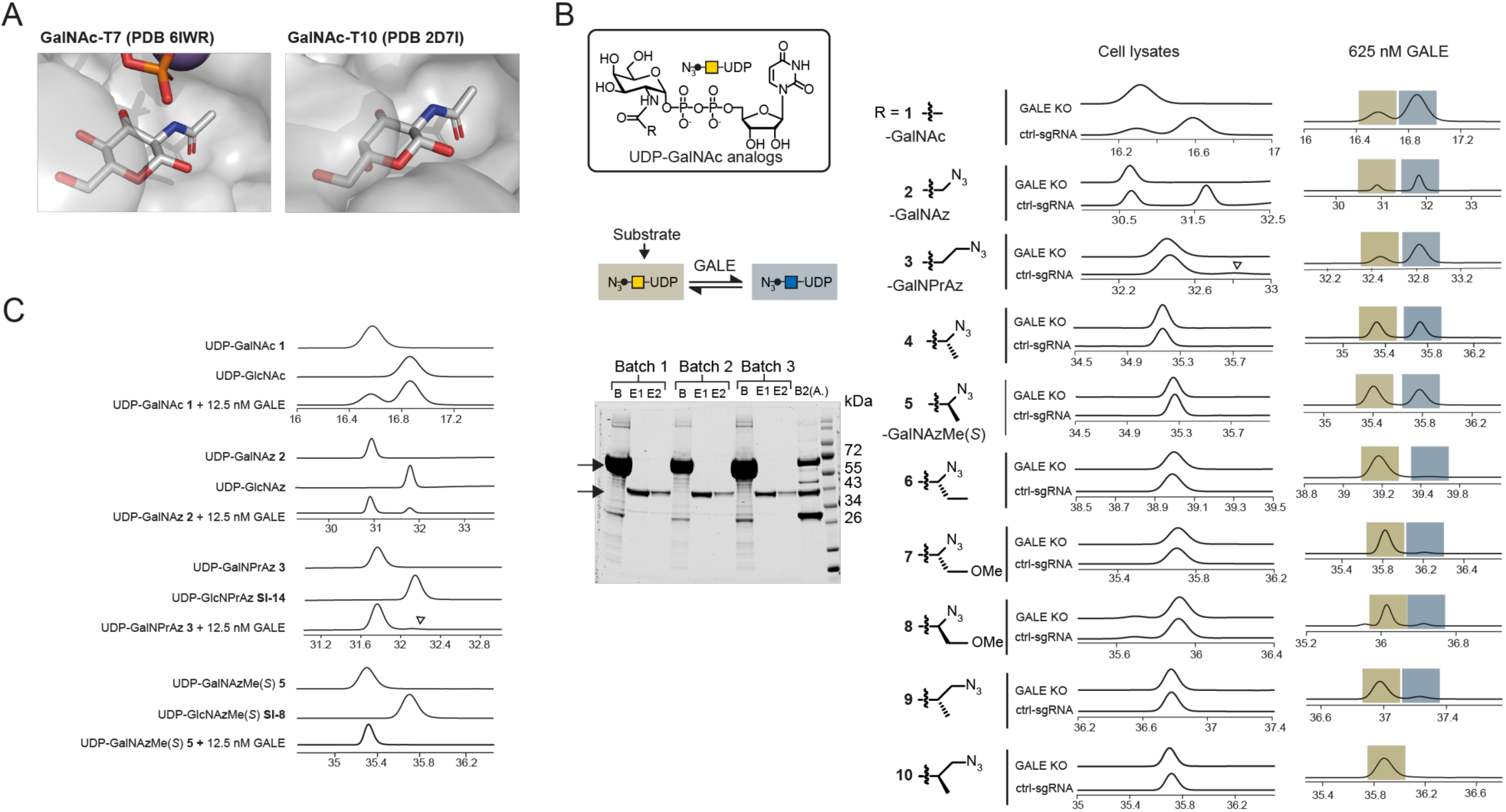
Determinants of UDP-GalNAc analog acceptance by GalNAc-Ts and GALE. *A*, Modeling of GalNAc binding by GalNAc-T7 and T10. *B*, Ion pairs HPLC traces of *in vitro* glycosylation assays, using UDP-GalNAc analogs as substrates and either cell lysates from control or GALE-KO K-562 cells or a high concentration (625 nM) of purified GALE, shown in insert (arrows pointing to GALE before and after elution), as enzyme sources. Data are representative of three independent replicates (lysate samples) or from one experiment (625 nM GALE samples). *C*, selected traces from Fig. 1B using 12.5 nM GALE as an enzyme source, with reference HPLC traces of synthetic standards for UDP-GalNAc and UDP-GlcNAc analogs. Arrowhead depicts epimerization of compound **3**. Traces depict relative intensity of absorbance at 260 nm.

**Fig. S2:**
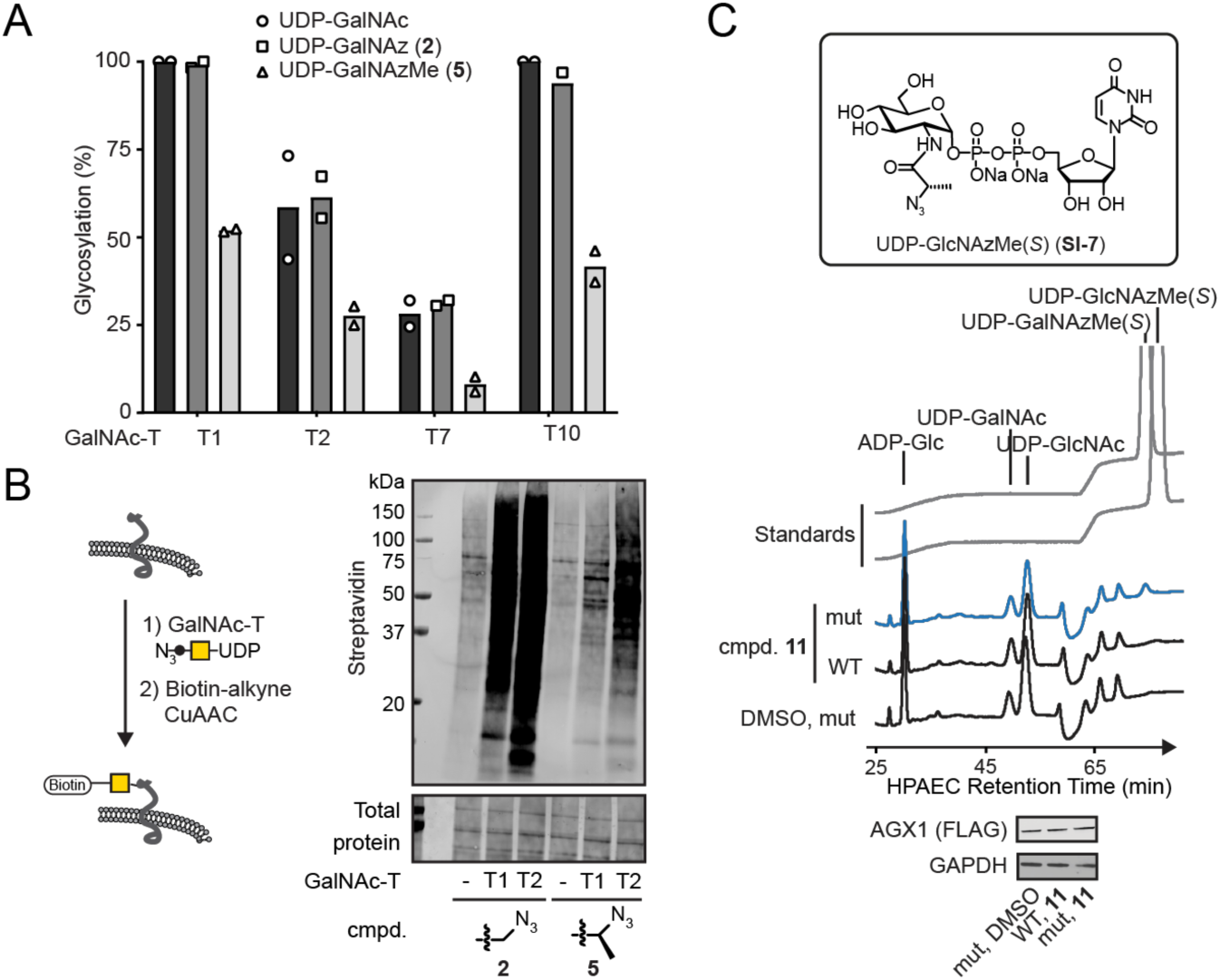
UDP-GalNAzMe 5 recognition by GalNAc-Ts and delivery to the living cell. *A, in vitro* peptide glycosylation by purified GalNAc-Ts. Data are biological duplicates as average of technical duplicates. *B*, lysate protein glycosylation by GalNAc-T1 and GalNAc-T2. A membrane preparation was used as a lysate protein source, probed with soluble GalNAc-Ts and azide-tagged UDP-sugars, and subjected to CuAAC with clickable biotin. Streptavidin blot was used to visualize glycosylation. Data are from one representative out of three independent experiments. *C*, Biosynthesis of UDP-GalNAzMe in K-562 cells stably transfected with WT- or mut-AGX1, as assessed by HPAEC-PAD. Standards include UDP-GalNAzMe (**5**) and its C4-epimer UDP-GlcNAzMe (**SI-7**).

**Fig. S3:**
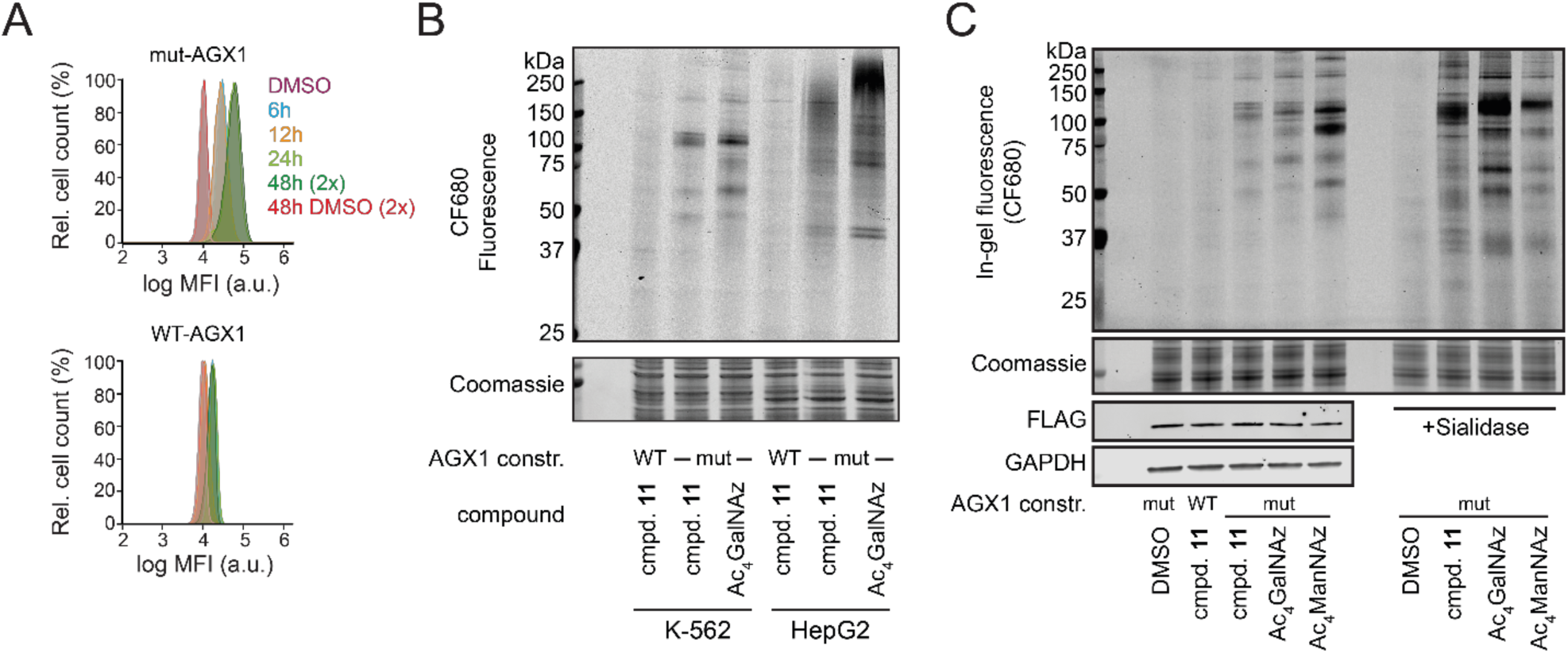
Live cell labeling using caged GalNAzMe-1-phosphate 11. *A*, time-course of cell surface labeling with 100 µM compound **11**, as assessed by flow cytometry. “2x” denominates feeding for a total of 48 h, with feeding at 0 h and 24 h. *B*, comparison of labeling by K-562 and HepG2 cells stably expressing WT-AGX1 or mut-AGX1, as assessed by in-gel fluorescence. Data are from at least five (K-562) or one (HepG2) independent experiments. *C*, cell surface glycoprotein labeling by GalNAzMe, GalNAz, and ManNAz. K-562 cells stably expressing WT-AGX1 or mut-AGX1 were fed with DMSO, 3 µM Ac_4_GalNAz, 100 µM compound **11**, or 1.5 µM Ac_4_ManNAz and treated with CF680-alkyne under CuAAC conditions. Cells were optionally treated with 10 nM *Vibrio cholerae* sialidase before the click reaction. Data are from one experiment.

**Fig. S4:**
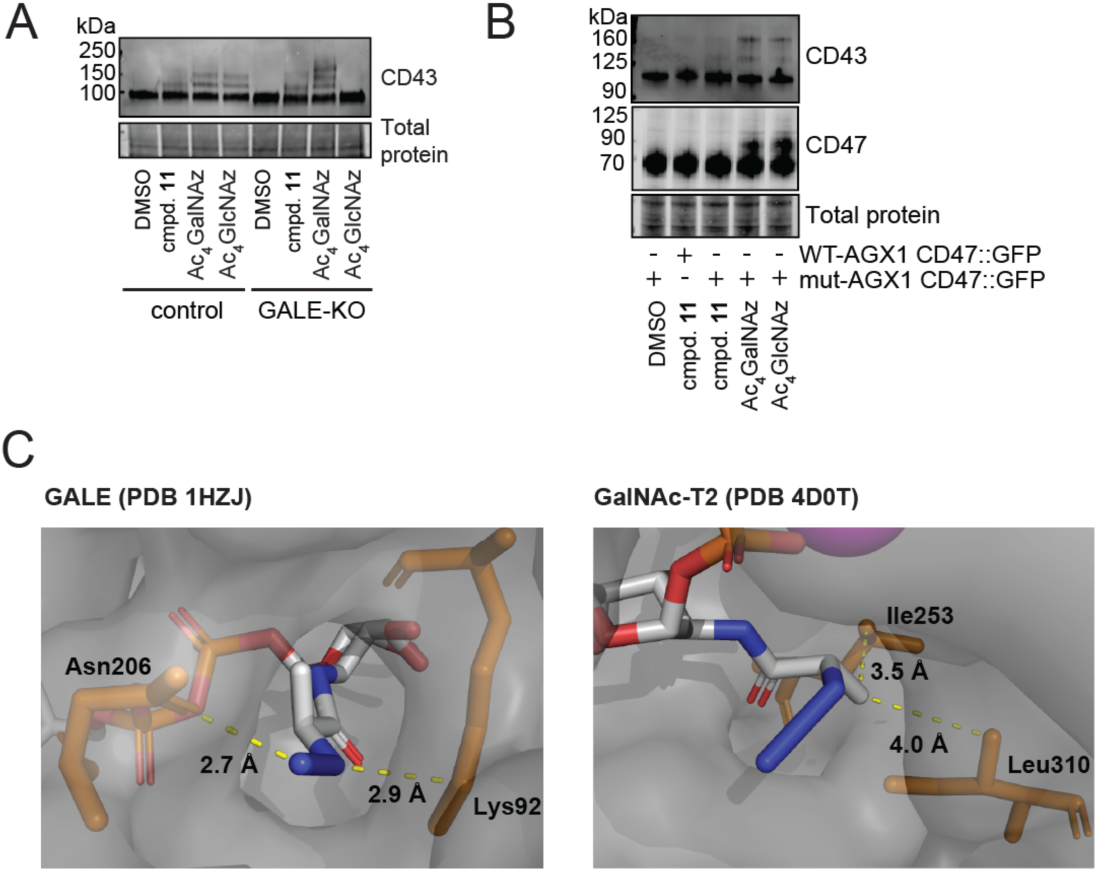
Specific O-GalNAc labeling by GalNAzMe. *A*, K-562 GALE-KO or control cells were treated with DMSO, 100 µM compound **11**, 3 µM Ac_4_GalNAz, or 8 µM Ac_4_GlcNAz, and cell lysates were treated with a clickable 10 kDa PEG mass tag under SPAAC conditions. *B*, K-562 cells stably expressing WT- or mut-AGX1 GFP::CD47 were fed and subjected to PEG mass tagging as in *A* (replicate of Fig. 3D). Samples from the same SPAAC reaction were run side by side for detection of CD43 and CD47. *C*, docking of UDP-GalNAzMe into the active sites of GALE and GalNAc-T2. Distances shown are the closest interactions to amino acid residues in the active site after energy minimization.

**Fig. S5:**
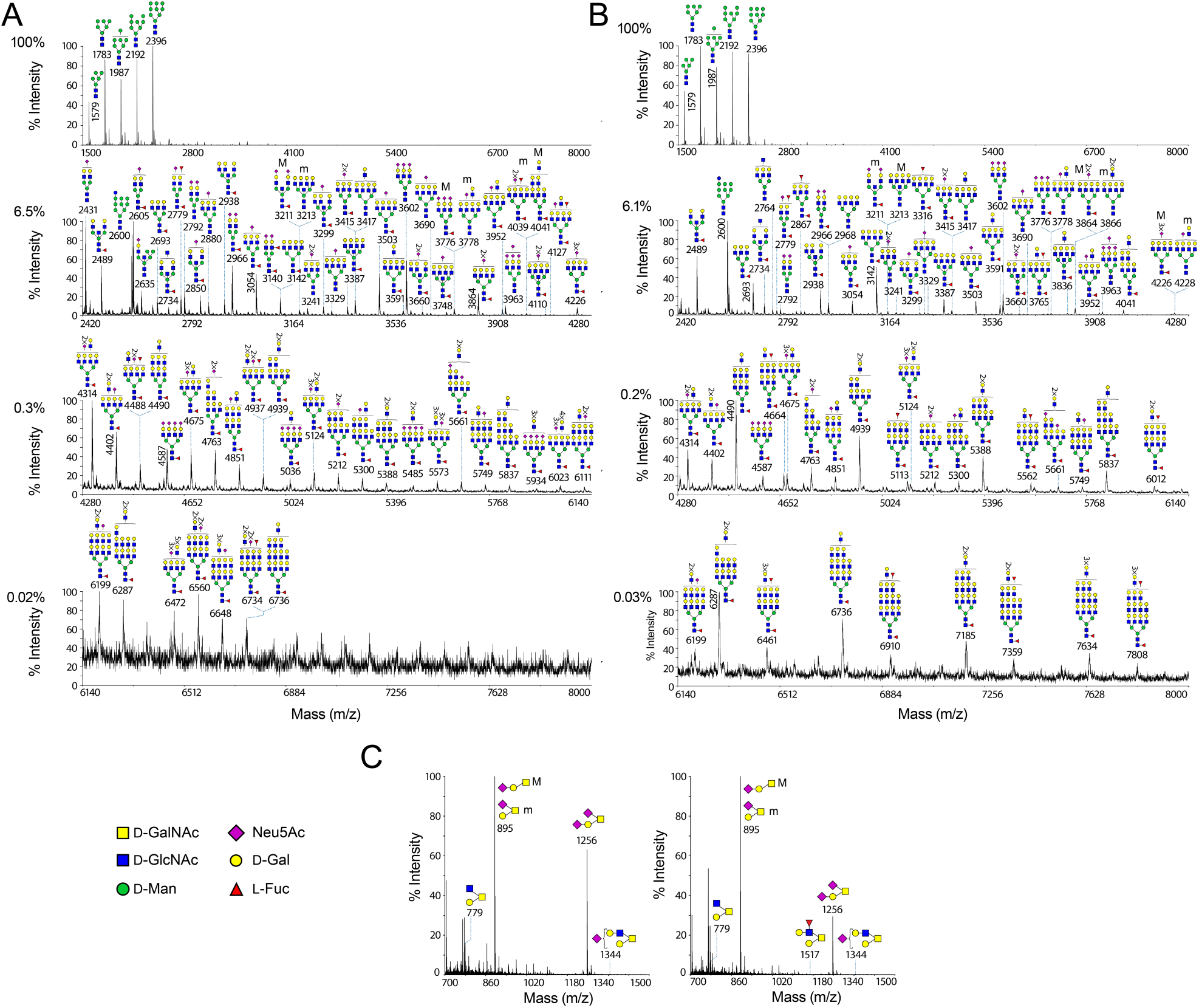
The glycomes of cells treated with Ac_4_GalNAz and compound 11 are not substantially altered. *A, B*, MALDI-TOF mass spectra of permethylated N-glycans from lysates of K-562 cells treated with 3 µM Ac_4_GalNAz (*A*) or 100 µM compound **11** (*B*). *C*, O-glycomes from the same lysates as in *A* and *B* from cells treated with Ac_4_GalNAz (left panel) and compound **11** (right panel). Structures outside a bracket have not been unequivocally defined. “M” and “m” designations indicate major and minor abundances, respectively. Top panels in (*A, B*) depict the full spectra (*m/z* 1500-8000), while lower panels *(C)* depict partial MALDI-TOF MS spectra of the corresponding areas. Percentages in (*A, B*) on the left of each partial MALDI-TOF MS panel correspond to the relative intensity of the corresponding panel relative to the full spectrum (top panel). Putative structures are based on molecular ion composition, tandem MS/MS, and knowledge of biosynthetic pathways. All molecular ions are [M+Na]^+^.

**Fig. S6:**
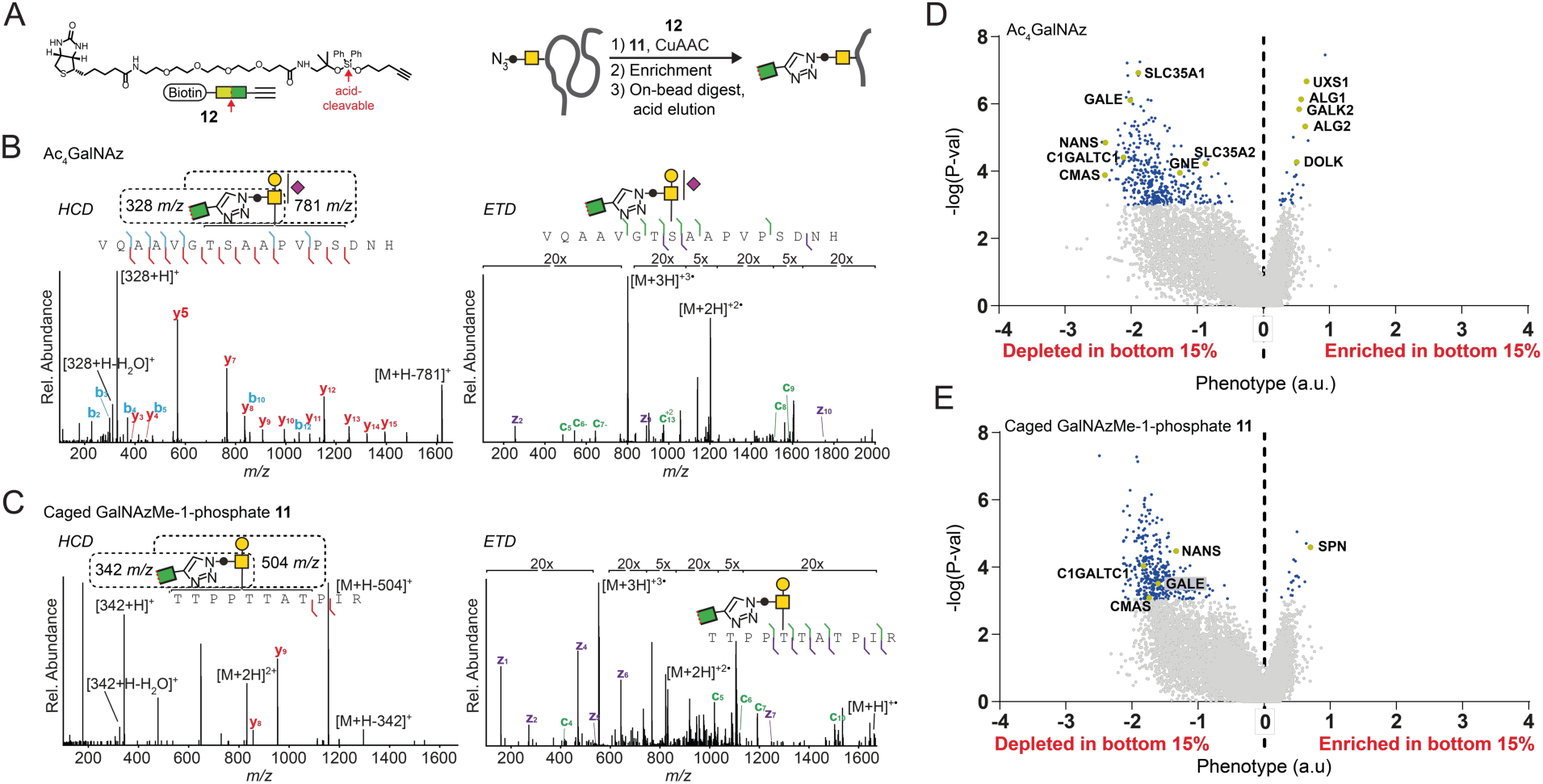
GalNAzMe as a reporter molecule in glycoproteomics and a genome-wide CRISPR KO screen. *A*, MS glycoproteomics workflow using DADPS Biotin Alkyne **12**. *B*, exemplary mass spectra from GalNAz- and *C*, GalNAzMe-containing glycopeptides. *D* and *E*, Volcano plots of a genome-wide CRISPR-KO screen of K-562 cells treated as outlined in Fig. 4C. Genes with phenotypes (5% FDR) in the respective screens are highlighted in blue, and relevant genes shown in Fig. 4C are highlighted in yellow and annotated.

**Fig. S7:**
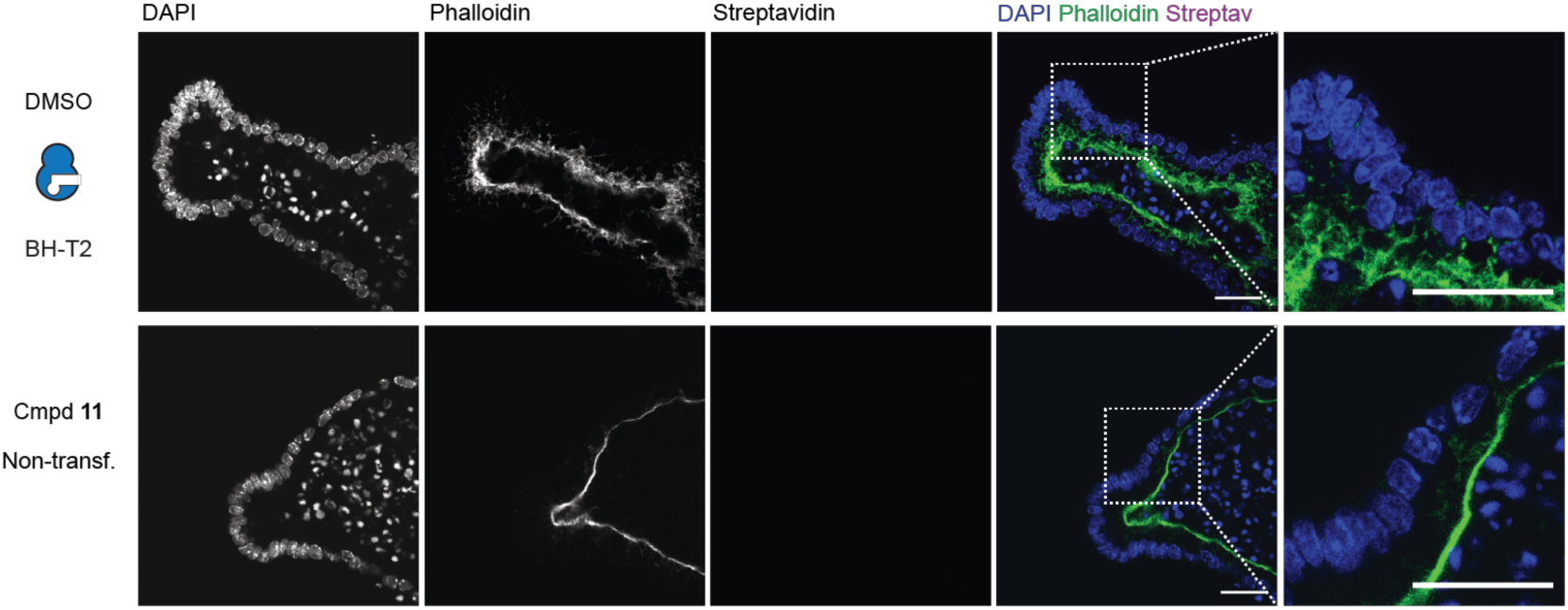
Glycosylation in intestinal organoids transfected with mut-AGX1 and BH-GalNAc-T2 or non-transfected, control samples. Organoids were fed with DMSO (top) or 50 µM compound **11** (bottom), fixed and treated with biotin alkyne under CuAAC conditions followed by Streptavidin Alexa Fluor 647 staining. Data are from one representative out of two independent experiments and shown as grayscale images for each channel and a color merge image of all three channels. Scale bar, 100 µm.

## Supporting Information

### Expression and purification of human GALE from insect cells

The coding sequence of human GALE in pDONR221 (Clone ID HsCD00040708) was from DNASU (1– 3). The coding sequence was cloned into pTriEx6-His-GST-3C-MCS (an in-house construct modified from pTriEx-6, Merck, Darmstadt, Germany) using the primers CCCTAAGCTTGGATCCAATGGCAGAGAAGGTGCTGG and GCTCGGTACCAGATCTCTAGGCTTGCGTGCCAAAG with a BamHI/BGI11 cloning strategy using the In-Fusion HD Cloning Kit (Takara, Kusatsu, Japan). Recombinant baculovirus was generated based on the *flash*BAC™ system (Oxford Expression Technologies, Oxford, UK). Sf21 cells were transfected with 0.5 µg of transfer plasmid and 100 ng of *flash*BAC™ DNA using Fugene HD (Promega, Madison, USA) according to manufacturer’s instructions. After incubation overnight, 1 mL growth media with fungizone (1:1000, Thermo Fisher, Waltham, USA) was added and cells were incubated at 125 rpm, 27 °C for 5 days. Success of transfection and infection was judged by change in cell diameter and growth. To amplify the amount of viral stock (P1 to P2); 30 mL of Sf21 cells (9×10^5^ cells/mL) were seeded at 6-well plates, incubated overnight and transferred to 30 mL Sf21 cell suspension for incubation at 125 rpm, 27 °C for 3 days. Supernatant was then collected (2000 *x g*, 5 min, 4 °C). Fetal bovine serum (FBS) was added to a concentration of 2% (v/v) to the filtered supernatant (0.22 µm filter). Viral supernatant (P2) was stored at 4 °C until required. For a final amplification of viral stock (P2 to P3); 100 mL of insect cells (9×10^5^ cells/mL) incubated overnight and 100 µL of P2 was added and cells incubated at 125 rpm, 27 °C for 3 days. Supernatant was collected, purified and stored as described above (P3). The virus MOI was determined by qPCR.

GALE was expressed first by seeding 0.5 L Sf21 cells (9×10^5^ cells/mL) and incubating at 27 °C. The following day, cells were infected with viral stocks (P3) using a MOI of 2. After incubation for 3 days, cells were harvested (2000 *x g*, 5 min, 4 °C) and stored at −80 °C. Pellets were thawed at room temperature and resuspended in 50 mL GALE Lysis Buffer (50 mM HEPES-KOH (pH 7.5), 150 mM NaCl, 1 mM EDTA, 1 mM DTT) with cOmplete protease inhibitors (Roche, Penzberg, Germany) and BaseMuncher mix (1:10,000, Expedeon, Cambridge, UK), and left at 4 °C for 1 h. Cells were then lysed by sonication using a Sonifier 450 (Branson, Hampton, USA) prior to ultra-centrifugation (108472 g, 30 min). The supernatant was collected and incubated overnight with 0.5 mL pre-equilibrated GST-4B Sepharose beads (Sigma Aldrich, St. Louis, USA) in ice-cold GALE Lysis Buffer containing 10% (v/v) glycerol. The supernatant was then collected (FT) (2000 *x g*, 3 min, 4 °C). The beads were washed twice with 10 mL GALE Lysis Buffer containing 10% (v/v) glycerol. An aliquot of 50 µL HRV 3C protease (produced in-house) and 2 mL of GALE Lysis Buffer containing 10% (v/v) glycerol was added before incubating at 4 °C for 2 h. Another 50 µL of protease were added, the incubation step was repeated and the supernatant collected (E1). The beads were further washed with 2 mL GALE Lysis Buffer containing 10% (v/v) glycerol and the supernatant collected (E2). Beads were then incubated overnight with 150 µL of HRV 3C protease and 2 mL GALE Lysis Buffer containing 10% (v/v) glycerol at 4 °C. The supernatant was collected (E3) and the beads washed twice with 2 mL GALE Lysis Buffer containing 10% (v/v) glycerol (E4, E5). E1-E5 were pooled and concentrated to 1 mL using a Vivaspin6 30K centrifugal tube (Sartorius, Göttingen, Germany). The concentrated sample was injected onto an ÄKTA™ Pure system, running a Superdex™ S75 16/60 gel filtration column (GE Life Sciences, Marlborough, USA), collecting 1 mL fractions in GALE Lysis Buffer containing 10% (v/v) glycerol. Fractions were pooled, aliquoted (15.15 µM concentration) and stored at - 80 °C.

### In vitro epimerization

The protocol for *in vitro* epimerization was based on Kingsley et al. (4). K-562 GALE-KO and the corresponding control cells carrying a non-targeting sgRNA were prepared previously (5) and grown using RPMI with 10% (v/v) FBS, penicillin (100 U/mL), streptomycin (100 µg/mL), 20 µM galactose and 200 µM *N*-acetylgalactosamine. Five million cells were harvested (500 *x g*, 5 min, 4 °C), washed with PBS once, harvested and frozen at −80°C. Cells were treated with 250 µL 100 mM glycine-HCl, pH 8.7, and subjected to three freeze/thaw cycles between dry ice and room temperature (5 min each). Cell debris was harvested (12000 *x g*, 15 min, 4 °C), the supernatant was transferred to a fresh tube and treated with 35 µL 80% (v/v) glycerol. Samples were aliquoted and stored at −80 °C. The protein concentration was between 6 µg/µL and 9 µg/µL.

*In vitro* epimerization reactions were run in 25 µL reactions, containing either cell lysates (12 µg protein) or purified GALE (12.5 or 625 nM) in 25 mM Glycine-HCl (pH 8.7), 200 µM NAD and 250 µM UDP-GalNAc analog. Reactions run with cell lysates additionally contained 5 mM sodium pyruvate. Reactions were run for 30 min (with purified GALE) or overnight (with cell lysates) at 37 °C, diluted with 75 µL water and cooled to 4 °C. Samples were run on a 1260 HPLC with diode array detector using a Poroshell 120, EC-C18, 2.7 µm, 3.0 x 150 mm column (Agilent, Santa Clara, USA). Solvents were: A = 100 mM potassium phosphate, pH 6.4, 8 mM tetrabutylammonium bisulfate; B = 80% A, 20% acetonitrile. Gradients were either 0 min 0% B; 30 min 40% B; 32 min 100% B; 34 min 0% B; 44 min 0% B, or 0 min 0% B; 19 min 25% B; 32 min 100% B; 34 min 100% B; 35 min 0% B; 46 min 0% B.

Samples were also run on an ICS-6000 with a quaternary pump and pulsed amperometric detection (Thermo Fisher) on a CarboPac PA1 4×250 mm column and a 4×50 mm guard column. Solvents were: A = 1 mM NaOH in degassed water; B = 1 mM NaOH, 1M NaOAc in degassed water. 0 min 60% B; 40 min 100% B; 45 min 100% B; 60 min 100% B.

Commercial or synthetic standards (200-500 µM) were used as controls.

### Peptide glycosylation

HPLC was performed on a 1100 series HPLC system (Agilent). LC-MS experiments were carried out using a 1260 Infinity HPLC attached to a 6120 Quadrupole mass spectrometer (Agilent). Poroshell 120 EC-C18, 2.7 µm, 4.6 × 50 mm analytical LC columns (Agilent) were used for both HPLC and LC-MS.

In vitro glycosylation was performed according to our published protocol. Soluble GalNAcT-1, T-2 and T-10 enzymes were expressed according to a published procedure (6). Soluble GalNAcT-7 was expressed as a fusion construct with superfolder GFP in pGEn2-DEST (a kind gift from Kelley Moremen, University of Goergia, Athens, USA) in HEK293-F cells according to the manufacturer’s instructions (Thermo) (7). Briefly, a 30 mL culture was transfected with 293Fectin (Thermo) according to the manufacturer’s instructions. After 48 h, cells were harvested (500 g, 5 min, 4 °C) and supernatant was kept for protein isolation. The supernatant was centrifuged at (9000 g, 20 min, 4 °C), and a cOmplete protease inhibitor tablet (Roche, Basel, Switzerland) was added. Ni-NTA agarose (1 mL settled resin, Thermo) were washed with water and Phosphate Buffered Saline without Ca^2+^ or Mg^2+^ (PBS) containing 10 mM imidazole (pH 7.4), and added to the protein solution. The suspension was incubated for 1 h at 4 °C under rotation and poured into an empty polystyrene column (Bio-Rad, Hercules, USA). The resin was washed with 50 mL PBS containing 20 mM imidazole (pH 7.4), and elution was carried out with 10 mL PBS containing 250 mM imidazole (pH 7.4). GalNAcT-7 was concentrated with by centrifuge filtration (10 kDa MWCO, Millipore, Burlington, USA), washed with 25 mM Tris-Cl (pH 7.4), 150 mM NaCl and concentrated to approx. 1500 nM enzyme. Glycerol was added to a final concentration of 20% (v/v), and enzyme was frozen at −80 °C. Typically, 100-200 µg enzyme were obtained from a 30 mL culture.

For T-2, T-7 and T-10, chromophore-containing, isoenzyme-optimized peptide substrates were used with an HPLC-based assay to assess conversion (for peptide structures, see Table S1 and Choi, Wagner et al) (6). For T-1, EA2 peptide (Anaspec, Fremont, USA) was used as substrate with an LCMS-based assay to assess conversion, as azide-containing glycopeptides were not separable from the corresponding peptide substrate when a previously reported T-1-optimized peptide was used (6).

All reaction mixtures contained 20.8 mM Tris-HCl (pH 7.4), 50 mM NaCl, 10 mM MnCl_2_, 12.5% glycerol, GalNAcT enzymes (see Table S2), 250 µM UDP-sugar and 50 µM peptide substrate in 50 µL final volume. Reactions were carried out for the time indicated in Table X, quenched with 150 mM EDTA pH 8.0 (25 µL) and analyzed by HPLC or LCMS (Table Y) (6).

**Table S1.** Peptide glycosylation conditions. DAA = 2,4-dinitrophenyl-5-L-alanine amide (DAA); T* = α-D-GalNAc-*O*-Thr

| GalNAcT isoenzyme | Enzyme concentration | Peptide substrate | Reaction time | Analysis method |
| --- | --- | --- | --- | --- |
| T-1 | 80 nM | EA2 | 2 h | LCMS |
| T-2 | 25 nM | (DAA)GAGAPGTPGPAGAGK | 1 h | HPLC |
| T-7 | 50 nM | (DAA)GTT*PSPVPTTSTTSAP | 1 h | HPLC |
| T-10 | 60 nM | (DAA)GTT*PSPVPTTSTTSAP | 1 h | HPLC |

HPLC and LCMS conditions are provided in Table Y.

**Table S2.** HPLC methods used for to assess turnover in glycosylation reactions. Conditions depict the gradient as a percentage of acetonitrile in water with 0.1% (v/v) formic acid (FA). MS = mass spectrometry; UV = detection by absorption at 340 nm.

| Isoenzyme/substrate combination | Detection | Conditions |
| --- | --- | --- |
| T-1/all UDP-sugars | MS | 0 min 5%; 2 min 5%; 17 min 85%; 18 min 100%; 23 min 100%; 24 min 5%; 25 min 5% |
| T-2/UDP-GalNAc | UV | 0 min 21.5%; 35 min 21.5%; 36 min 100%; 40 min 100%; 41 min 21.5%; 45 min 21.5% |
| T-2/UDP-GalNAzMe | UV | 0 min 21.5%; 30 min 21.5%; 31 min 100%; 35 min 100%; 36 min 21.5%; 40 min 21.5% |
| T-2/UDP-GalNAz | UV | 0 min 20%; 40 min 20%; 41 min 100%; 45 min 100%; 46 min 20%; 50 min 20% |
| T-2/UDP-GalNPrAz | UV | 0 min 20%; 40 min 20%; 41 min 100%; 45 min 100%; 46 min 20%; 50 min 20% |
| T-7/all UDP-sugars; T-10/all UDP-sugars | UV | 0 min 22.5%; 25 min 22.5%; 26 min 100%; 30 min 100%; 31 min 22.5%; 35 min 22.5% |

For reactions containing T-1, extracted ion chromatograms (EIC) were generated of singly and doubly glycosylated species, integrated and related to the intensity of unglycosylated peptide. The m/z values [M+2H]^2+^ used were 658.71 (EA2), 760.31 (EA2-GalNAc), 861.9 (EA2-2xGalNAc), 780.81 (EA2-GalNAz), 902.915 (EA2-2xGalNAz), 787.83 (EA2-GalNAzMe, EA2-GalNPrAz), 917.94 (EA2-2xGalNAzMe, EA2-2xGalNPrAz). Of note, total intensities of peptide and glycopeptide species inversely correlated with the abundance of doubly glycosylated species. As doubly glycosylated species were mainly present in glycosylations with UDP-GalNAc and UDP-GalNAz, we note that the abundance of unglycosylated peptides was likely overestimated for glycosylations using UDP-GalNAzMe and UDP-GalNPrAz.

### Plasmids

The plasmids pIRES-puro containing AGX1^WT^, AGX1^F381G^, AGX1^F383G^, AGX1^F381G/F383G^, AGX1^F381A^, AGX1^F383A^, AGX1^F381A/F383A^, pSBtet-AGX1^WT^ and pSBtet-AGX1^F383A^ were generated in our previous work (5, 8). The terminus “mut-AGX1” depicts AGX1^F383A^ throughout this manuscript. GFP_CD47_SU (65473) were used to generate GFP::CD47 K562 cell lines. GFP_CD47_SU was a gift from Christine Mayr (Addgene plasmid # 65473; http://n2t.net/addgene:65473; RRID:Addgene_65473) (9). pCMV(CAT)T7-SB100 was a gift from Zsuzsanna Izsvak (Addgene plasmid #34879; http://n2t.net/addgene:34879; RRID:Addgene_34879) (10).

### Cell transfection

Tet-system approved FBS (Takara) was used to propagate all cell lines transfected with pSBtet-based plasmids. K-562 cells were a gift from Jonathan Weissman (University of California, San Francisco). K-562 cells with stable expression of *Streptococcus pyogenes* Cas9 (K-562-spCas9) were prepared in-house. Cells were grown using RPMI with 10% (v/v) FBS, penicillin (100 U/mL) and streptomycin (100 µg/mL). Cells were transfected with pSBtet-based plasmids using Lipofectamine LTX (Thermo) according to the manufacturer’s instructions, with a 20:1 (m/m) mixture of pSBtet and <u>pCMV(CAT)T7-SB100</u> plasmid DNA. After 24 h, cells were harvested and selected in growth medium containing 150 µg/mL hygromycin B (Thermo) for 7-10 days to obtain stable cells.

HepG2 cells (ATCC HB-8065) were propagated in low-glucose DMEM (Caisson Labs, Smithfield, USA) with 10% (v/v) FBS, penicillin (100 U/mL) and streptomycin (100 µg/mL). Cells were transfected with Lipofectamine 3000 (Thermo Fisher) according to the manufacturer’s instructions, using a 20:1 (m/m) mixture of pSBtet and <u>pCMV(CAT)T7-SB100</u> plasmid DNA. After 24 h, medium was aspirated, and cells were treated with fresh growth medium containing 600 µg/mL hygromycin B for two weeks to obtain stable cells. Following selection, cells were propagated in 200 µg/mL hygromycin B in growth medium.

HEK293T (ATCC CRL-3216) were grown in DMEM (Thermo Fisher) with 10% (v/v) FBS (Thermo Fisher), penicillin (100 U/mL) and streptomycin (100 µg/mL, GE Healthcare, Chicago, USA). Cells were transfected with pIRES-puro3 plasmids containing AGX1 constructs using TransIT-293 (Mirus Bio LLC, Madison, USA) according to the manufacturer’s instructions and 37.5 µg DNA per 15 cm dish or 15 µg DNA per 10 cm dish. After 24 h, medium was aspirated, and cells were treated with fresh growth medium and compounds for analysis of nucleotide-sugar biosynthesis (see below).

### Lysate labeling

Membrane lysate labeling was performed as previously reported (5). Briefly, HepG2 cells were fractionated using the Subcellular Fractionation Kit for Cultured Cells (Thermo Fisher) according to the manufacturer’s instructions. The membrane fraction was heat-inactivated to abrogate endogenous GalNAcT activity. Glycosylation reactions were performed on 10 µg membrane protein in 20 µL reaction volume containing 62.5 mM Tris-HCl (pH 7.4), 150 mM NaCl, 10 mM MnCl_2_, 250 µM UDP-GalNAc analog and soluble GalNAcT-1 or T-2 at a final concentration of 20 nM (T-1) or 10 nM (T-2) at 37 °C for 12 h. Reactions were heat-inactivated at 95 °C for 20 s and subsequently cooled to 4 °C. Then, azide-containing reaction mixtures were sequentially treated with equal volumes (1.25 µL each) of 2 mM biotin-PEG_4_-alkyne (Thermo Fisher), 2 mM BTTAA (Click Chemistry Tools, Scottsdale, USA), 20 mM CuSO_4_ and 100 mM sodium ascorbate (final concentrations 100 µM biotin probe, 100 µM BTTAA, 1 mM CuSO_4_ and 5 mM sodium ascorbate). Reactions were performed at room temperature for 2 h and quenched with 50 mM EDTA. Protein mixtures were then subjected to SDS-PAGE and blotted on nitrocellulose membranes. The total protein amount was assessed using the REVERT protein staining kit (LI-COR Biosciences, Lincoln, USA), and biotin signal was detected using IRDye 800CW Streptavidin (LI-COR Biosciences) according to the manufacturer’s instructions.

### Modelling

The crystal structures of human GALE, GalNAcT-2 (PDB 4D0T), GalNAcT-7 (PDB 6IWR), GalNAcT-10 (PDB 2D7I) and AGX1 (PDB 1JV3) were visualized with Pymol 2.0.0 (Schrodinger LLC, New York) (11–15). Modelling of UDP-GalNAzMe into GALE and GalNAcT-2 was performed using simple minimization in COOT (16). Ligand restraints for UDP-GalNAzMe were generated using phenix.elbow (17).

### In vitro epimeriszation assay

In vitro epimeriszation was performed according to a literature precedent (4). Briefly, 5 million K-562 GALE-KO or control cells stably transfected with pSBtet-AGX1^F383A^ were harvested, washed once with PBS and frozen at −80 °C. Cells were thawed on ice and treated with the Cytosolic Extraction Buffer of the Subcellular Fractionation Kit for Cultured Cells (250 µL). Cells were lysed according to the manufacturer’s instructions. The protein content of the cytosolic extract was assessed by BCA and equal protein amounts were used for *in vitro* epimerization. Assays contained 2 µL protein extract, 100 µM NAD and 1 mM UDP-sugar in 20 µL water. Reactions were carried out for 16 h at 37 °C, quenched by the addition of 1 µL 100 µM NaOH, and diluted to 50 µL with water. Samples were analyzed by HPAEC-PAD.

### Analysis of nucleotide-sugar biosynthesis by High Performance Anion Exchange Chromatography

Cells expressing AGX1-FLAG (transient or stably transfected HEK293T in 20 mL growth medium in a 10 cm dish or 5 million K-562 or K-562 GALE-KO cells stably transfected with pSBtet-AGX1^WT^ or pSBtet-AGX1^F383A^ in 4 mL growth medium) were fed Ac_4_GalNAz or caged GalNAzMe-1-phosphate analog **11** (100 µM final concentration from a 100 mM stock solution in DMSO) or DMSO vehicle. After 7 h, cells were harvested. K-562 cells were centrifuged at 500 *x g*, 5 min, 4 °C and resuspended in PBS (1 mL). HEK293T cells were washed once on the plate with cold PBS (8 mL), scraped in cold 1 mM EDTA in PBS (8 mL), transferred to a conical tube and harvested (300 g, 5 min, 4 °C). Cell pellets were resuspended in PBS (1 mL). 0.9 mL cell suspension was transferred to O-ring tubes (1.5 mL, Thermo Fisher) and centrifuged. Zirconia/silica beads (0.1 mm, BioSpec, Bertlesville, USA) were added at a similar volume to the cell pellet, followed by 1:1 acetonitrile/water (1 mL). Cells were lysed using a bead beater (FastPrep-24, MP Biomedicals, Santa Ana, USA) at 6 m/s for 30 s, and the lysate was cooled at 4 °C for 10 min. Samples were centrifuged (14000 *x g*, 10 min, 4 °C), and the supernatant was transferred to a new tube. The solvent was evaporated by speed vac. The residue was dissolved in LCMS-grade water (Thermo Fisher, 0.2-0.4 mL) containing 15 µM ADP-α-D-glucose (Sigma Aldrich). The solution was dialyzed (30 min, 14000 *x g*) using a 3 kDa Amicon Ultra Centrifugal Filter Unit (Merck). High performance anion exchange chromatography was used to analyze lysates.

The residual cell suspension in PBS (0.1 mL) was centrifuged, and the pellet was resuspended in M-PER lysis buffer (Thermo Fisher) with cOmplete protease inhibitor (0.2 mL). The solution was incubated at room temperature for 10 min and centrifuged (14000 *x g*, 10 min, 4 °C). The supernatant was transferred into a new tube, the protein concentration was measured by BCA, and samples were used for analysis of protein expression.

High performance anion exchange chromatography was carried out using an ICS-5000 with a quaternary pump and pulsed amperometric detection (Thermo Fisher) on a CarboPac PA1 4×250 mm column and a 4×50 mm guard column. Solvents were: A = 1 mM NaOH in degassed water; B = 1 mM NaOH, 1M NaOAc in degassed water. 0 min 5% B; 20 min 40% B; 60 min 40% B; 63 min 50% B; 83 min 50% B; 87 min 100% B; 95 min 100% B; 97 min 5% B; 105 min 5% B. Commercial or synthetic standards (200-500 µM) were used as controls.

### Metabolic cell surface labelling, flow cytometry and in-gel fluorescence

K-562 cells stably transfected with pSBtet-AGX1^WT^ or pSBtet-AGX1^F383A^ were seeded into well plates at a density of 250,000 cells/mL in growth medium without hygromycin. Cells were treated with DMSO, caged GalNAc-1-phosphate analog **11**, Ac_4_GalNAz, Ac_4_ManNAz, or Ac_4_GlcNAz at the indicated concentrations and using either GalNAc or GlcNAc as additives in the indicated concentrations. Cells were grown for another 20 h.

Cells were optionally treated with enzymes before fluorescence-based readout. StcE (50 nM final concentration) was added directly to the cell suspension, while for sialidase treatment, cells were harvested, washed once with serum-free RPMI media and treated with *Vibrio cholerae* sialidase (10 nM final concentration, generated in-house) in serum-free media (18). Enzyme treatment was performed for 2 h at 37 °C.

For in-gel fluorescence, cells were harvested in a V-shaped 96 well plate and washed twice with 2% FBS in PBS (Labeling Buffer, 0.2 mL). Cells were resuspended in Labeling Buffer (35 µL), treated with a solution of 200 µM CuSO_4_, 1200 µM BTTAA (Click Chemistry Tools, Scottsdale, USA), 5 mM sodium ascorbate, 5 mM aminoguanidinium chloride and 200 µM CF680 picolyl azide in Labeling Buffer (35 µL), and incubated for 7 min at room temperature on an orbital shaker. The click reaction was quenched with 3 mM bathocuproinedisulfonic acid in PBS (35 µL). Cells were centrifuged, washed twice with Labeling Buffer and then with PBS, and treated with ice-cold Lysis Buffer (50 mM Tris-HCl pH 8, 150 mM NaCl, 1% (v/v) Triton X-100, 0.5% (v/v) sodium deoxycholate, 0.1% (w/v) SDS, 1 mM MgCl_2_, and 100 mU/µL benzonase (Merck) containing cOmplete protease inhibitors, (0.1 mL). Cells were lysed for 20 min at 4 °C on an orbital shaker and centrifuged (1500 *x g*, 20 min, 4 °C). Supernatant was transferred to a new plate and BCA was used to measure protein concentration. For enzyme treatment, equal amounts of protein (typically 15 µg) were diluted to 40 µL with Lysis Buffer or PBS, treated with either SialEXO (4 µL of a 4 U/µL solution in 50 mM Tris-HCl (pH 6.5), Genovis, Lund, Sweden) or the glycoprotease StcE (50 nM in PBS) (19) and incubated for 2 h at 37 °C. The reaction was quenched by heating to 95 °C for 10 s with subsequent cooling at 4 °C. Loading buffer (a 1:1:1:0.5 (v/v/v/v) mixture of 1 M Tris-HCl pH 6.5, 80% (v/v) glycerol, 10% (w/v) SDS and 1 M DTT) was added, samples were heated at 95 °C for 30 s, run on a 10% Criterion™ gel (Bio-Rad, Hercules, USA) for SDS-PAGE, and imaged on an Odyssey CLx imager (LI-COR Biosciences, Lincoln, USA). Total protein was stained with Coomassie using Acquastain (Bulldog Bio, Portsmouth, USA). Protein expression was assessed by Western blot with a different set of samples, using antibodies against GALE (sc-390407, Santa Cruz Biotechnology, Dallas, USA), GAPDH (ab128915, abcam) or FLAG tag (mouse anti-FLAG M2, Sigma Aldrich).

For flow cytometry, cells were harvested, washed twice with Labeling Buffer, resuspended in 100 µM DIBAC-sulfo-biotin (DBCO-sulfo-biotin, Jena Bioscience, Jena, Germany) in Labeling Buffer (100 µL) and incubated for 1 h at room temperature on an orbital shaker. Cells were washed twice, and treated with DTAF-streptavidin (1:000, Jackson ImmunoResearch, Cambridge, UK). Cells were incubated for 1 h at room temperature, washed twice and treated with SYTOX red (1:1000, Thermo) in Labeling Buffer. Flow cytometry was performed on an Accuri C6 flow cytometer (Becton Dickinson, Franklin Lakes, USA).

### Superresolution microscopy

K-562 cells stably transfected with pSBtet-AGX1^WT^ or pSBtet-AGX1^F383A^ were seeded into well plates at a density of 180,000 cells in 680 µL growth medium without hygromycin. Cells were treated with either DMSO, 100 µM compound **11** or 5 µM Ac_4_GalNAz. Cells were incubated for 16 h. An 8-chamber coverslip (Lab-Tek II 155409, Thermo) was coated with 250 µL human Fibronectin (20 µg/mL, Sigma Aldrich) for 1 h at 37 °C, and washed with PBS. Cells were transferred to one well each and further incubated for 5 h at 37 °C. Cells were moved to 4 °C, medium was aspirated, and cells were washed with ice-cold PBS (4x 300 µL). Cells were then treated first with 100 µL PBS and 100 µL of a freshly prepared solution containing 1200 µM BTTAA, 200 µM CuSO4, 5 mM sodium ascorbate, 5 mM aminoguanidinium chloride and 400 µM biotin-PEG3-alkyne (Click Chemistry Tools). The reaction was carried out for 6 min at 4 °C, the supernatant was removed and cells were washed with PBS (4x 300 µL). Cells were incubated with 20 µg/mL AF647-streptavidin (Jackson ImmunoResearch) for 30 min at 4 °C, then washed again. Cells were fixed with 4% (v/v) paraformaldehyde (Thermo) and 0.2% (v/v) glutaraldehyde (Sigma Aldrich) in PBS for 30 min at 4 °C, then washed with PBS (4×300 µL).

The instrument setup is based on an inverted microscope (IX71, Olympus, Tokyo, Japan). The laser used for illumination (120 mW 647 nm, CW, Coherent, Santa Clara, CA) was spectrally filtered (ff01-631/36-25 excitation filter, Semrock, Rochester, NY) and circularly polarized (LPVISB050-MP2 polarizers, Thorlabs, Newton, NJ, WPQ05M-633 quarter-wave plate, Thorlabs). The beam was expanded and collimated using Keplerian telescopes. Shutters were used to toggle the lasers (VS14S2T1 with VMM-D3 driver, Vincent Associates Uniblitz, Rochester, NY). The laser was introduced into the back port of the microscope via a Köhler lens. The sample was mounted onto an XYZ stage (PInano XYZ Piezo Stage and High Precision XY Microscope Stage, Physik Instrumente, Karlsruhe, Germany). Emitted light was detected using a high NA detection objective (UPLSAPO100XO, x100, NA 1.4, Olympus) and spectrally filtered (Di01-R405/488/561/635 dichroic, Semrock; ZET647NF notch filter, Chroma, Bellows Falls, VT; ET700/75m bandpass filter, Chroma, 3RD650LP longpass filter, Omega Optical, Austin, TX), and focused by the microscope tube lens. The emitted light entered a 4f imaging system (f= 90 mm) and was focused onto an EMCCD camera (iXon3 897, Andor, Belfast, UK) by the second lens of the 4f imaging system.

PBS was replaced by a reducing, oxygen scavenging buffer (20), consisting of 20 mM cysteamine, 2 µL/mL catalase, 560 µg/mL glucose oxidase (all Sigma-Aldrich), 10% (w/v) glucose (BD Difco, Franklin Lakes, USA), and 100 mM Tris-HCl (Life Technologies). Imaging was performed at 647 nm excitation with a laser intensity of 5 kW/cm^2^. The exposure time was 50 ms and the calibrated EM gain was 186. SR reconstructions were reconstructed from approx. 40000 frames using the ImageJ plugin Thunderstorm (21). Images were filtered with a B-spline filter of order 3 and scale 2.0. Single-molecule signals were detected with 8-neighborhood connectivity and a threshold of three times the standard deviation of the first wavelet level. Detected local maxima were fitted with a 2D-Gaussian using least squares. Drift correction was done by cross-correlation, followed by filtering (σ of the fitted Gaussian <300 nm; uncertainty of localization <30 nm). Images were reconstructed as 2D histograms with a bin size of 32 nm, corresponding to a five-time magnification compared to the pixel size of 160 nm.

### PEG Mass Tagging

Mass tagging was performed according to Woo et al. (22). Briefly, K-562 or K-562 GALE-KO cells stably transfected with pSBtet-AGX1^WT^ or pSBtet-AGX1^F383A^ were fed with azide-containing sugars as described above, and lysed using approx. 100 µL /500,000 cells of Lysis Buffer supplemented with 50 µM PUGNAc (Sigma Aldrich). Lysate corresponding to 30 µg protein was treated with 20% (v/v) to a final concentration of 1% and incubated for 10 min at 65 °C. The solution was treated with iodoacetamide (Sigma Aldrich) to a final concentration of 15 mM, and incubated in the dark for 30 min at room temperature. DIBAC-PEG 10 kDa (DBCO-PEG 10, Jena Bioscience) was added to a final concentration of 200 µM, and the solution was incubated overnight at room temperature. A 1:1 (v/v) mixture of 1 M Tris-HCl (pH 6.5) and 80% (v/v) glycerol was added (10% of the final volume), and mass tagging was assessed by Western Blot.

### Genome-wide CRISPR knockout screen

A CRISPR sgRNA library targeting all 20,500 human protein-coding genes was synthesized, cloned and packaged into lentivirus as described previously (23). 250 million K-562-Cas9 cells stably transfected with pSBtet-AGX1^F383A^ were infected with lentivirus at a multiplicity of infection of 0.4 for 24 h in media containing 8 µg/mL polybrene. Cells were subsequently selected for 128 h with 1 µg/mL puromycin, then changed into fresh media without puromycin and allowed to recover for 24 h. Cells were pooled, harvested in four batches of 50 million cells each, and resuspended in 200 mL medium each. Cells were treated with caged GalNAc-1-phosphate analog **5** (100 µM as 200 µL of a 100 mM stock solution in DMSO diluted to 2 mL with medium), DMSO (200 µL diluted to 2 mL with medium), or Ac_4_GalNAz (10 µM as 200 µL of a 10 mM stock solution in DMSO diluted to 2 mL with medium). Two t = 0 samples of 100 million cells each were washed with PBS once and frozen at −80 C.

After 20 h, cells were harvested (500 *x g*, 5 min, 4 °C) and washed twice with Labeling Buffer. The DMSO-treated samples were frozen at −80 °C except for a 400 µL aliquot that was harvested and resuspended in 400 µL 50 µM MB488-DIBAC (MB488-DBCO, Click Chemistry Tools) in Labeling Buffer.

GalNAc analog treated samples were resuspended in 50 µM MB488-DIBAC in Labeling Buffer (50 mL, MB488-DBCO, Click Chemistry Tools). Samples were incubated for 30 min at room temperature. Cells were harvested, washed twice with Labeling Buffer resuspended as a 15 million cells/mL suspension containing 5 nM SYTOX Red. Intact, viable cells were defined by sorting on FSC/SSC and SYTOX Red channels. A cell population representing both the top and bottom 15% of the fluorescence distribution for GalNAz/GalNAzMe was then isolated. Sorting was conducted until at least 50 million events (cells) had been processed for each sample. Sorted cells were then pelleted and frozen at −80 °C in preparation for subsequent processing. Aliquots of 50 million unsorted cells from a DMSO-treated sample were also pelleted and frozen down in parallel for normalization.

#### CRISPR Screen DNA Extraction and Data Analysis

Frozen cell pellets were thawed and genomic DNA extraction was performed using either the QIAamp DNA Blood Maxi Kit (Qiagen, Hilden, Germany) for unsorted samples or the GeneElute Mammalian Genomic DNA Miniprep kit (Sigma) for sorted samples according to manufacturer’s specifications. The sgRNA-encoding regions were amplified via nested PCR and sequenced on a NextSeq500 (Illumina, San Diego, USA). Reads were aligned to the sgRNA library and the log2 fold change was calculated for each sgRNA. Median phenotypes for each gene were calculated as previously described (24). P-values were calculated using a Mann-Whitney U-test and adjusted FDRs were computed using the Benjamini–Hochberg procedure.

### Click & enrichment of HepG2 secretome

HepG2 cells stably transfected with pSBtet-AGX1^F383A^ were seeded into one 10 cm dish per treatment (8 mL) without hygromycin B. After 24 h (40% confluency), cells were fed with either GalNAzMe (100 µM), Ac_4_GalNAz (3µM), or DMSO vehicle. After another 24 h, the medium was aspirated, cells were washed with pre-warmed serum-free low-Glucose DMEM and fed with GalNAzMe (100 µM), Ac_4_GalNAz (3µM) or DMSO in serum-free medium and incubated for 20 h. Conditioned supernatant was collected and centrifuged at 500 *x g* for5 min. The supernatant was concentrated to 2 mL using an Amicon Ultra-15 Centrifugal Filter Unit (3 kDa MWCO, Merck). Samples were treated with PNGase F (Promega, 5 µL of a 1:10 dilution in PBS) and incubated for 4 h at 37 °C. Then, azide-containing reaction mixtures were sequentially treated with 1200 µM BTTAA (stock solution 50 mM in 9:1 DMSO : water), 600 µM CuSO4 (stock solution 20 mM in water), 5 mM sodium ascorbate, 5 mM aminoguanidine chloride, and 100 µM DADPS Biotin Alkyne (Click Chemistry Tools, stock solution 10 mM in DMSO). The click reaction was carried out for 3 h at room temperature with inversion. Then, samples were transferred into 15 mL Falcon tubes and treated with 10 mL (5-fold excess) ice-cold methanol. Samples were left at −80 °C overnight, when a white precipitate had formed. Samples were centrifuged at 3700 *x g*, 4 °C, 20 min. Supernatant was discarded, and pellets were washed with 5 mL methanol twice, with centrifugation each time. Supernatant was completely removed by air-drying, and samples were treated with 250 µL 0.1% RapiGest in PBS. Samples were sonicated (water bath) for 25 min, then centrifuged at 3700 *x g* for 5 min. Supernatant was saved, and pellets were treated with 250 µL 6 M urea in PBS. Samples were sonicated and centrifuged again, and the supernatant was saved. The pellets were treated with 250 µL of PBS, sonicated and centrifuged again. RapiGest, urea and PBS supernatants were combined, and samples were diluted with PBS to 2 mL. Dimethylated Sera-Mag SpeedBeads Neutravidin Magnetic Beads (150 µL slurry = 75 µL settled resin) were washed with PBS twice in LoBind tubes (Eppendorf) and added to the lysate (25, 26). Samples were incubated for 16 h at 4 °C under rotation. The beads were harvested and the supernatant was discarded. The beads were washed sequentially with 1% RapiGest in PBS (3x), 6 M urea in PBS (3x), and PBS (2x), and resuspended in PBS (200 µL). Beads were treated with 100 mM DTT in PBS (10 µL), and shaken for 30 min, at room temperature, 950 rpm. Then, 500 mM iodoacetamide in PBS (4 µL) was added and samples were shaken for another 30 min in the dark. Beads were harvested and washed with PBS and 50 mM ammonium bicarbonate in LC/MS-grade water (3x, “ABC buffer”). Beads were resuspended in ABC buffer (200 µL), treated with RapiGest to a final concentration of 0.05% (v/v), and LysC was added (500 ng in 3 µL of ABC buffer). Samples were shaken at 37 °C for 2-3 h, and another 500 ng LysC was added. The reactions were shaken overnight at 37 °C. The beads were harvested, washed with ABC (200 µL) and with LC-MS grade water (3×200 µL). Supernatants and washes were combined and centrifuged (18000 *x g*, 5 min, RT) and concentrated by SpeedVac which formed the peptide fraction to be analysed by mass spectrometry. Then, beads were treated with 150 µL of 0.1% aq. formic acid (FA, Optima grade, Thermo Fisher) and shaken for 30 min, at room temperature, 950 rpm. This step was repeated, beads were washed with 100 µL of water, all washes were combined and centrifuged (18000 *x g*, 5 min, room temperature), and finally concentrated by SpeedVac. Remaining beads were treated with 2% FA and subjected to the same washes as above. All samples were then resuspended with 25 ng trypsin in 100 µL of 50 mM ABC buffer and incubated for 6 h at 37 °C, 450 rpm. Samples were then dried by SpeedVac and desalted by Strata-X columns using 0.1% aq. FA as washing solution and 80% MeCN/water with 0.1% FA (v/v) as elution buffer. The eluted samples dried by SpeedVac.

Peptides dried by vacuum centrifugation into 0.5 mL LoBind tubes (Eppendorf) were resuspended into 16 µL of 0.1 % (v/v) FA with sonication using an ultrasonic water bath followed by vortexing. The solubilised peptides were centrifuged for 5 min at 18,000 *x g* and the solution transferred into Total Recovery vials (Waters, Milford, USA) for injection. Samples were analysed by online nanoflow LC-MS/MS using an Orbitrap Fusion Lumos mass spectrometer (Thermo Scientific) coupled to an Ultimate 3000 RSLCnano (Thermo Scientific). Sample (15 µL) was loaded via autosampler into a 20 µL sample loop and pre-concentrated onto an Acclaim PepMap 100 75 µm x 2 cm nanoviper trap column with loading buffer, 2% v/v acetonitrile, 0.05% v/v trifluoroacetic acid, 97.95% water (Optima grade, Fisher Scientific) at a flow rate of 7 µL/min for 6 min in the column oven held at 40 °C. Peptides were gradient eluted onto a C_18_ 75 µm x 50 cm, 2 µm particle size, 100Å pore size, reversed phase EASY-Spray analytical column (Thermo Scientific) at a flow rate of 275 nL/min and with the column temperature held at 40 °C, and a spray voltage of 2100 V using the EASY-Spray Source (Thermo Scientific). Gradient elution buffers were A 0.1% v/v FA, 5% v/v DMSO, 94.9% v/v water and B 0.1% v/v FA, 5% v/v DMSO, 20% v/v water, 74.9% v/v acetonitrile (all Optima grade, Fisher Scientific aside from DMSO, Honeywell Research Chemicals). The gradient elution profile was 2% B to 40% B over 98 minutes. The instrument method used an MS1 Orbitrap scan resolution of 120,000 at FWHM m/z 200, quadrupole isolation, mass range 300-1500 m/z, RF Lens 30%, AGC target 4e5, maximum injection time 50 ms and spectra were acquired in profile. Monoisotopic Peak Determination was set to the peptide mode, and only precursors with charge states 2-6 were permitted for selection for fragmentation. Dynamic Exclusion was enabled to exclude after n=3 times within 10 s for 10 s with high and low ppm mass tolerances of 10 ppm. HCD was performed on all selected precursor masses using a cycle time-based data dependant mode of acquisition set to 3 s. MS2 scans were acquired in the Orbitrap at a resolution of 30000 FWHM m/z 200, following HCD fragmentation with fixed collision energy of 28% after quadrupole isolation with an isolation window width of 2 m/z. The parameters used for the HCD MS2 scan were first mass 100 m/z, AGC target 5e4, maximum injection time 54 ms and the scan data was acquired in centroid mode. ETD fragmentation was only performed if precursors were within the precursor selection range m/z 300-1000 and if 2 of the following list of mass trigger ions were present in the HCD MS2 spectra +/- 0.1 m/z and above the relative intensity threshold of 10% (126.055, 138.0549, 144.0655, 168.0654, 186.076, 204.0855, 274.0921, 292.1027, 343.1617, 329.1461 m/z). ETD MS2 scans were recorded in the ion trap with rapid scan rate following quadrupole isolation with an isolation window width of 3 m/z. ETD activation used calibrated charge-dependent ETD parameters, the automatic scan range mode was used with the first mass set to 100 m/z and parameters were set for the AGC target 1e4, maximum injection time 100 ms and scan data acquired in centroid mode.

Data evaluation was performed with Byonic™ (Protein Metrics, Cupertino, USA). Data files were first searched against the Uniprot human proteome (downloaded June 26, 2016). Search parameters included semi-specific cleavage specificity at the C-terminal site of R and K, with two missed cleavages allowed. Mass tolerance was set at 10 ppm for MS1s, 0.1 Da for HCD MS2s, and 0.35 Da for ETD MS2s. Methionine oxidation (common 2), asparagine deamidation (common 2), and N-term acetylation (rare 1) were set as variable modifications with a total common max of 3, rare max of 1. Cysteine carbamidomethylation was set as a fixed modification. Peptide hits were filtered using a 1% FDR. Additionally, a cut-off value of Log Prob = 5 was set for any further analysis. Proteins that were found in these searches were entered into a “focused database” for glycopeptide searches. Then, the raw files were searched against these “focused databases” containing only those proteins found in the corresponding peptide samples. Search parameters for the glycopeptide analysis included semi-specific cleavage specificity at the C-terminal site of R and K, with two missed cleavages allowed. Mass tolerance was set at 10 ppm for MS1s, 0.1 Da for HCD MS2s, and 0.35 Da for ETD MS2s. Methionine oxidation (common 2), asparagine deamidation (common 2), and N-term acetylation (rare 1) were set as variable modifications with a total common max of 2, rare max of 1. O-glycans were also set as variable modifications (common 2), using a custom database, whereby HexNAc, HexNAc-NeuAc, HexNAc-Hex, HexNAc-Hex-NeuAc, and HexNAc-Hex-NeuAc2 were searched with an additional 139.0746 (GalNAzMe) or 125.0589 (GalNAz) to account for the chemical modifications. HCD was used to confirm that the peptides were glycosylated, as the modified sugars have signature ions present at 343.1617/325.1506 m/z (GalNAzMe) or 329.1461/311.1435 m/z (GalNAz). Following confirmation, ETD spectra were used for site-localisation of glycosylation sites. All spectra with these modifications were manually annotated.

### Organoid culture and generation of stably overexpressing organoid lines

Organoids were established from freshly isolated wild type small intestine from adult mice, as previously described (27). Upon isolation intestinal crypts were cultured in Cultrex® BME, Type 2 RGF PathClear (Amsbio, 3533-010-02) and IntestiCult™ Organoid Growth Medium (Stem Cell technologies, #06005) was used to drive differentiation of all epithelial cell types. Generation of stably overexpressing organoid lines was performed as previously described (28). Briefly, organoids were dissociated in single cells with Accumax (Merck) and counted. 500,000 cells were electroporated using a NEPA21 electroporator with pSBtet-GalNAcT2^WT^-AGX1^F383A^ or pSBtet-GalNAcT2^DM^-AGX1^F383A^ and seeded at high confluency. Organoids were treated with Rho kinase inhibitor Y-27632 (Sigma Aldrich, Y0503) and Gsk3 inhibitor CHIR99021 (Tocris, 4423) for 48 h prior to and after electroporation and with DMSO for 24 h prior to and after electroporation to increase efficiency. After 4 days, organoids were treated with fresh medium containing 200 µg/ml Hygromycin B.

### Organoid labelling and immunofluorescence

Organoids were seeded on an 8-well glass chamber slide (Nunc, 154534), allowed to establish for 72 h and, subsequently, fed twice with 1.5 µM Ac_4_GalNAz or 50 µM compound **11** for 2 days. After feeding, organoids were washed twice with PBS and fixed with 10% Formalin for 20 min at room temperature. The reaction was quenched with addition of 50 mM NH_4_Cl for 5 min. Organoids were then treated with 150 µL PBS and 150 µL of a freshly prepared solution containing 1200 µM BTTAA, 200 µM CuSO4, 10 mM sodium ascorbate, 10 mM aminoguanidine chloride and 200 µM biotin-PEG3-alkyne (Click Chemistry Tools) for 10 min, then washed with PBS (3 x 300µL), and incubated with 20 µg/mL AF647-streptavidin (Jackson ImmunoResearch) for 30 min at 4 °C. After washing with PBS (3 x 300µL), organoids were permeabilized with 0.8 % Triton X-100 for 20 min at room temperature, blocked with 1 % BSA for 30 min at room temperature and incubated with Phalloidin-FITC (Merck, P5282) and 4’,6’-diamidino-2-phenylindole (DAPI) for 1 h at room temperature. Slides were washed (3 x 300µL) and mounted with ProLong Gold Antifade mountant (Thermo Fisher, P36934). Samples were imaged using a Leica TCS SPE confocal microscope.

### Synthetic chemistry

Solvents and reagents were of commercial grade. Anhydrous solvents were obtained from a Dry Solvent System. Water-sensitive reactions were carried out in heat-dried glassware and under a nitrogen atmosphere. Thin layer chromatography was performed on Kieselgel 60 F254 glass plates pre-coated with silica gel (0.25 mm thickness). Spots were developed with ceric ammonium molybdate stain (5% (w/v) ammonium molybdate, 1% (w/v) cerium (II) sulfate and 10% (v/v) sulfuric acid in water) or sugar stain (0.1% (v/v) 3-methoxyphenol, 2.5% (v/v) sulfuric acid in EtOH) dipping solutions. Flash chromatography was carried out on Fluka Kieselgel 60 (230-400 mesh). Solvents were removed under reduced pressure using a rotary evaporator and high vacuum (1 mbar). Medium pressure chromatography was performed on an Isolera Prime system (Biotage, Uppsala, Sweden).

1H, ^13^C and 2D NMR spectra were measured with an AS400 spectrometer, an AS600 spectrometer (Varian, Palo Alto, USA) or a Bruker Avance-400 MHz spectrometer at 298 K. Chemical shifts (σ) are reported in parts per million (ppm) relative to the respective residual solvent peaks (CDCl_3_: σ 7.26 in ^1^H and 77.16 in 13C NMR; acetone-D_6_: σ 2.05 in ^1^H and 29.84 in ^13^C NMR). Two-dimensional NMR experiments (HH-COSY, CH-HSQC) were performed to assign peaks in ^1^H spectra. The following abbreviations are used to indicate peak multiplicities: s singlet; d doublet; dd doublet of doublets; dt doublet of triplets; m multiplet. Coupling constants (*J*) are reported in Hertz (Hz). High resolution mass spectrometry by electrospray ionization (ESI-HRMS) was performed at Stanford University Mass Spectrometry, with a micrOTOF-Q II hybrid quadrupole time-of-flight mass spectrometer (Bruker, Billerica, USA) equipped with a 1260 UPLC (Agilent). Low resolution mass spectrometry spectrometry by electrospray ionization (ESI-LRMS) was performed on an UPLC-MS (Waters) equipped with ACQUITY UPLC® BEH C18 column. HPLC purification was performed on Perkin Elmer 200 Series equipped with ZORBAX 300SB-C8 column (21.2×250 mm, 7 µm), UV detector and a flow rate of 8 mL/min. Elution was monitored at 214 nm.

Compounds **SI-1, 3, 4, 5, 6, 7, 8, 9** and **10** were made previously (6). Compounds **1, 2**, UDP-GlcNAc and UDP-GlcNAz are commercially available (Thermo).

### Bis(*S*-acetyl-2-thioethyl) 3,4,6-tri-*O*-acetyl-2-[2-(*S*)-azidopropionamido]-2-deoxy-2-α-D-galactopyranosyl phosphate (11)

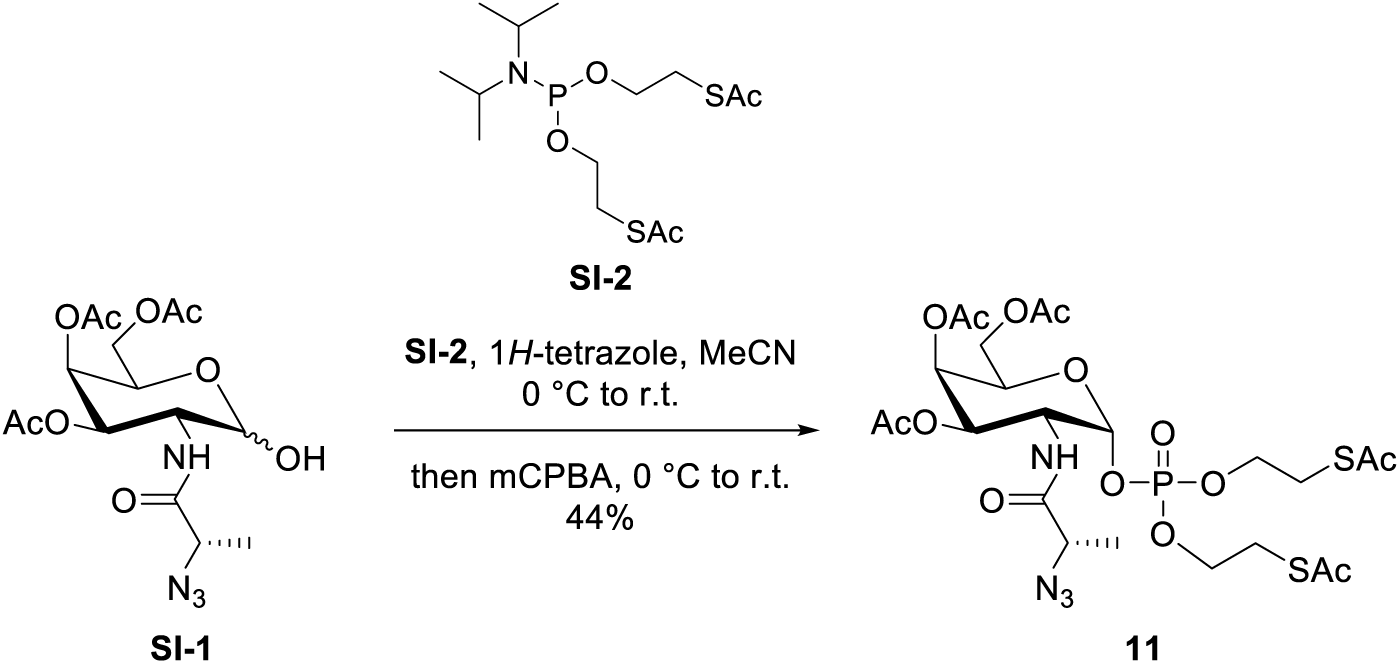

To a stirred solution of lactol **SI-1**(6) (100 mg, 249 µmol) and phosphoramidite **SI-2**(29) (128 mg, 347 µmol) in MeCN (1.6 mL) was added at 0 °C 1*H*-tetrazole (26 mg, 371 µmol, 867 µL of a 3% (w/v) solution in MeCN). The reaction was warmed to room temperature and stirred for 1 h. The mixture was cooled to 0 °C and treated with mCPBA (64 mg, 372 µmol). The mixture was warmed to room temperature, stirred for 30 min, diluted with EtOAc (10 mL) and quenched with 10% aq. Na_2_SO_3_ (10 mL). The solution was washed with 10% aq. Na_2_SO_3_ (10 mL), sat. aq. NaHCO_3_ (2×30 mL) and brine (50 mL). After every washing step, the aqueous layer was back-extracted with EtOAc (10 mL). The combined organic layers were dried over MgSO_4_, filtered and concentrated. The residue was purified by flash chromatography (hexanes/EtOAc 1:1 to 0:1) to give phosphotriester **11** (75 mg, 109 µmol, 44%) as a clear oil. ^1^H NMR (400 MHz, acetone-D_6_) δ 7.59 (d, *J* = 8.3 Hz, 1H), 5.88 – 5.77 (m, 1H), 5.58 – 5.47 (m, 1H), 5.25 (dd, *J* = 11.7, 3.2 Hz, 1H), 4.63 – 4.45 (m, 2H), 4.32 – 3.98 (m, 7H), 3.30 – 3.14 (m, 4H), 2.36 (d, *J* = 1.2 Hz, 6H), 2.16 (s, 3H), 2.01 (s, 3H), 1.94 (s, 3H), 1.45 (d, *J* = 6.9 Hz, 3H); ^13^C NMR (100 MHz, acetone-D_6_) δ 195.2, 195.0, 172.0, 170.7, 170.6, 170.5, 97.2, 69.6, 67.9, 67.8, 67.1, 67.0, 67.0, 62.4, 58.7, 48.7, 20.7, 20.6, 20.6, 17.5; HRMS (ESI) calcd. for C_23_H_35_N_4_O_14_PS_2_ (M+Na+) 709.1227 found 709.1219 *m/z*.

### 2-[(*S*)-Azidopropionamido]-2-deoxy-1,3,4,5-tetra-*O*-acetyl-D-glucopyranose (SI-3)

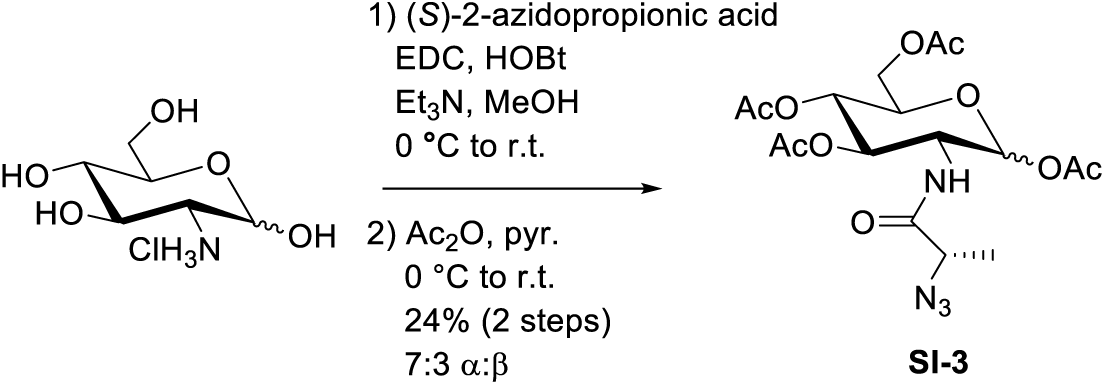

To a stirred solution of acid (*S*)-2-azidopropionic acid(30) (1.7 g, 14.8 mmol) in MeOH (60 mL) were added D-glucosamine hydrochloride (1.32 g, 6.1 mmol) and triethylamine (2.28 mL, 16.28 mmol). The solution was cooled to 0 °C, and EDC (2.53 g, 16.28 mmol) and HOBt (1.13 g, 7.4 mmol) were added. The reaction was warmed to room temperature and was stirred for 72 h. The mixture was concentrated and passed twice through a plug of silica gel (CH_2_Cl_2_/MeOH 9:1) to give a residue that was dissolved in MeOH (2.5 mL), treated with CHCl_3_ (100 mL) and precipitated at −20 °C to give the intermediary amide as a yellow precipitate that still contained traces of EDC and HOBt.

To a stirred solution of the intermediary amide in anhydrous pyridine (11 mL) was added at 0 °C acetic anhydride (6.3 mL, 66.3 mmol). The reaction was warmed to room temperature and stirred for 6 h. The mixture was diluted with water (100 mL) and EtOAc (100 mL), and the layers partitioned. The aqueous phase was extracted with EtOAc (3×50 mL), the combined organic extracts were washed with 0.2 M aq. HCl (3×50 mL), water (50 mL) and brine (50 mL), dried over MgSO_4_, filtered and concentrated. The residue was co-evaporated with toluene (20 mL) and purified by flash chromatography (hexanes/EtOAc 1:0 to 1:3 to 1:2 to 1:1) to give tetraacetate **SI-3** (642 mg, 1.44 mmol, 24% over two steps, 7:3 α:β) as a clear oil. ^1^H NMR (400 MHz, CDCl_3_) δ 6.78 – 6.62 (m, 0.3H), 6.45 (d, *J* = 8.7 Hz, 0.7H), 6.17 (t, *J* = 3.7 Hz, 0.7), 5.76 (dd, *J* = 8.7, 1.7 Hz, 0.3H), 5.37 – 4.96 (m, 2H), 4.45 – 3.76 (m, 5H), 2.27 – 1.79 (m, 12H), 1.50 – 1.30 (m, 3H); ^13^C NMR (100 MHz, CDCl_3_) δ 171.5, 170.8, 170.7, 170.7, 170.5, 170.2, 169.4, 169.3, 169.2, 168.7, 92.3, 90.2, 72.9, 72.0, 70.3, 69.9, 68.0, 67.5, 61.7, 61.6, 59.3, 59.1, 53.0, 51.4, 29.7, 20.9, 20.8, 20.8, 20.7, 20.7, 20.7, 20.7, 20.6, 17.2, 17.2. HRMS (ESI) calcd. for C_17_H_24_N_4_O_10_ (M+Na^+^) 467.1390 found 467.1387 *m/z*.

### 2-[(*S*)-Azidopropionamido]-2-deoxy-3,4,5-tri-*O*-acetyl-D-glucopyranose (SI-4)

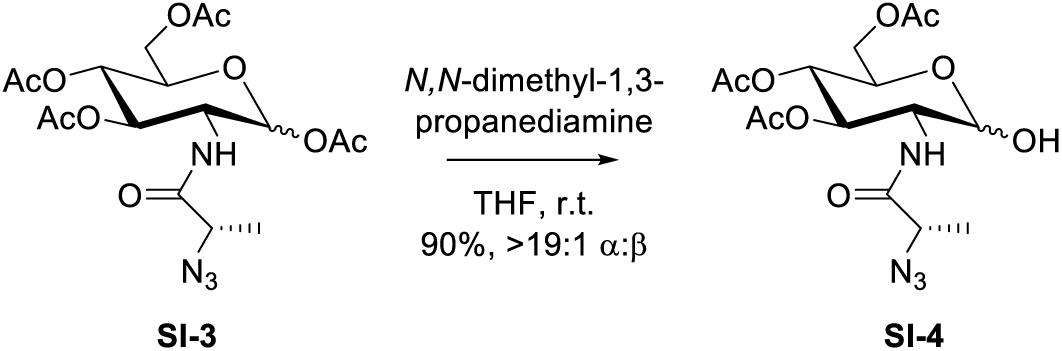

To a stirred solution of tetraacetate **SI-3** (320 mg, 0.72 mmol) in THF (3.5 mL) was added 3- (dimethylamino)-1-propylamine (0.26 mL, 2.15 mmol). The reaction was stirred at room temperature for 2 h, diluted with CH_2_Cl_2_ (50 mL), washed with 1 N aq. HCl (2×20 mL) and brine (20 mL). The organic phase was dried over MgSO_4_, filtered and concentrated. The residue was purified by flash chromatography (hexanes/EtOAc 1:0 to 2:1) to give lactol **SI-4** (263 mg, 0.65 mmol, 90%, >19:1 α:β) as a clear oil. Rf (hexanes/EtOAc 3:2) = 0.35. ^1^H NMR (400 MHz, CDCl_3_) δ 6.68 (d, *J* = 9.3 Hz, 1H), 5.31 (t, *J* = 9.4 Hz, 1H), 5.23 (s, 1H), 5.11 (t, *J* = 9.4 Hz, 1H), 4.29 – 4.17 (m, 3H), 4.14 – 4.06 (m, 1H), 4.01 – 3.92 (m, 1H), 2.07 (s, 3H), 2.02 (s, 3H), 1.99 (s, 3H), 1.46 (d, *J* = 7.1 Hz, 3H); ^13^C NMR (100 MHz, CDCl_3_) δ 171.3, 171.1, 170.6, 169.6, 91.4, 70.9, 68.3, 67.6, 62.2, 58.9, 52.5, 20.9, 20.8, 20.7, 17.2; HRMS (ESI) calcd. For C_15_H_22_N_4_O_9_P (M+Na^+^) 425.1285 found 425.1283 *m/z*.

### Bis-*O*-allyl 2-[(*S*)-Azidopropionamido]-2-deoxy-3,4,5-tetra-*O*-acetyl-α-D-glucopyranosyl phosphate (SI-5)

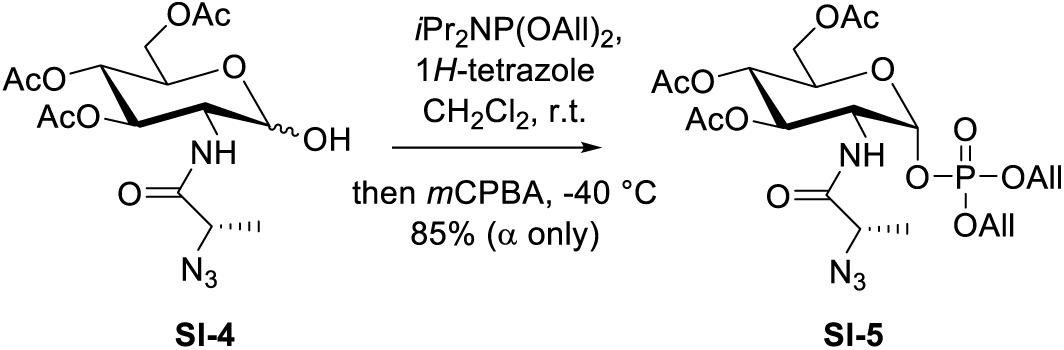

Lactol **SI-4** (210 mg, 0.52 mmol) and 1*H*-tetrazole (174 mg, 2.48 mmol) were co-evaporated with anhydrous toluene (5 mL), suspended in anhydrous toluene (5 mL) and sonicated for 1 h at room temperature in a bath sonicator. The solvent was evaporated, and the residue dissolved in anhydrous CH_2_Cl_2_ (7.8 mL). The stirred solution was cooled to 0 °C, and diallyl *N,N*-diisopropylphosphoramidite (220 µL, 0.83 mmol) was added. After 20 min, the solution was cooled to −40 °C, and mCPBA (222 mg, 0.99 mmol) was added. Another 126 mg (0.56 mmol) mCPBA was added after 10 min to drive the reaction to completion. After another 10 min, the reaction was quenched with 20% aq. Na_2_SO_3_ (10 mL), and warmed to room temperature. The mixture was diluted with CH_2_Cl_2_ (20 mL), and the layers were separated. The organic phase was washed with sat. aq. NaHCO_3_ (10 mL), and the combined aqueous phase was re-extracted with CH_2_Cl_2_ (20 mL). The combined organic phase was washed with brine (20 mL), the organic phase wash dried over MgSO_4_, and concentrated. The residue was purified by flash chromatography (hexanes/EtOAc 1:0 to 1:2 to 3:2) to give phosphate **SI-5** (248 mg, 0.44 mmol, 85%, α anomer only) as a clear oil. Rf (hexanes/EtOAc 3:2) = 0.3. ^1^H NMR (400 MHz, CDCl_3_) δ 6.74 (d, *J* = 8.9 Hz, 1H), 6.00 – 5.81 (m, 2H), 5.67 (dd, *J* = 6.2, 3.3 Hz, 1H), 5.41 – 5.19 (m, 5H), 5.14 (t, *J* = 9.7 Hz, 1H), 4.61 – 4.49 (m, 4H), 4.31 (m, 1H), 4.24 – 4.12 (m, 2H), 4.05 (dd, *J* = 12.2, 2.0 Hz, 1H), 3.96 (q, *J* = 7.0 Hz, 1H), 2.03 (s, 3H), 2.00 (s, 3H), 1.98 (s, 3H), 1.44 (d, *J* = 7.0 Hz, 3H); ^13^C NMR (100 MHz, CDCl_3_) δ 171.1, 170.6, 170.5, 169.2, 132.0, 132.0, 132.0, 131.9, 119.1, 119.0, 95.7, 95.6, 69.9, 69.7, 68.9, 68.85, 68.8, 68.7, 67.4, 61.5, 58.9, 52.2, 52.1, 20.7, 20.6, 20.6, 17.2; HRMS (ESI) calcd. for C_21_H_31_N_4_O_12_P (M+Na^+^) 585.1573 found 585.1565 *m/z*.

### Uridine 5’-diphospho-2-((*S*)-azidopopionamido)-2-deoxy-α-D-glucopyranoside tributylammonium salt (SI-7)

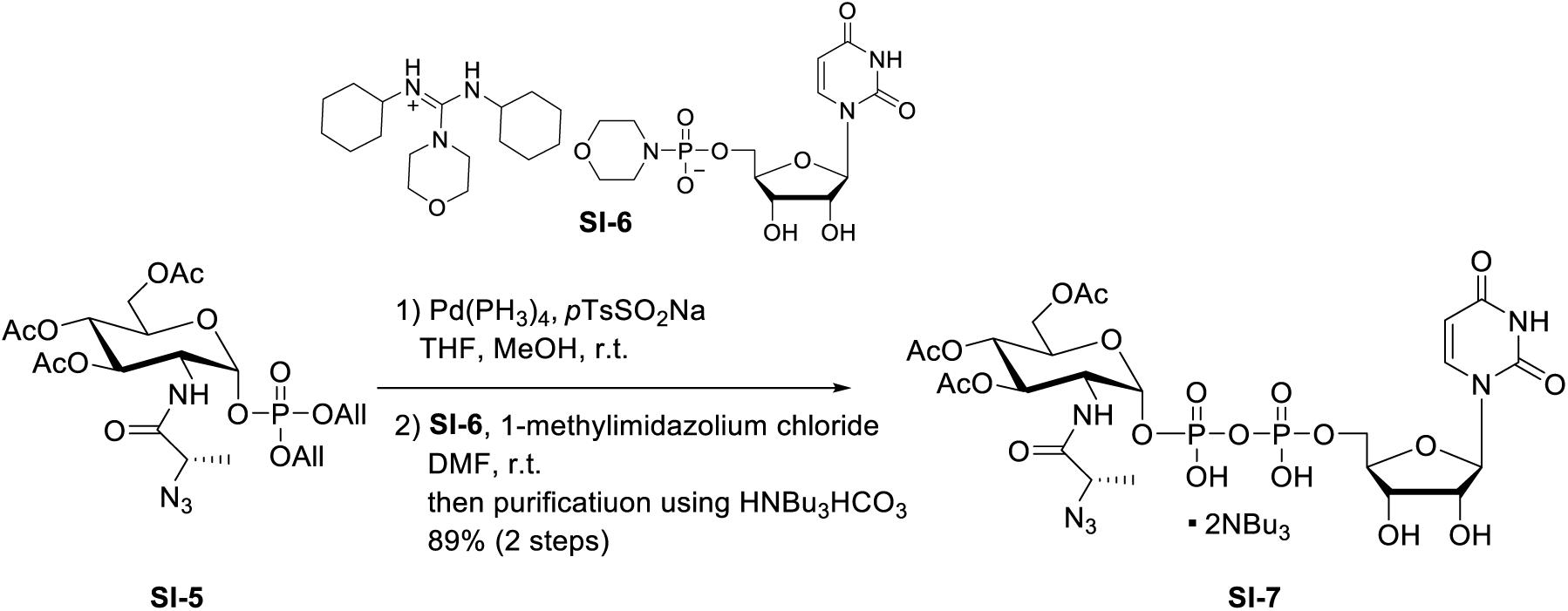

To a stirred solution of diallyl phosphotriester **SI-5** (50 mg, 89 µmol) in THF/MeOH (1:1, 1.78 mL) were added tetrakis(triphenylphosphine)palladium (5 mg, 4.5 µmol) and sodium *para*-toluoenesulfinate (32 mg, 178 µmol). The reaction was stirred at room temperature for 3 h, and tetrakis(triphenylphosphine)palladium (5 mg, 4.5 µmol) and sodium *para*-toluoenesulfinate (32 mg, 178 µmol) were added to drive the reaction to completion. The mixture was stirred at room temperature for 16 h, and the solvents were evaporated. The residue was co-evaporated with anhydrous toluene (2×5 mL) and dissolved in DMF (1.78 mL). Uridine monophosphomorpholidate **SI-6** (98 mg, 145 µmol) and 1-methylimidazolium chloride (57 mg, 0.49 µmol) were added and the reaction was stirred at room temperature for 16 h. The mixture was concentrated and purified by medium-pressure flash chromatography (60 g SNAP C18 column; A: 10 mM tributylammonium bicarbonate,(6) B: MeOH; 6 column volumes 100% A; 9 column volumes linear gradient to 100% B, then 6 column volumes 100% B), and fractions were concentrated and lyophilized repeatedly to give pyrophosphate **SI-7** as the tributylammonium salt (92 mg, 79 µmol, 89% over two steps) as a white foam. Rf (CH_2_Cl_2_/MeOH 2:1 + 1% AcOH) = 0.3. ^1^H NMR (600 MHz, CD_3_OD) δ 8.08 (d, *J* = 8.2 Hz, 1H), 5.93 (d, *J* = 4.4 Hz, 1H), 5.81 (d, *J* = 8.1 Hz, 1H), 5.65 (dd, *J* = 7.3, 3.3 Hz, 1H), 5.28 (t, *J* = 10.0 Hz, 1H), 5.11 (t, *J* = 9.8 Hz, 1H), 4.47 – 4.04 (m, 10H), 2.05 (s, 3H), 1.98 (s, 3H), 1.92 (s, 3H), 1.37 – 1.33 (m, 2H); 13C NMR (150 MHz, CD_3_OD) δ 174.6, 172.5, 171.7, 171.4, 166.3, 159.3, 152.7, 142.7, 103.1, 95.8, 90.3, 85.0 84.9, 75.9, 73.4, 70.7, 69.9, 69.6, 65.8, 62.9, 58.2, 56.0, 55.3, 55.2, 53.9, 53.1, 53.1, 22.6, 21.1, 20.7, 20.7, 18.1; HRMS (ESI) calcd. for C_24_H_33_N_6_O_20_P2 (M-H^+^) 787.1225 found 787.1227 *m/z*.

### Uridine 5’-diphospho-2-((*S*)-azidopopionamido)-2-deoxy-α-D-glucopyranoside sodium salt (SI-8)

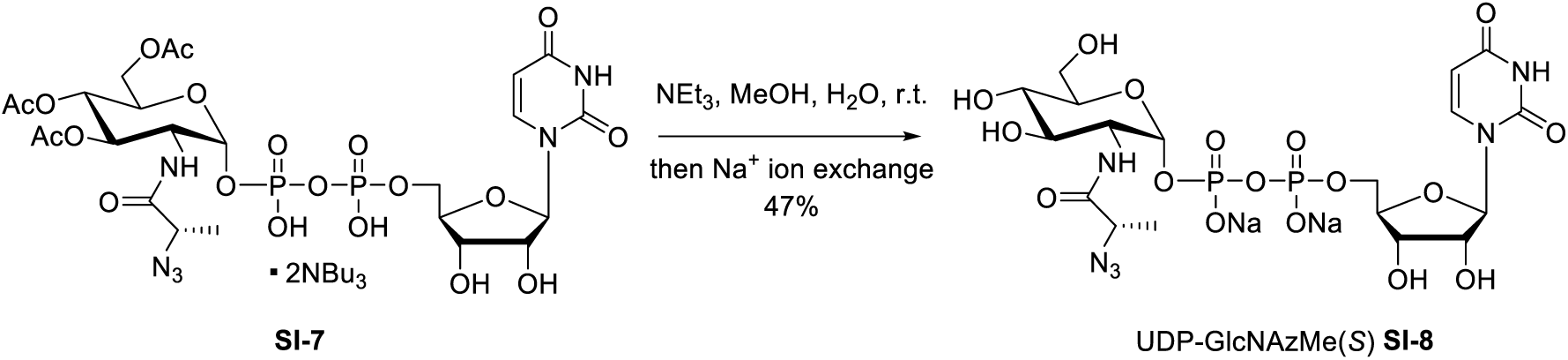

To a stirred solution of triester **SI-7** (8.3 mg, 7.2 µmol) in MeOH/water (5:2, 1.5 mL) was added triethylamine (300 µL). The reaction was stirred at room temperature for 16 h and concentrated. The residue was lyophilized repeatedly, passed through a short (3 g resin) ion exchange column (Dowex 50W X8 Na^+^ form, Sigma Aldrich), and concentrated. The residue was purified by reverse-phase solid-phase extraction (HyperSep™ C18, Thermo Fisher Scientific, Waltham, USA) and lyophilized to give UDP-GlcNAzMe(*S*) **SI-8** as the disodium salt (2.4 mg, 3.4 µmol, 47%) as a white solid. ^1^H NMR (600 MHz, D_2_O) δ 7.97 (d, *J* = 8.1 Hz, 1H), 6.03 – 5.86 (m, 2H), 5.56 (dd, *J* = 7.0, 3.3 Hz, 1H), 4.44 – 4.35 (m, 2H), 4.34 – 4.23 (m, 3H), 4.22 – 4.15 (m, 1H), 4.05 (m, 1H), 3.96 (m, 1H), 3.92 – 3.80 (m, 3H), 3.57 (dd, *J* = 10.1, 9.2 Hz, 1H), 1.50 (d, *J* = 7.0 Hz, 3H); ^13^C NMR (150 MHz, D_2_O) δ 174.2, 141.4, 102.6, 94.5, 94.4, 88.4, 83.1, 83.0, 73.7, 73.0, 70.8, 69.6, 69.5, 64.9, 60.2, 58.1, 53.6, 53.6, 16.8; HRMS (ESI) calcd. for C_9_H_15_N_4_O_5_^+^ (oxacarbenium ion) 259.1042 found 259.1046 *m/z*.

### 2-[3-Azidopropionamido]-2-deoxy-1,3,4,5-tetra-*O*-acetyl-D-glucopyranose (SI-10)

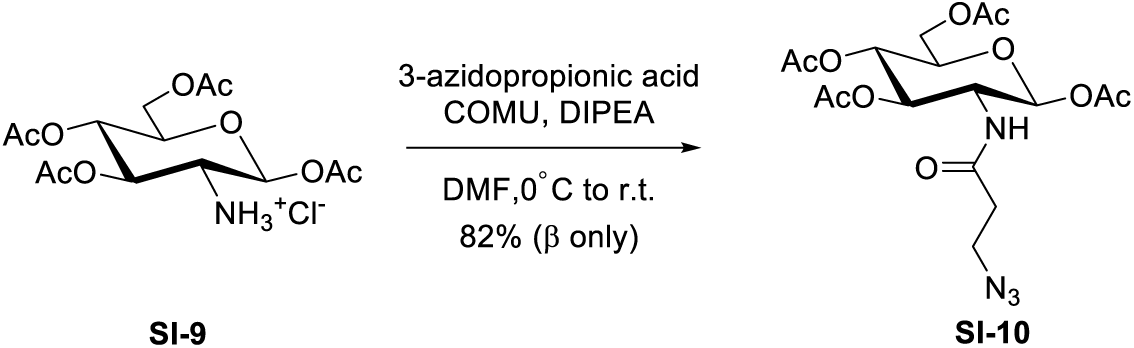

A mixture of peracetylated glucosamine hydrochloride **SI-9** (31) (384 mg, 1 mmol), 3-azidopropionic acid (93 µl, 1 mmol) and DIPEA (0.522 ml, 3 mmol) in DMF (8 ml) was cooled to 0 °C. COMU (856 mg, 2 mmol) was added and the reaction mixture stirred at 0 °C for 1 h. The solution was warmed to room temperature and stirred for another 3 h. The mixture was diluted with EtOAc (100 ml) and the organic layer was washed with 1 N aq. HCl (2×50 mL), sat. NaHCO_3_ (2×50 ml) and brine, dried over MgSO_4,_ filtered and concentrated. The residue was purified by medium-pressure flash chromatography (25g SNAP-KP-SIL; A: cyclohexane, B: EtOAc; 20 CV linear gradient from 30% to 70%, then 5 CV from 70% to 100%B) to give tetraacetate **SI-10** (366 mg, 0.82 mmol, 82% β-anomer only) as a clear oil. ^1^H NMR (400 MHz, CDCl_3_) δ 5.72 (d, *J* = 8.8 Hz, 1H), 5.66 (d, *J* = 9.4 Hz, 1H), 5.20 – 5.11 (m, 2H), 4.36 – 4.24 (m, 2H), 4.13 (dd, *J* = 12.5, 2.3 Hz, 1H), 3.81 (m, 1H), 3.62 – 3.55 (m, 2H), 2.34 (m, 2H), 2.12 (s, 3H), 2.09 (s, 3H), 2.05 (d, *J* = 3.7 Hz, 6H); ^13^C NMR (100 MHz, CDCl_3_) δ 171.3, 170.7, 170.0, 169.5, 169.2, 92.5, 73.0, 72.4, 67.7, 61.6, 53.2, 47.1, 35.9, 20.8, 20.7, 20.6; LRMS (ESI) calcd. for C_17_H_24_N_4_O_10_ (M-H^+^) 444.4 found 443.2 *m/z*.

### 2-[3-Azidopropionamido]-2-deoxy-3,4,5-tri-*O*-acetyl-D-glucopyranose (SI-11)

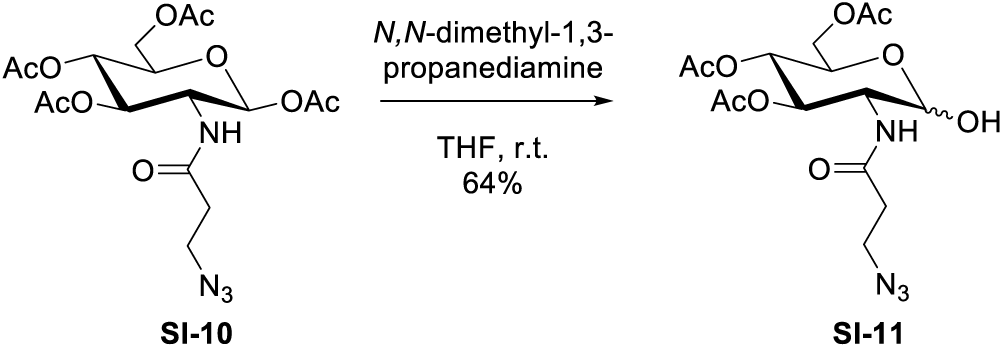

To a stirred solution of tetraacetate **SI-10** (151 mg, 0.34 mmol) in THF (1.65 mL) was added 3- (dimethylamino)-1-propylamine (0.123 mL, 1.02 mmol). The reaction was stirred at room temperature for 2 h, diluted with CH_2_Cl_2_ (25 mL), washed with 1 N aq. HCl (2×10 mL) and brine (10 mL). The organic phase was dried over MgSO_4_, filtered and concentrated. The residue was purified by medium-pressure flash chromatography (10 g SNAP; A: cyclohexane, B: EtOAc; 20 CV linear gradient from 30% to 100% B, then 5 CV 100% B) to give tetraacetate **SI-11** (88 mg, 0.22 mmol, 64%, >19:1 α:β) as a clear oil. ^1^H NMR (400 MHz, CDCl_3_) δ 5.91 (d, *J* = 9.3 Hz, 1H), 5.35 – 5.28 (m, 2H), 5.18 – 5.11 (m, 1H), 4.35 (m, 1H), 4.26 – 4.10 (m, 3H), 3.59 (t, *J* = 6.3 Hz, 2H), 3.01 (dd, *J* = 3.8, 1.6 Hz, 1H), 2.39 (td, *J* = 6.2, 4.0 Hz, 2H), 2.10 (s, 3H), 2.04 (d, *J* = 2.3 Hz, 6H). ^13^C NMR (100 MHz, CDCl_3_) δ 171.5, 171.0, 170.1, 169.4, 91.6, 70.8, 68.2, 67.7, 62.1, 52.3, 47.2, 35.7, 20.8, 20.7, 20.6; LRMS (ESI) calcd. for C_15_H_22_N_4_O_9_ (M-H^+^) 402.14 found 401.21 *m/z*.

### Bis-*O*-allyl 2-[3-Azidopropionamido]-2-deoxy-3,4,5-tetra-*O*-acetyl-α-D-glucopyranosyl phosphate (SI-12)

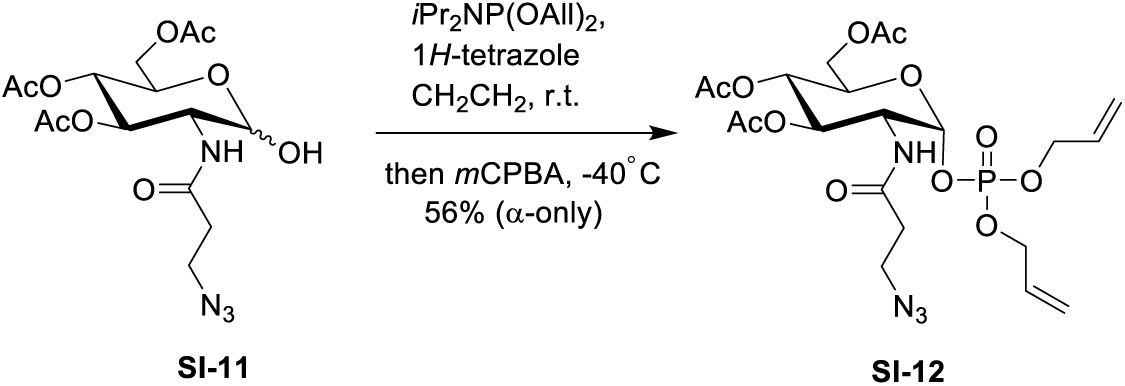

Lactol **SI-11** (88 mg, 0.22 mmol) and 1*H*-tetrazole (69 mg, 2.2 mL of 0.45M solution in ACN, 1 mmol) were co-evaporated with anhydrous toluene (2 mL), suspended in anhydrous toluene (2 mL) and sonicated for 1 h at room temperature in a bath sonicator. The solvent was evaporated and the residue dissolved in anhydrous CH_2_Cl_2_ (3.2 mL). The stirred solution was cooled to 0 °C and diallyl *N,N*-diisopropylphosphoramidite (93 µL, 0.35 mmol) was added. After 20 min, the solution was cooled to −40 °C, and *m*CPBA (149 mg, 0.66 mmol) was added. After 30 min, the reaction was quenched with 1 M aq. Na_2_SO_3_ (8 mL), and warmed to room temperature. The mixture was diluted with CH_2_Cl_2_ (10 mL) and the layers were separated. The organic phase was washed with sat. aq. NaHCO_3_ (10 mL) and the combined aqueous phase was re-extracted with CH_2_Cl_2_ (2×20 mL). The combined organic phase was washed with brine (20 mL), dried over MgSO_4_, and concentrated. The residue was purified by medium-pressure flash chromatography (10g SNAP-KP-SIL; A: cyclohexane, B: EtOAc; 25 CV linear gradient from 30% to 95%, then 5 CV from 95%B) to give phosphate **SI-12** (70 mg, 0.12 mmol, 56%, α-anomer only) as a clear oil. ^1^H NMR (400 MHz, CDCl_3_) δ 6.41 (d, *J* = 9.1 Hz, 1H), 6.04 – 5.84 (m, 2H), 5.67 (dd, *J* = 6.3, 3.3 Hz, 1H), 5.47 – 5.12 (m, 6H), 4.58 (m, 4H), 4.43 (m, 1H), 4.28 – 4.16 (m, 2H), 4.14 – 4.05 (m, 1H), 3.56 (m, 2H), 2.47 – 2.32 (m, 2H), 2.06 (s, 3H), 2.01 (d, *J* = 2.3 Hz, 6H). ^13^C NMR (100 MHz, CDCl_3_) δ 171.2, 170.6, 170.2, 169.2, 132.2, 132.1, 132.0, 131.9, 119.1, 96.0, 70.0, 69.8, 68.9, 68.8, 68.8, 68.7, 67.5, 61.4, 51.9, 51.8, 47.0, 35.4, 29.7, 20.7, 20.6, 20.5; LRMS (ESI) calcd. for C_21_H_31_N_4_O_12_P (M-H^+^) 562.17 found 561.18 *m/z*.

### Uridine 5’-diphospho-2-(3-azidopropionamido)-2-deoxy-α-D-glucopyranoside ammonium salt (SI-13)

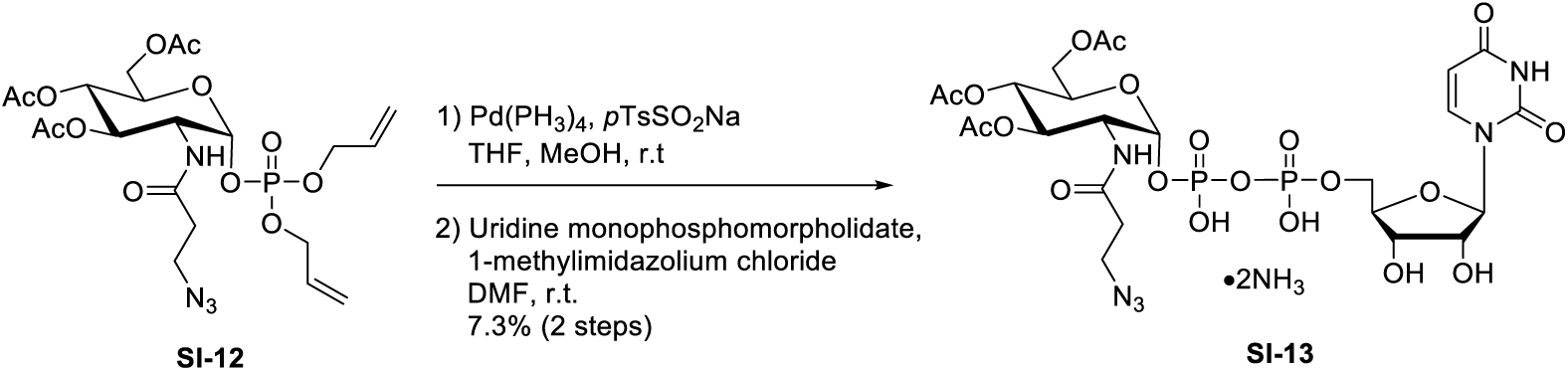

To a stirred solution of diallyl phosphotriester **SI-12** (66 mg, 117 µmol) in THF/MeOH (1:1, 2.4 mL) were added tetrakis(triphenylphosphine)palladium (14 mg, 12 µmol) and sodium *para*-toluoenesulfinate (84 mg, 468 µmol). The mixture was stirred at room temperature for 16 h, and the solvents were evaporated. The residue was co-evaporated with anhydrous toluene (2×10 mL) and dissolved in DMF (2.4 mL). Uridine monophosphomorpholidate (128 mg, 187 µmol) and 1-methylimidazolium chloride (75 mg, 0.64 µmol) were added and the reaction was stirred at room temperature for 16 h. Additional uridine monophosphomorpholidate (128 mg, 187 µmol) and 1-methylimidazolium chloride (75 mg, 0.64 µmol) were added to the mixture and left to react for 16 h. The mixture was concentrated and purified by medium-pressure flash chromatography (30 g SNAP C18 column; A: 10 mM ammonium acetate, B: MeOH; 5 CV 100% A; 20 CV linear gradient to 100% B, then 5 CV 100% B) and then by preparative HPLC (ZORBAX 300SB-C8; A: 10 mM Ammonium Acetate, B: MeOH; linear gradient from 0 to 60% B over 40 min). The fractions were concentrated and lyophilized to give pyrophosphate **SI-13** as the ammonium salt (7 mg, 8.5 µmol, 7.3% over two steps) as a white foam. ^1^H NMR (400 MHz, CD_3_OD) δ 8.09 (d, *J* = 8.1 Hz, 1H), 5.97 (d, *J* = 4.5 Hz, 1H), 5.86 (d, *J* = 8.1 Hz, 1H), 5.63 (dd, *J* = 7.3, 3.3 Hz, 1H), 5.31 (dd, *J* = 10.6, 9.4 Hz, 1H), 5.14 (dd, *J* = 10.2, 9.4 Hz, 1H), 4.45 – 4.36 (m, 3H), 4.35 (m, 1H); 4.31 (m, 2H), 4.22 – 4.16 (m, 2H), 3.66 – 3.49 (m, 2H), 2.69 – 2.57 (m, 2H), 2.07 (s, 3H), 2.02 (s, 3H), 1.97 (s, 3H). ^13^C NMR (100 MHz, CD_3_OD) δ 172.3, 171.1, 170.6, 170.0, 164.9, 151.3, 141.3, 101.7, 94.5, 88.9, 83.6, 81.4, 74.4, 71.7, 69.5, 68.5, 68.3, 64.6, 61.5, 51.7, 39.0, 34.6, 19.3; LRMS (ESI) calcd. for C_24_H_33_N_6_O_20_P_2_ (M-H^+^) 787.13 found 787.10 *m/z*.

### Uridine 5’-diphospho-2-(3-azidopropionamido)-2-deoxy-α-D-glucopyranoside sodium salt SI-14

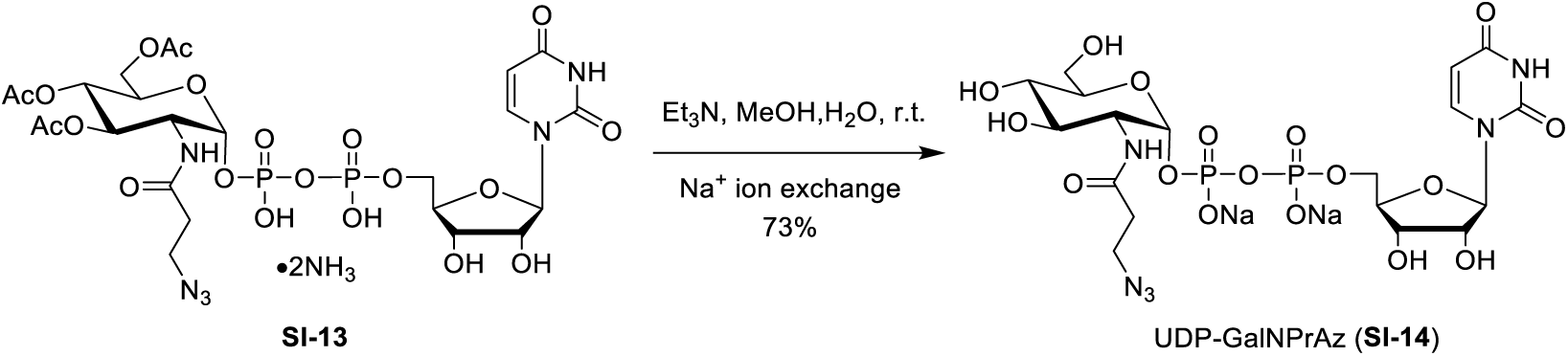

To a stirred solution of triester **SI-13** (7 mg, 8.5 µmol) in MeOH/water (5:2, 1.8 mL) was added triethylamine (375 µL). The reaction was stirred at room temperature for 16 h. The residue was concentrated and lyophilized repeatedly. The residue was passed through a short (4 g resin) ion exchange column of Dowex 50W X8 Na^+^ form (Serva, Heidelberg, Germany), and concentrated. The residue was purified by reverse-phase solid-phase extraction (Sep-Pak C18, 5g, Waters) and lyophilized to give UDP-GlcNAc analog **SI-14** as the disodium salt (4.4 mg, 6.2 µmol, 73%) as a white solid. ^1^H NMR (400 MHz, D_2_O) δ 7.80 (d, *J* = 8.1 Hz, 1H), 5.88 – 5.76 (m, 2H), 5.36 (m, 1H), 4.26 – 4.17 (m, 2H), 4.12 (m, 1H), 4.04 (m, 2H), 3.88 (m, 1H), 3.78 (m, 1H), 3.71 (dd, *J* = 12.5, 2.4 Hz, 1H), 3.66 – 3.62 (m, 2H), 3.47 – 3.36 (m, 3H), 2.49 (m, 2H). ^13^C NMR (100 MHz, D_2_O) δ 174.1, 166.4, 151.9, 141.6, 102.6, 94.6, 88.5, 83.1, 73.8, 73.0, 70.9, 69.6, 69.5, 65.0, 60.3, 53.7, 47.1, 34.8; LRMS (ESI) calcd. for C_18_H_28_N_6_O_17_P_2_ (M-H^+^) 661.10 found 661.10 *m/z*.

#### ^1^H NMR (400 MHz, acetone-D_6_)

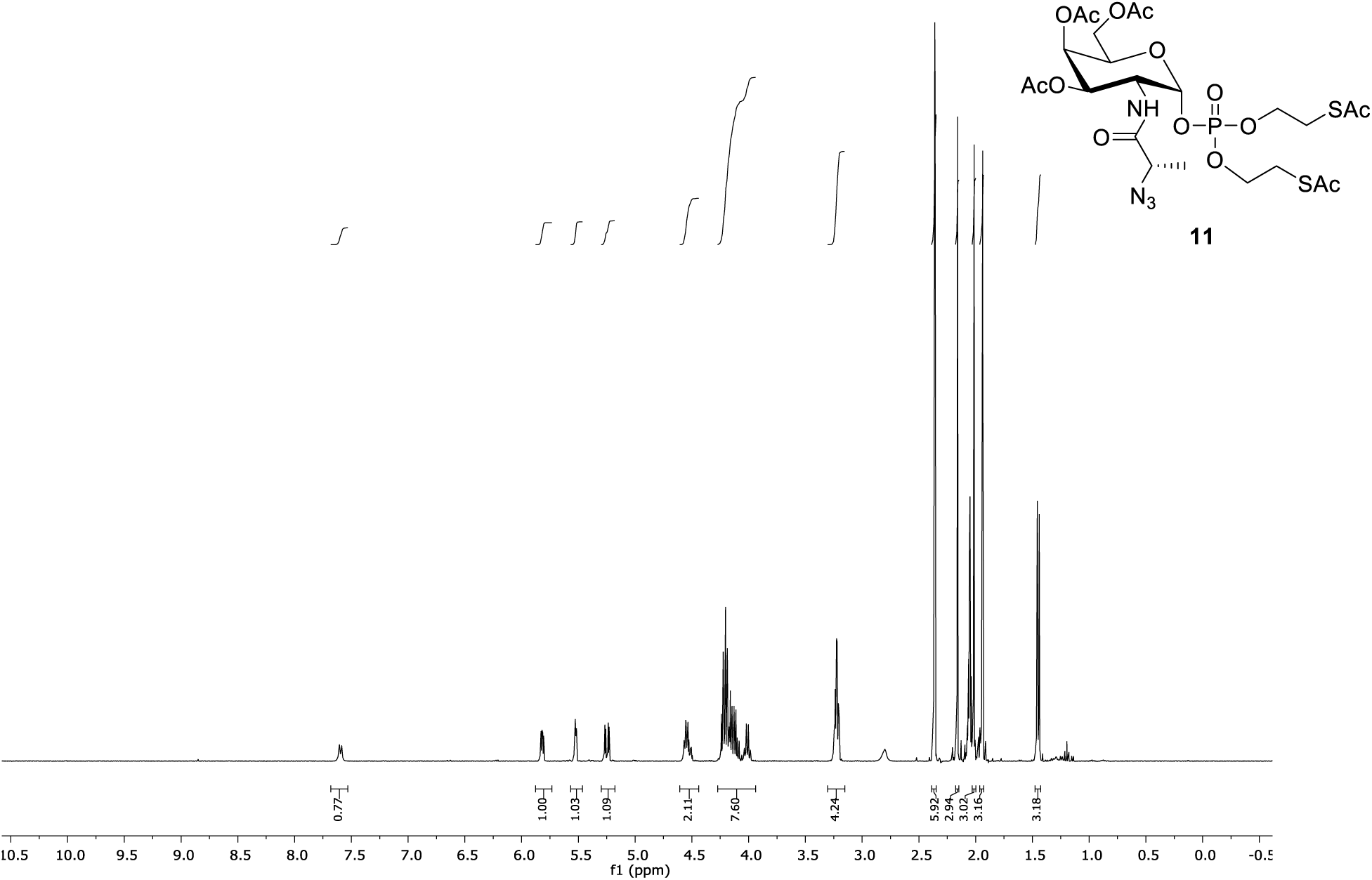

#### ^1^H NMR (400 MHz, CDCl_3_)

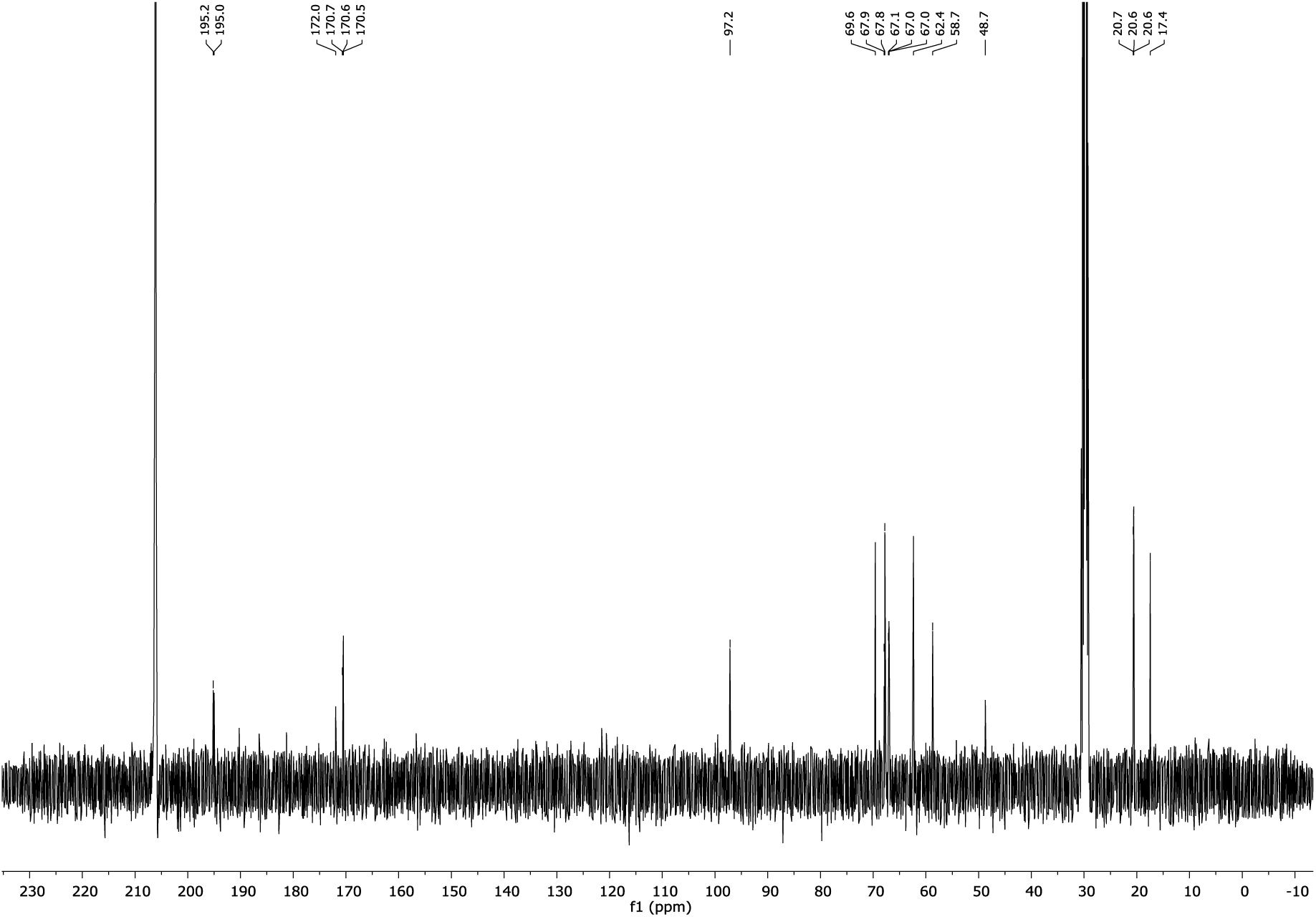

#### ^1^H NMR (400 MHz, CDCl_3_)

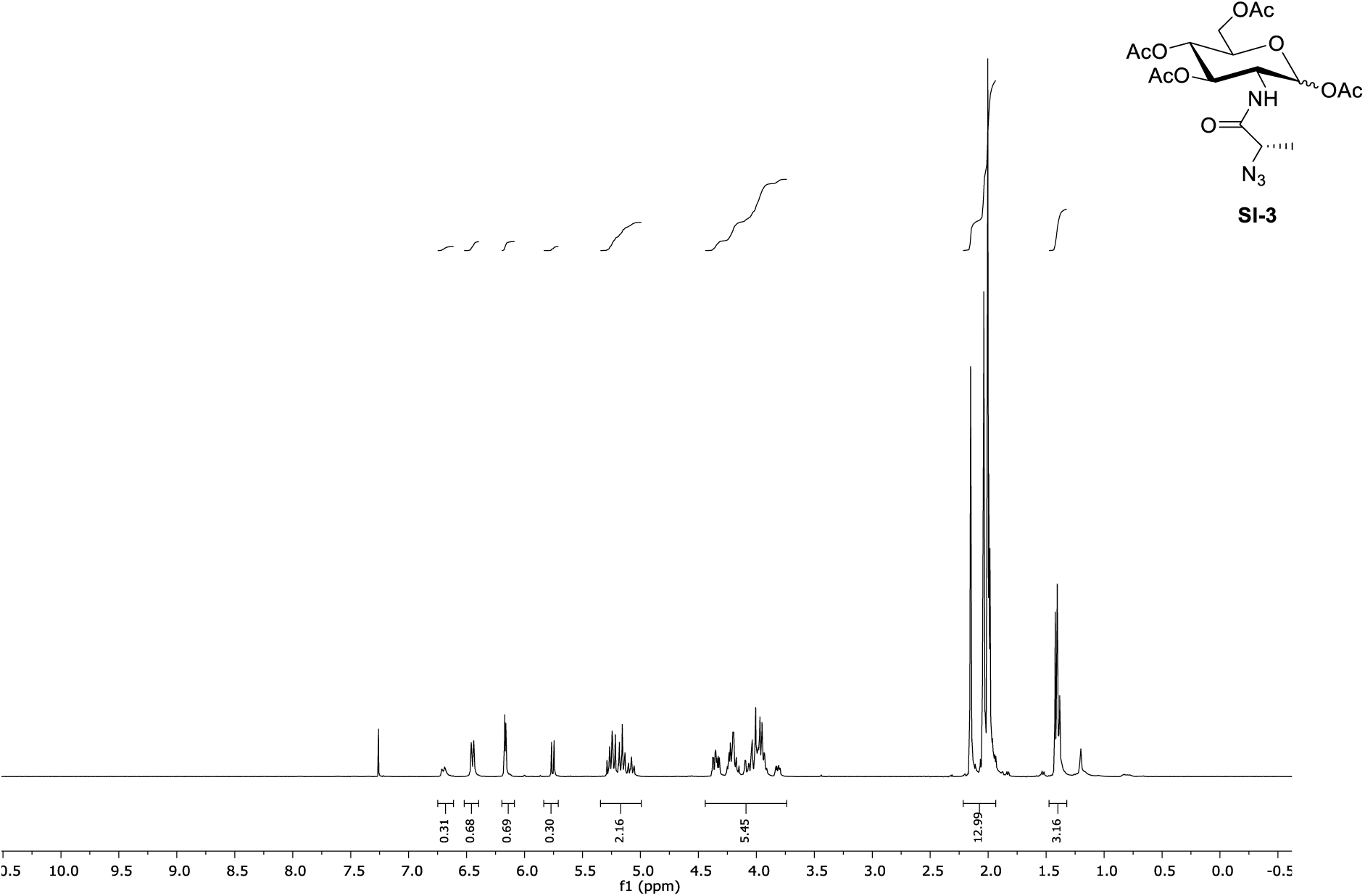

#### ^13^C NMR (100 MHz, CDCl_3_)

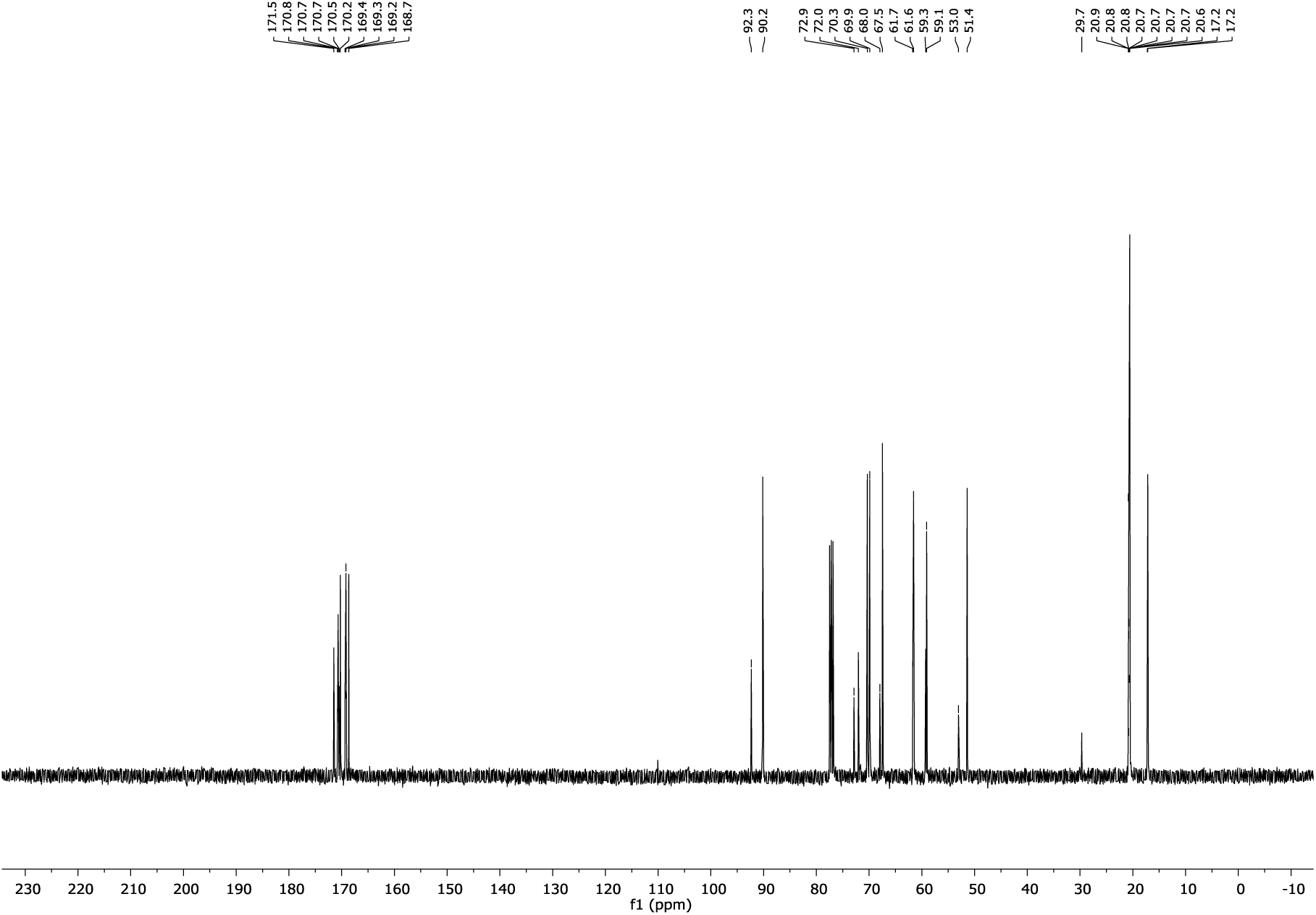

#### ^1^H NMR (400 MHz, CDCl_3_)

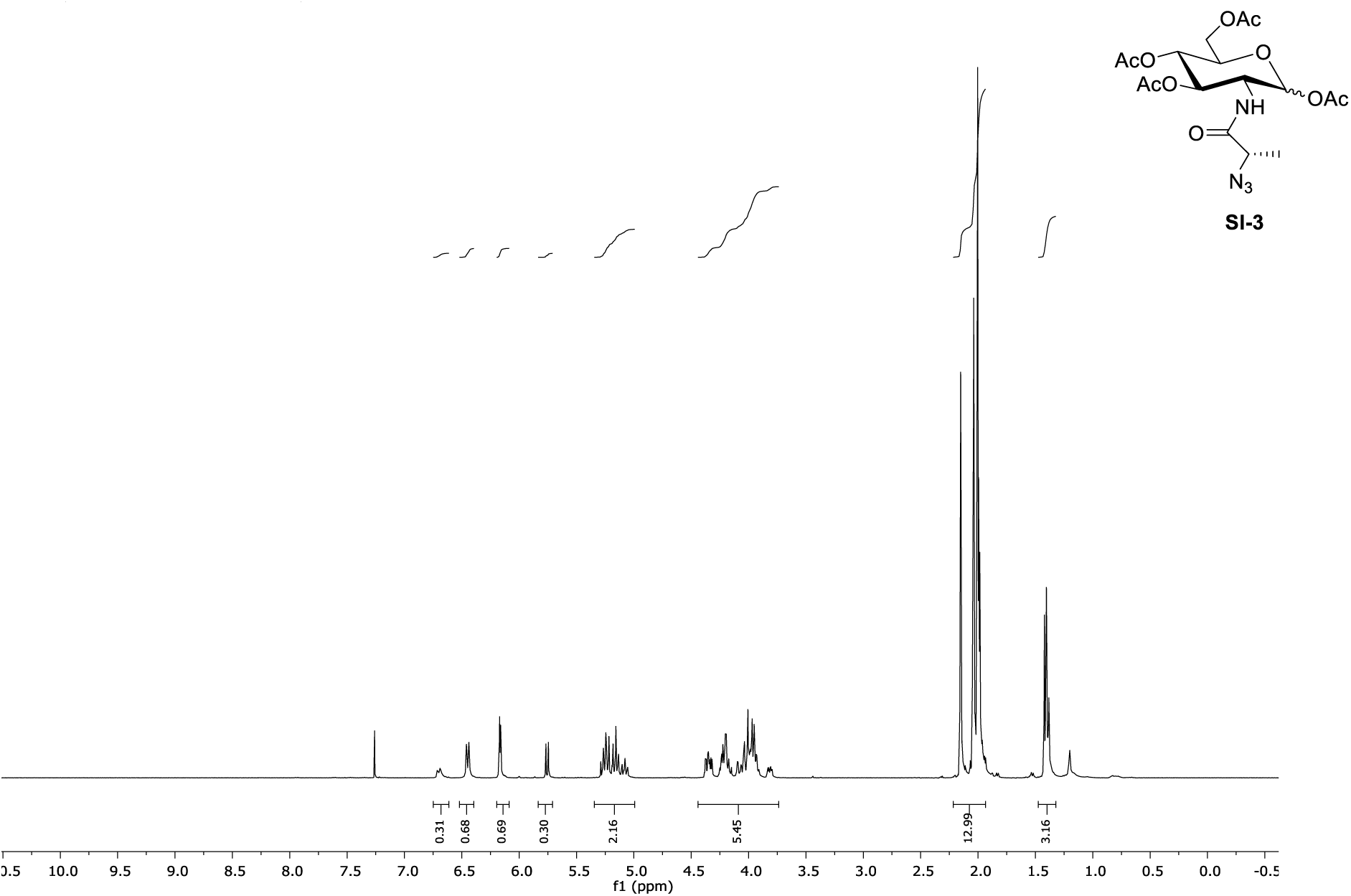

#### ^13^C NMR (100 MHz, CDCl_3_)

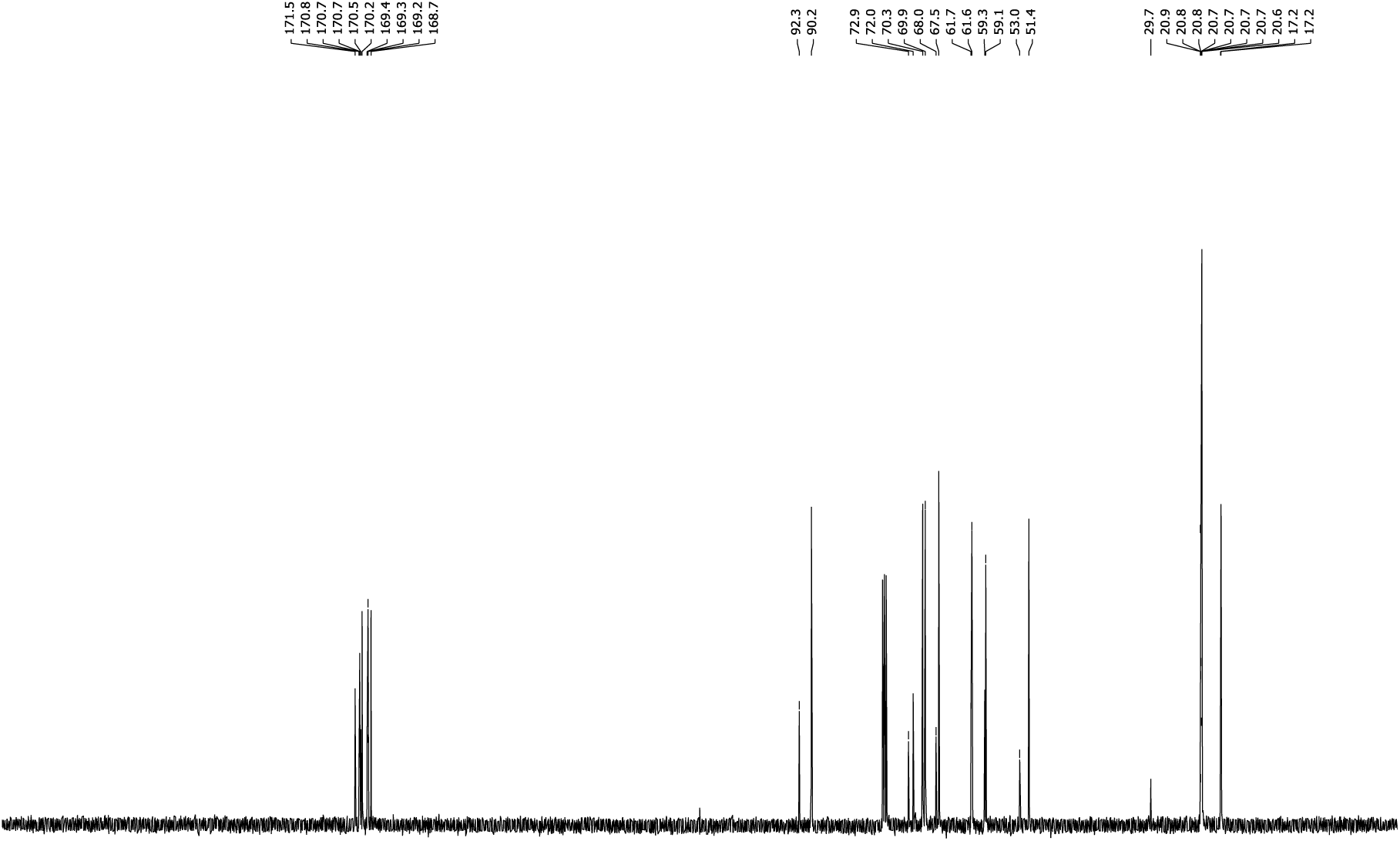

#### ^1^H NMR (400 MHz, CDCl_3_)

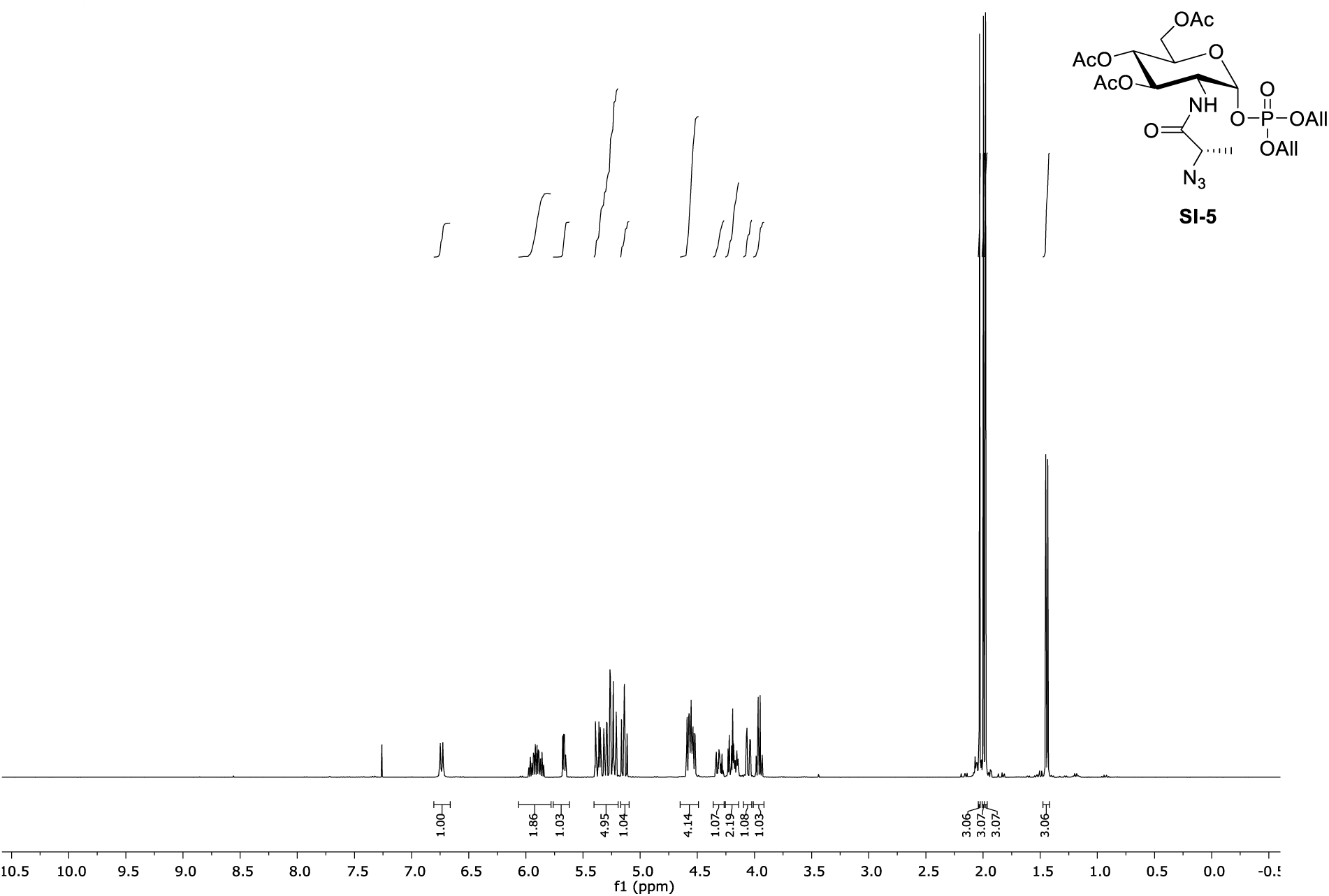

#### ^13^C NMR (100 MHz, CDCl_3_)

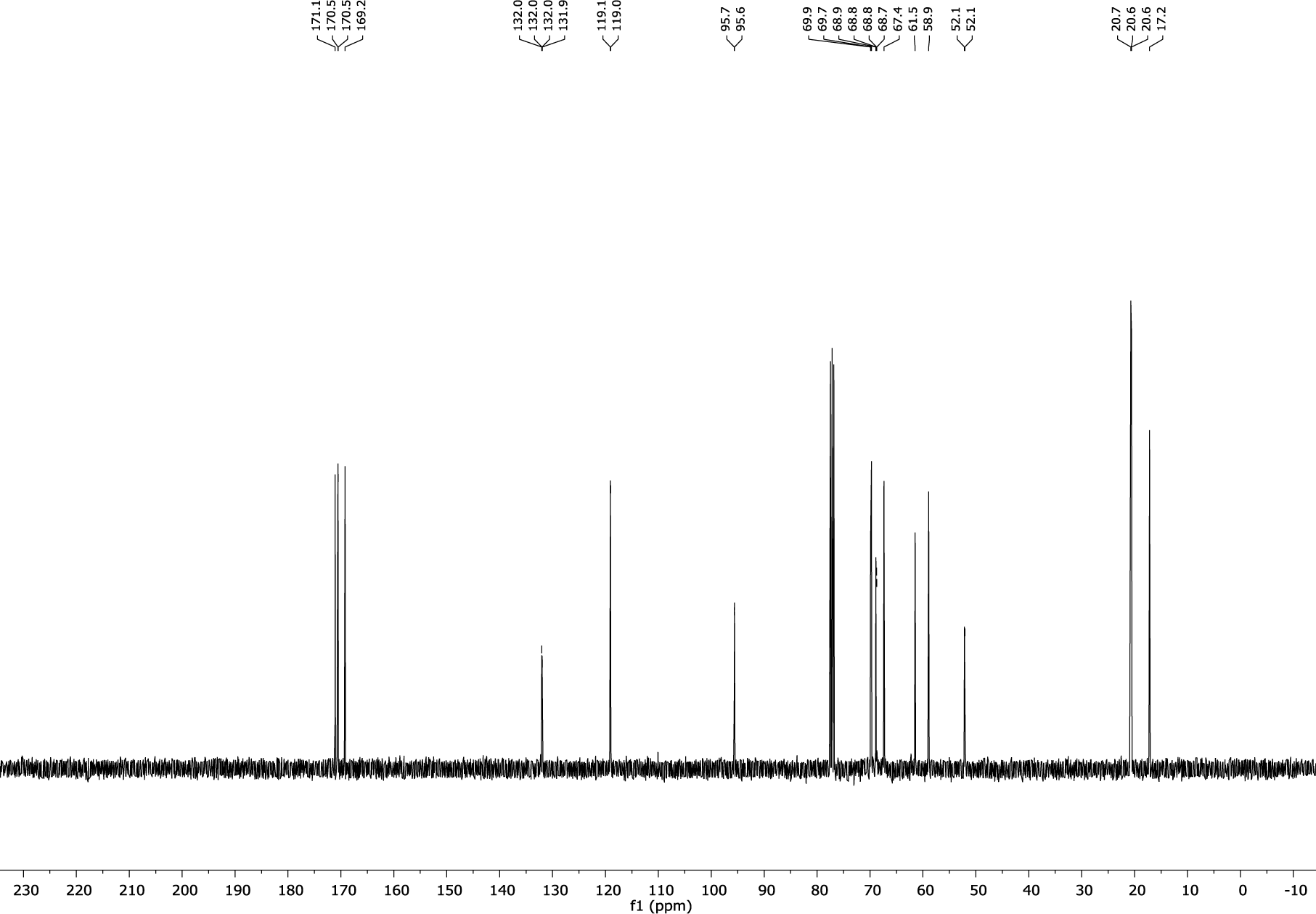

#### CH-HSQC NMR (400 MHz, CDCl_3_)

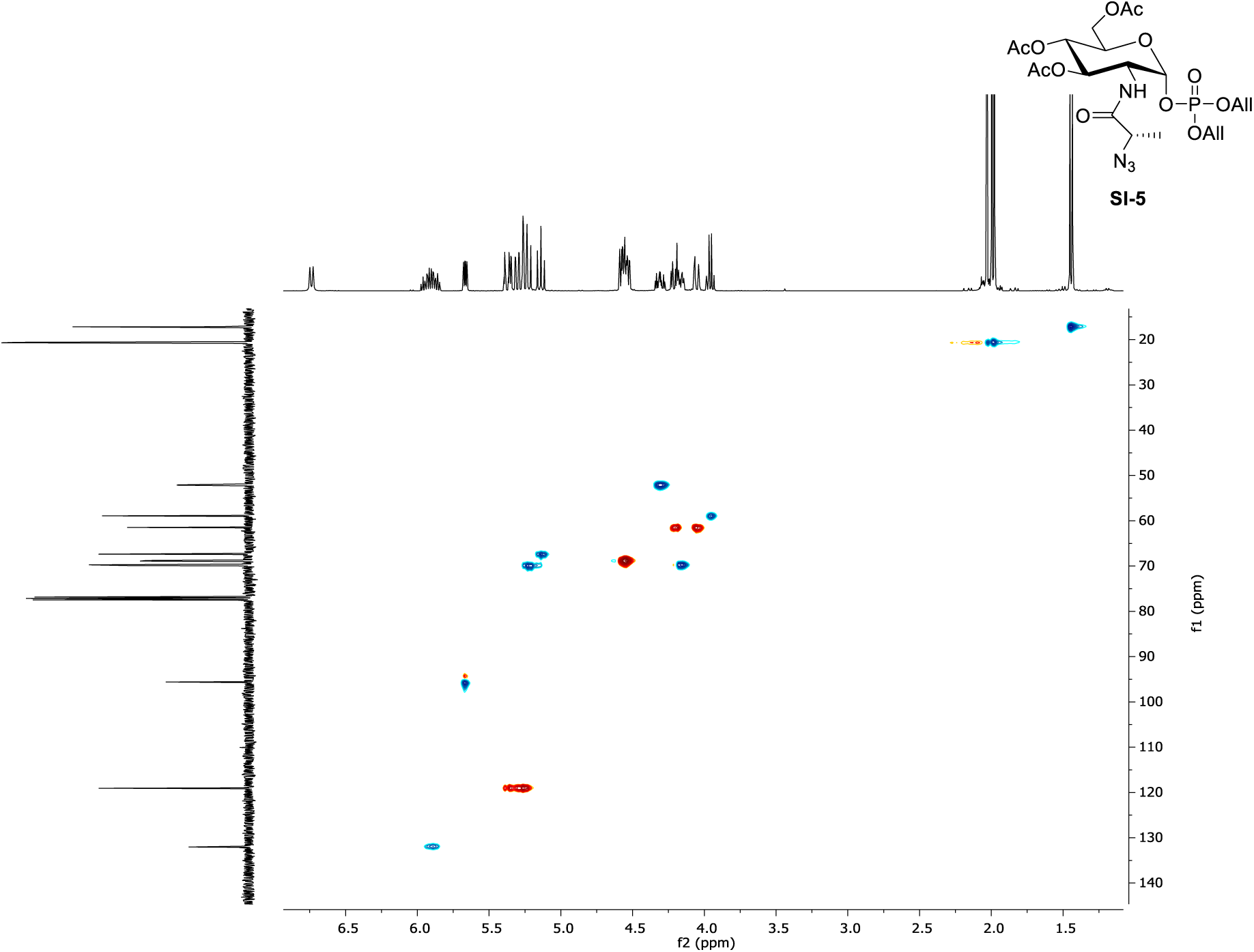

#### HH-COSY NMR (400 MHz, CDCl_3_)

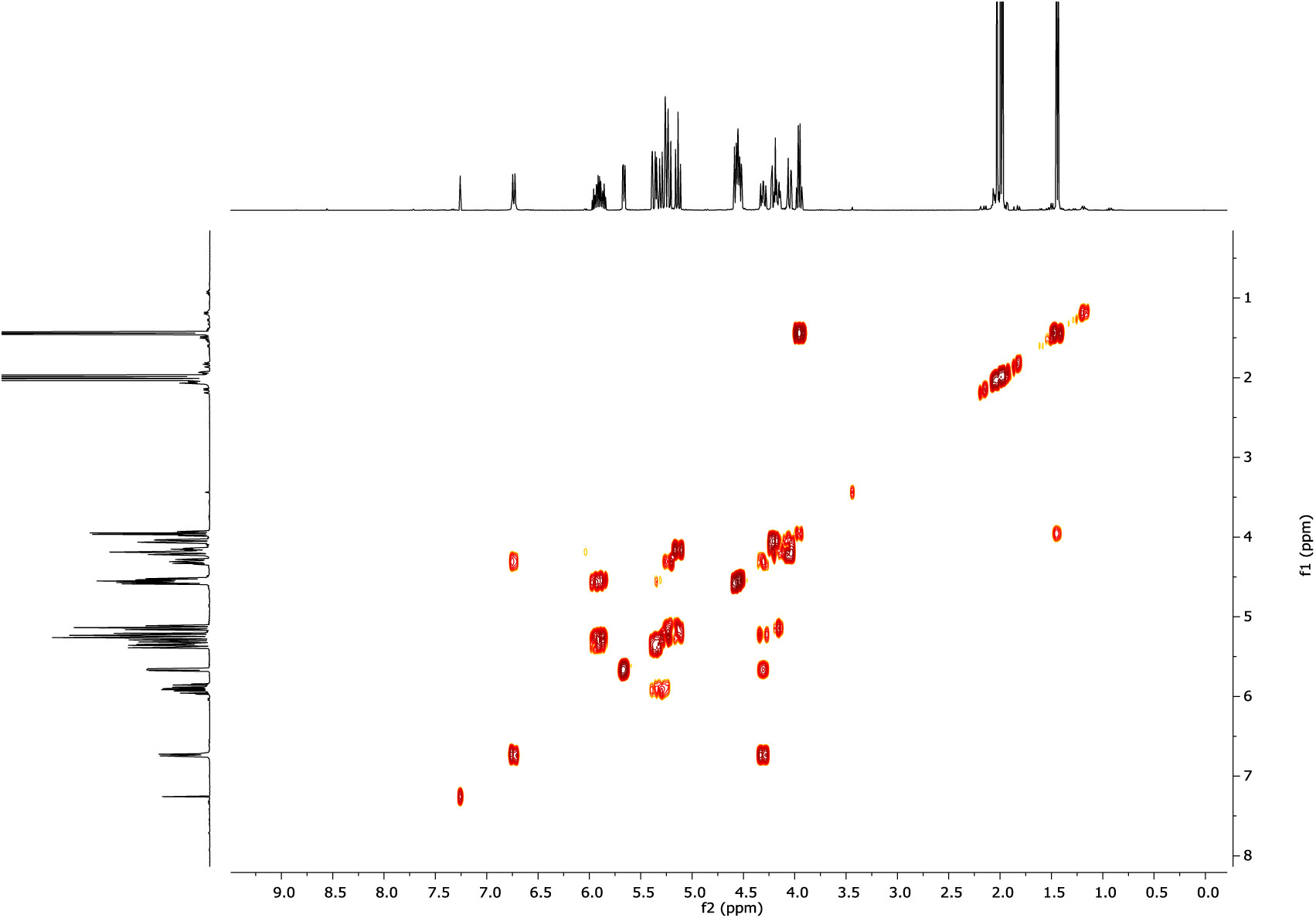

#### ^1^H NMR (600 MHz, CD_3_OD)

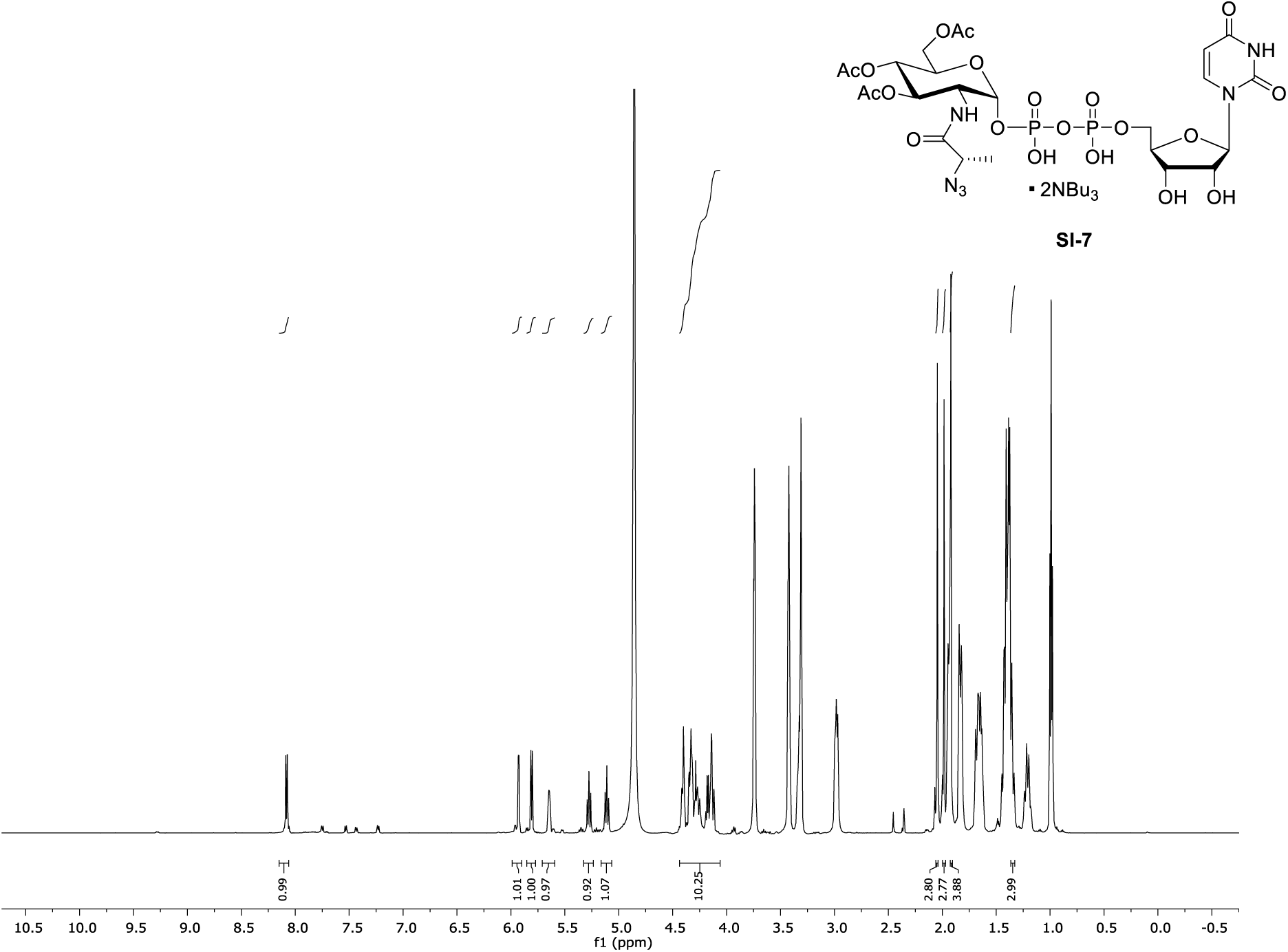

#### ^13^C NMR (150 MHz, CD_3_OD)

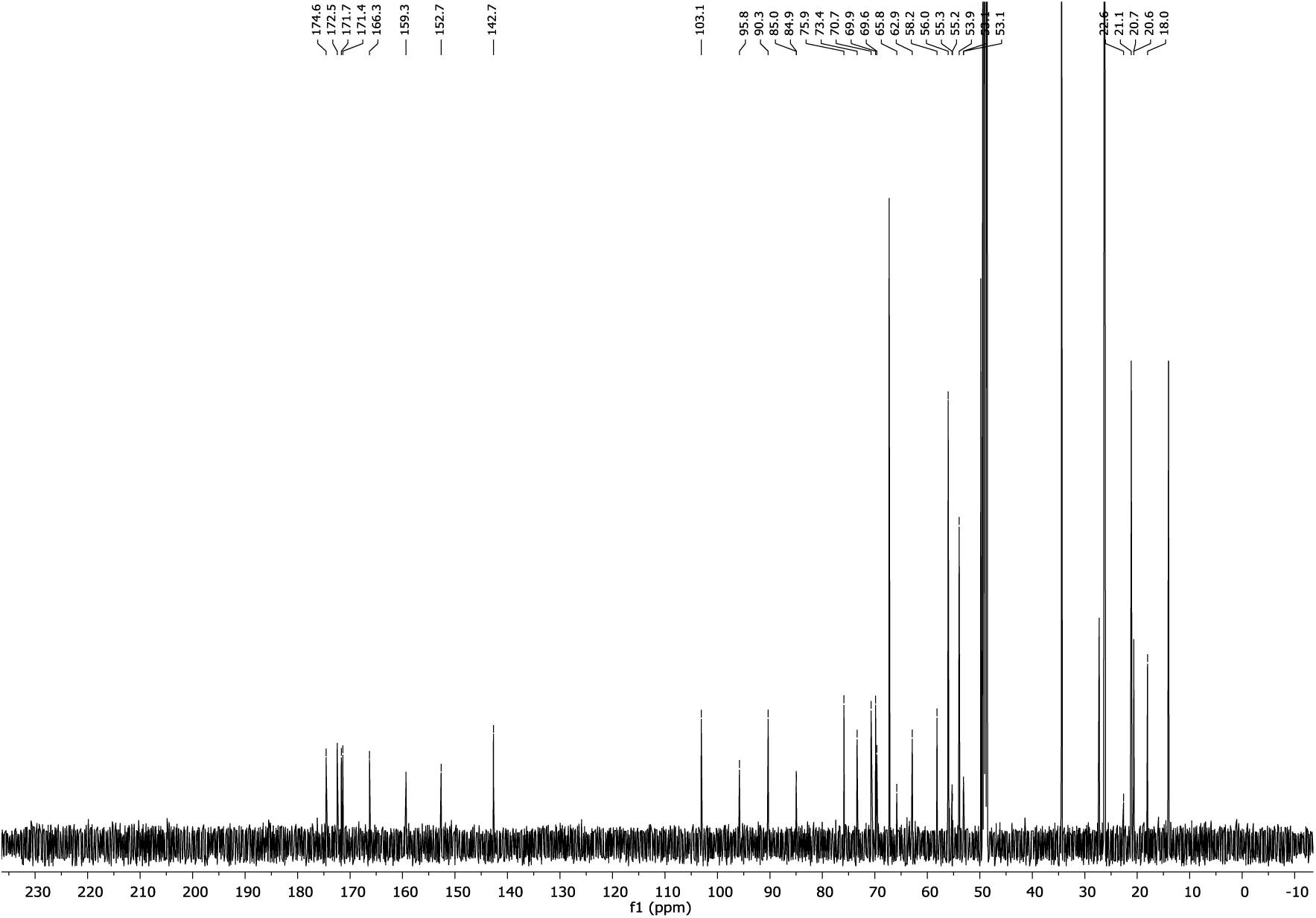

#### ^1^H NMR (600 MHz, CD_3_OD)

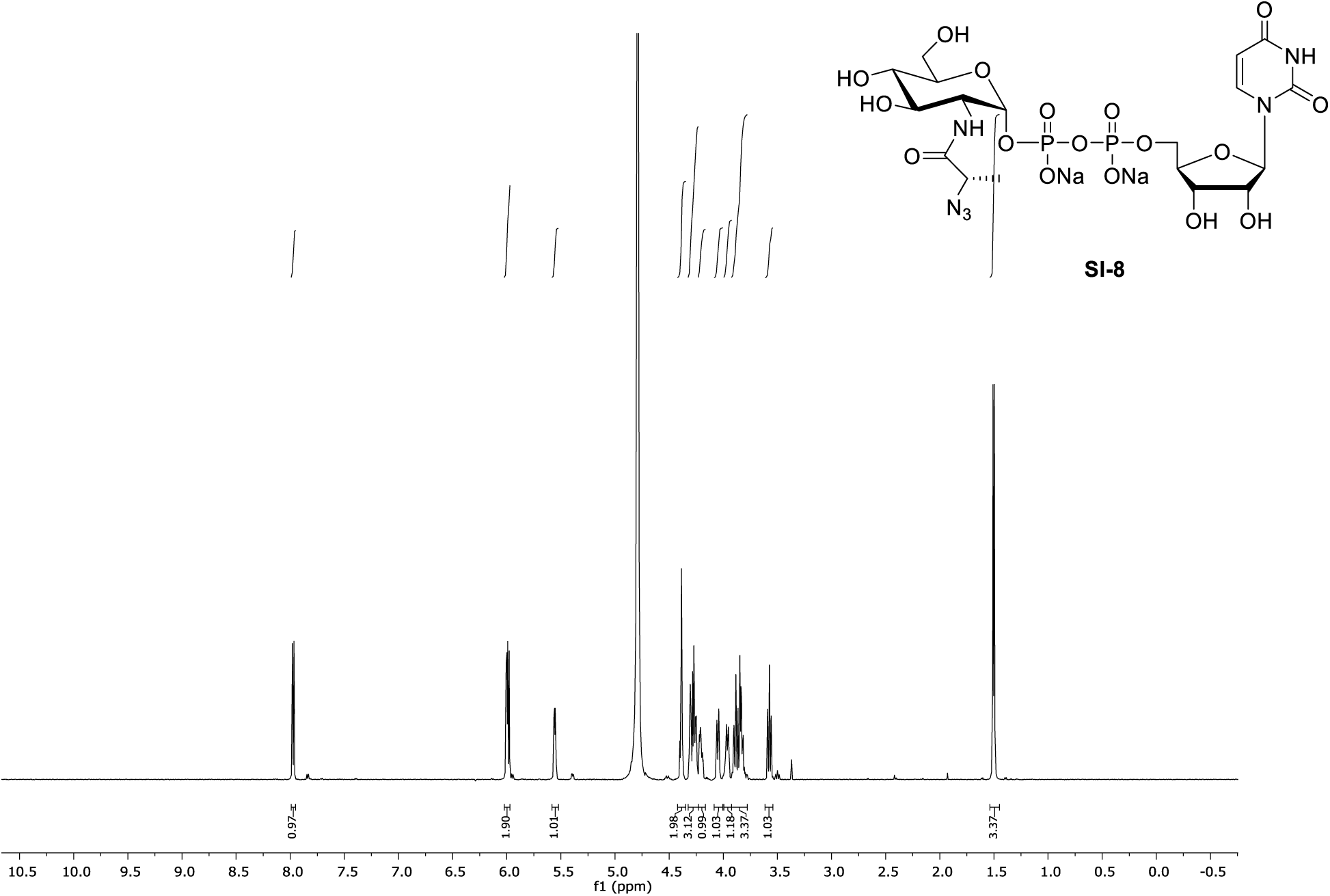

#### ^13^C NMR (150 MHz, D_2_O)

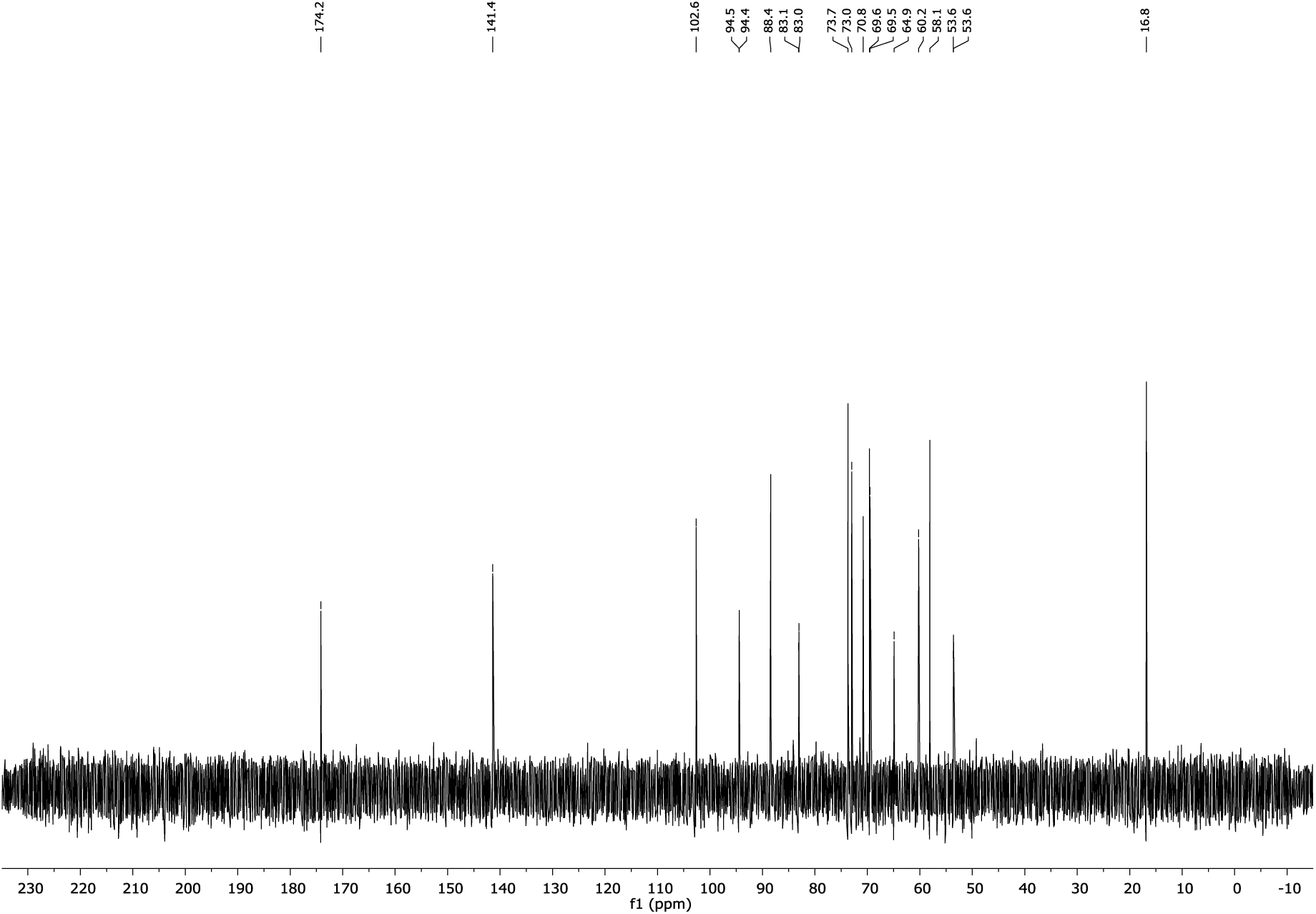

#### HH-COSY NMR (600 MHz, D_2_O)

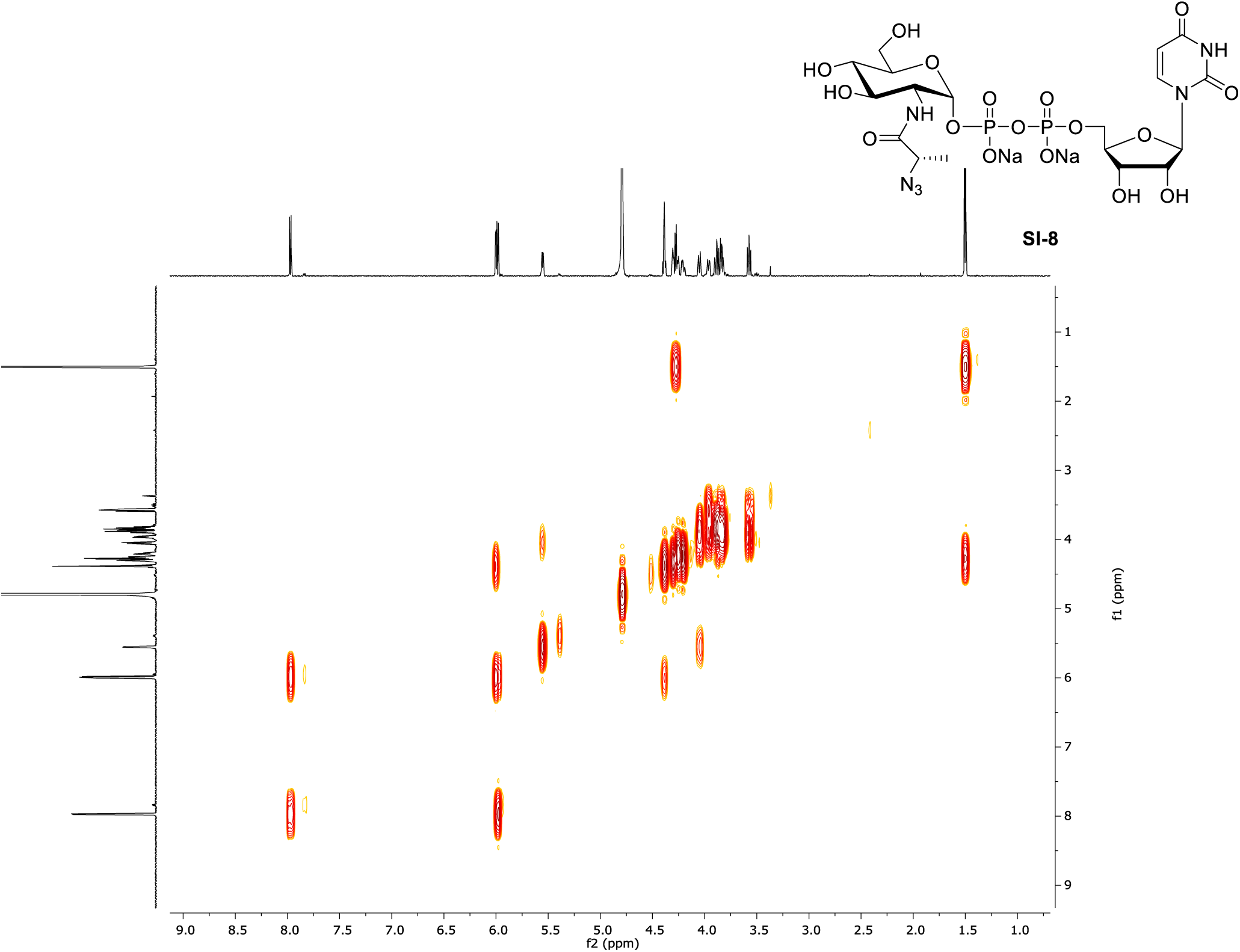

#### ^1^H NMR (400 MHz, CDCl_3_)

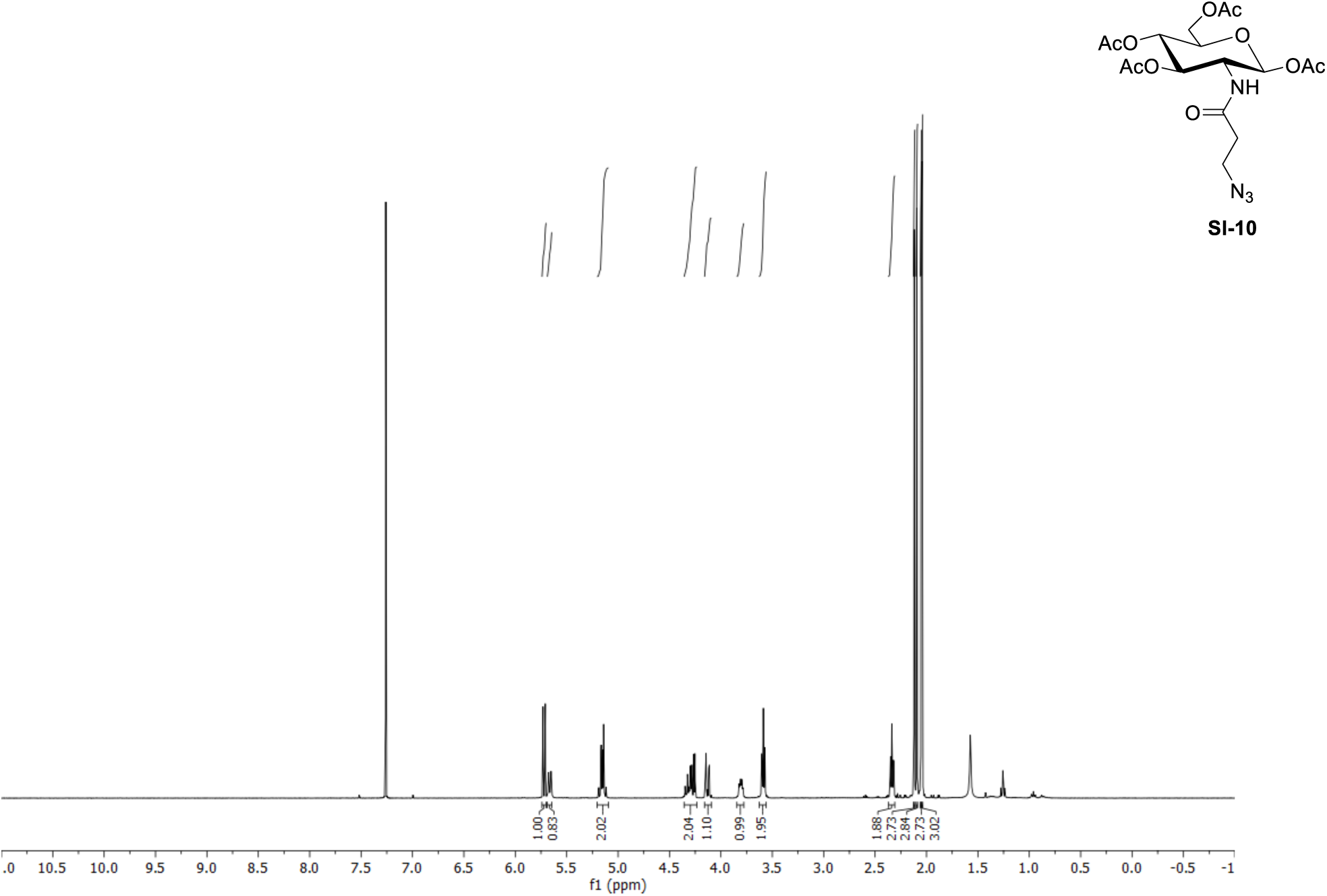

#### ^13^C NMR (100 MHz, CDCl_3_)

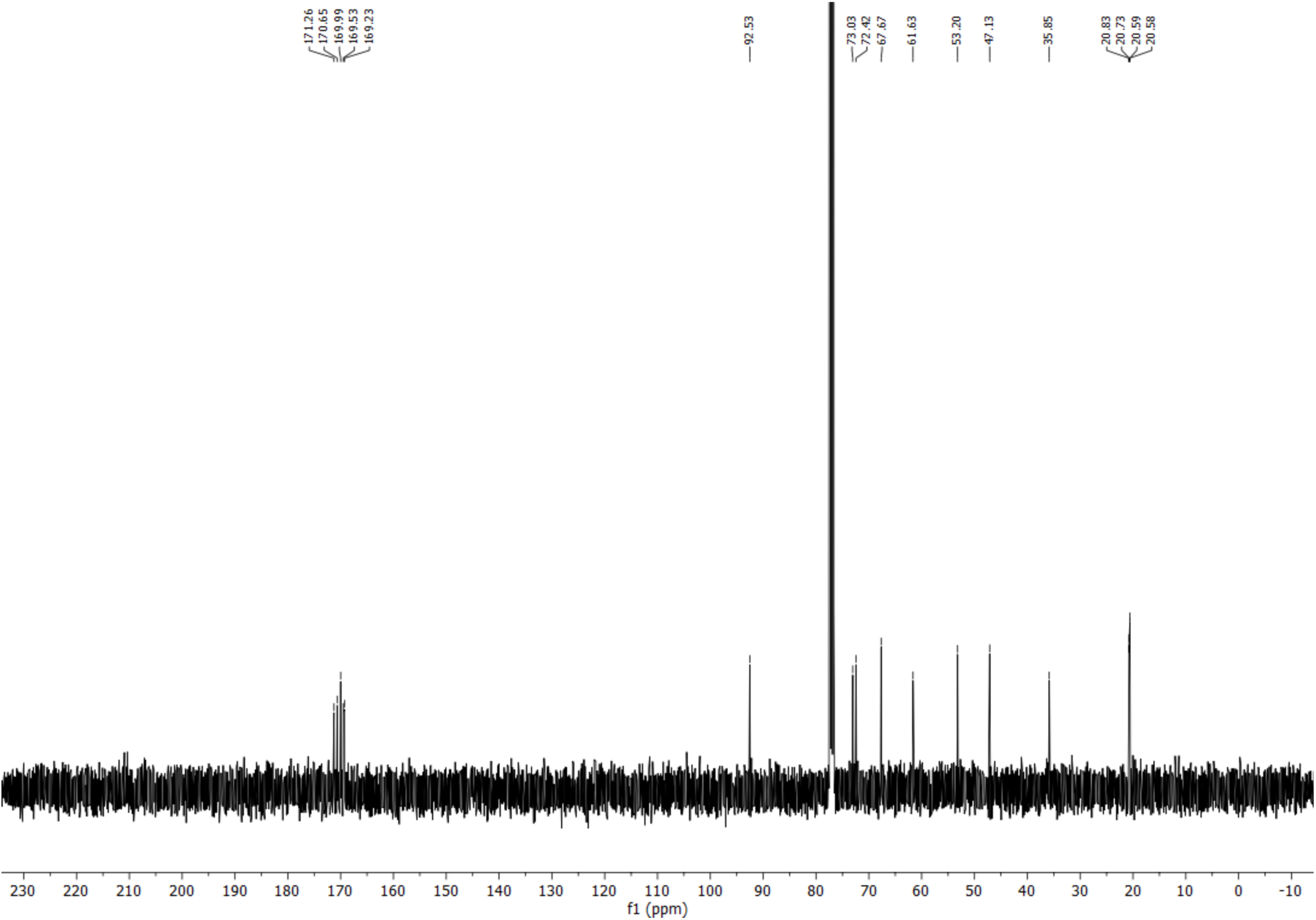

#### ^1^H NMR (400 MHz, CDCl_3_)

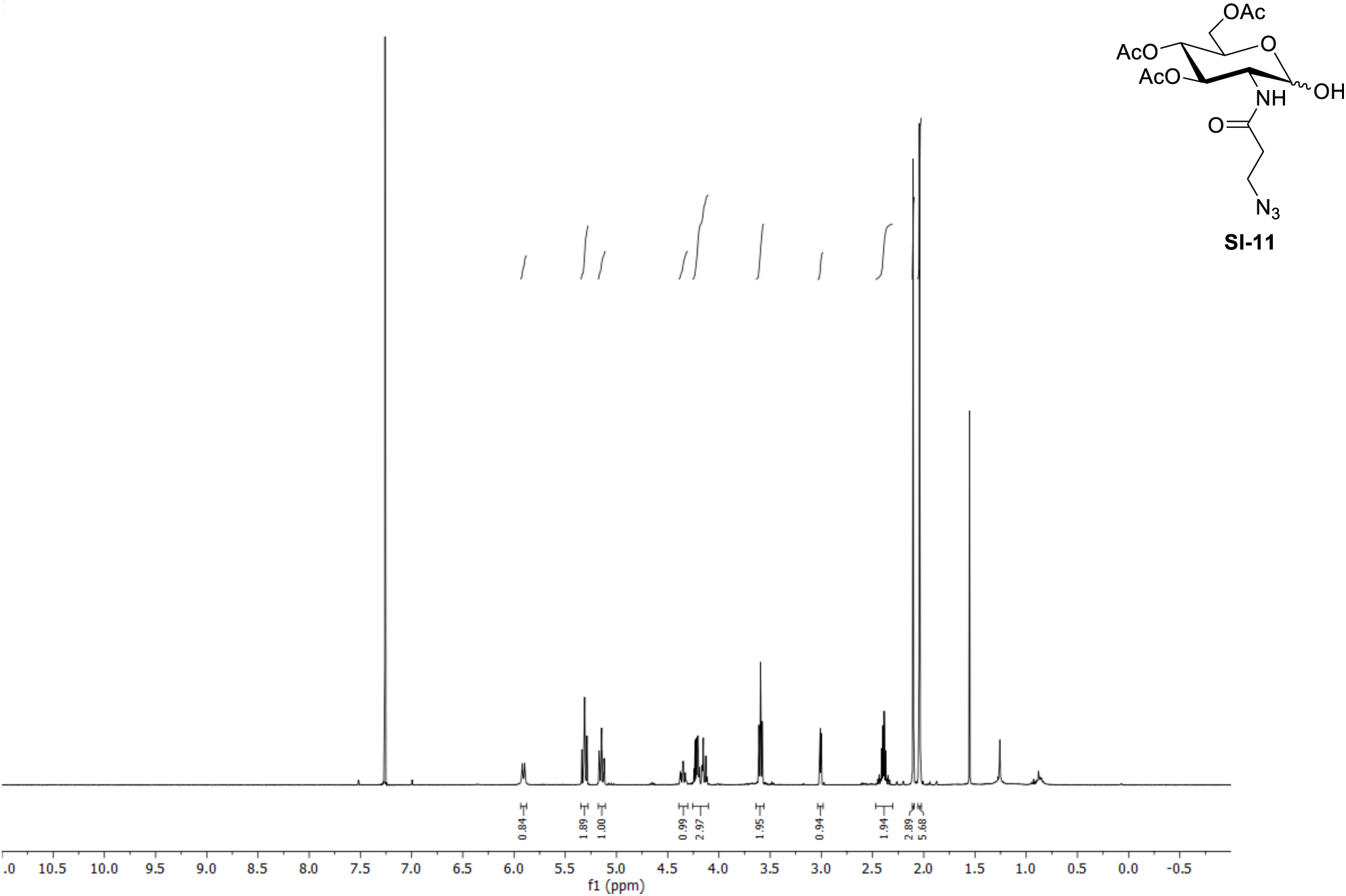

#### ^13^C NMR (100 MHz, CDCl_3_)

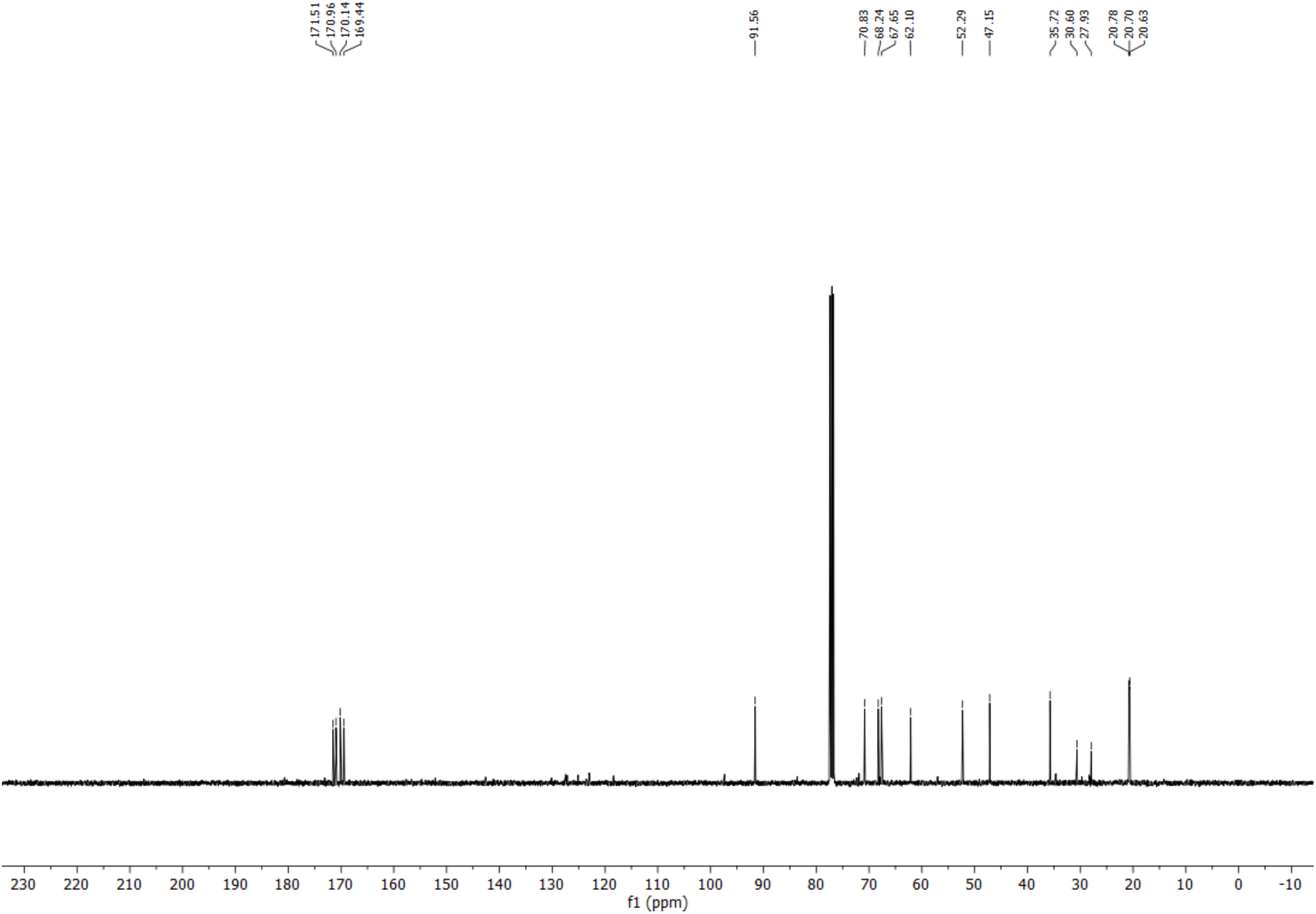

#### ^1^H NMR (400 MHz, CDCl_3_)

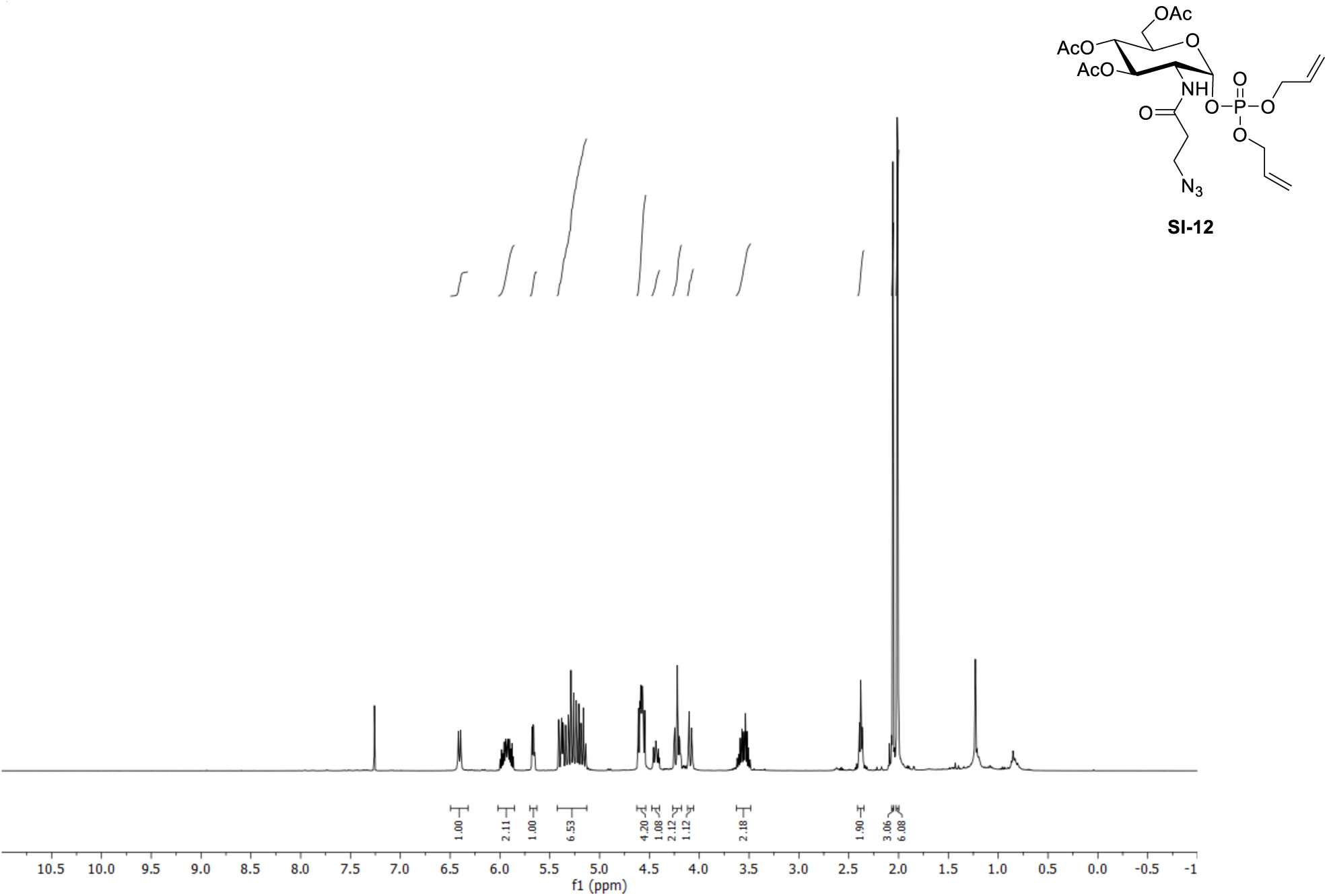

#### ^13^C NMR (100 MHz, CDCl_3_)

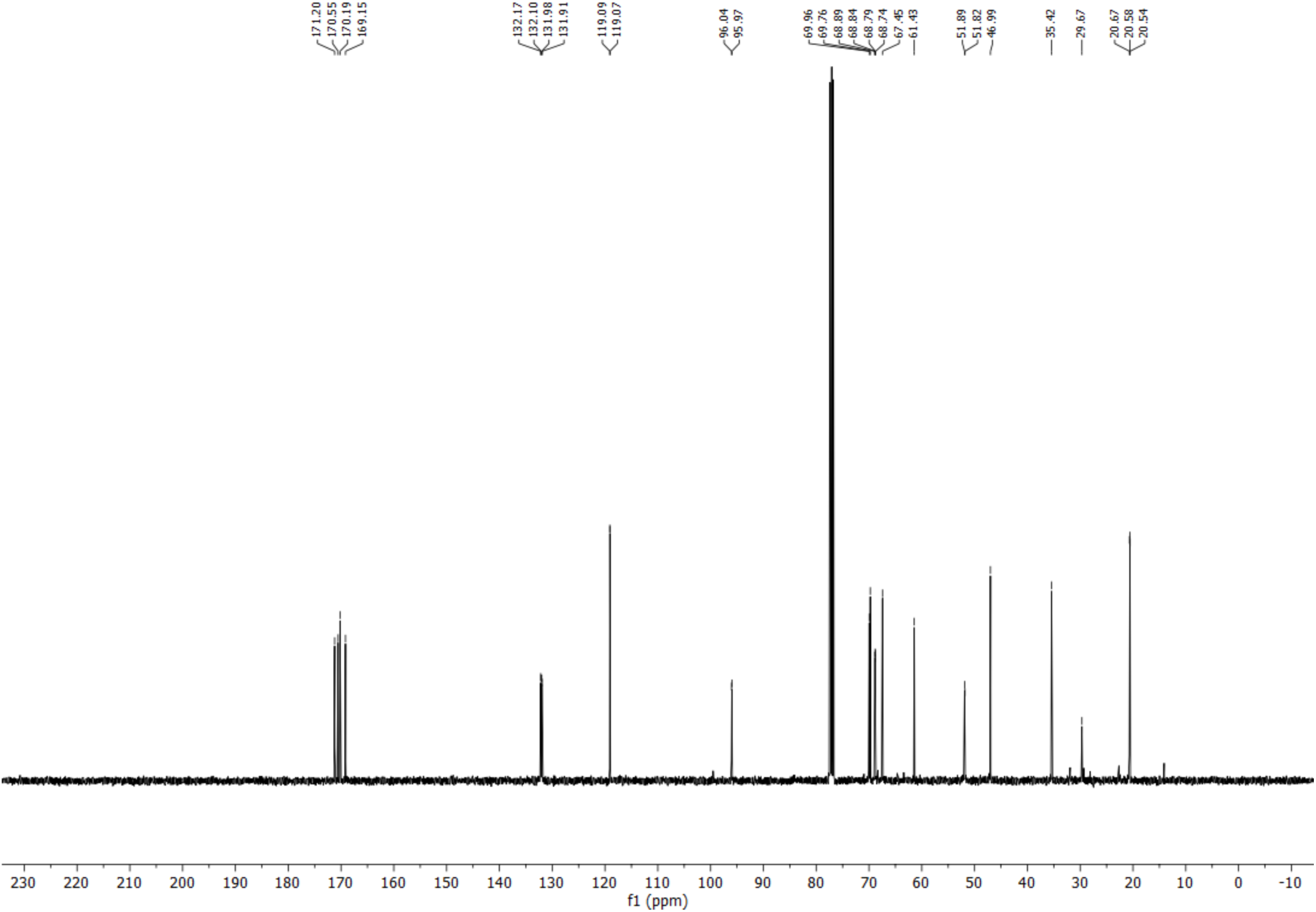

#### ^1^H NMR (400 MHz, CDCl_3_)

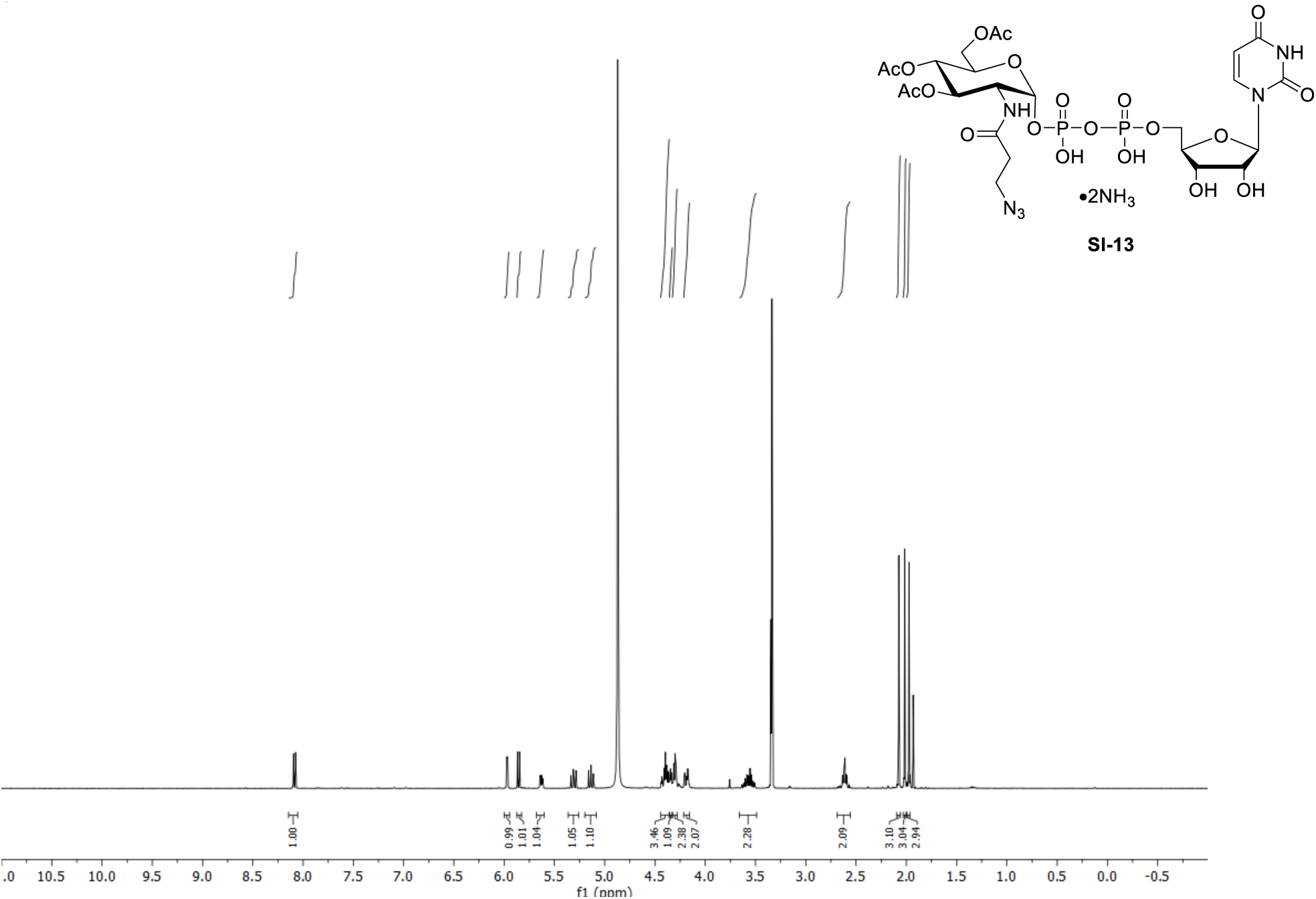

#### ^13^C NMR (100 MHz, CDCl_3_)

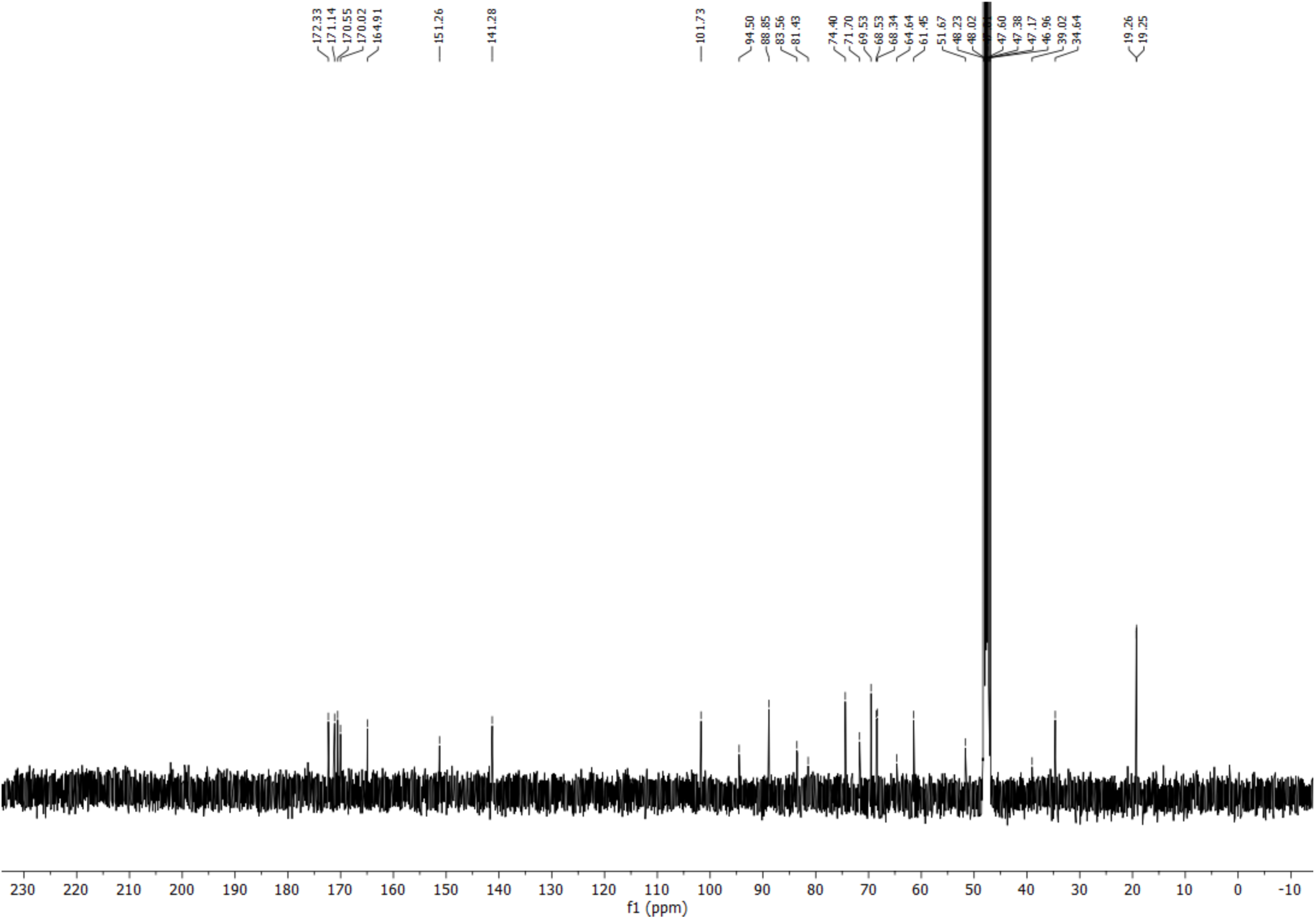

#### ^1^H NMR (400 MHz, CDCl_3_)

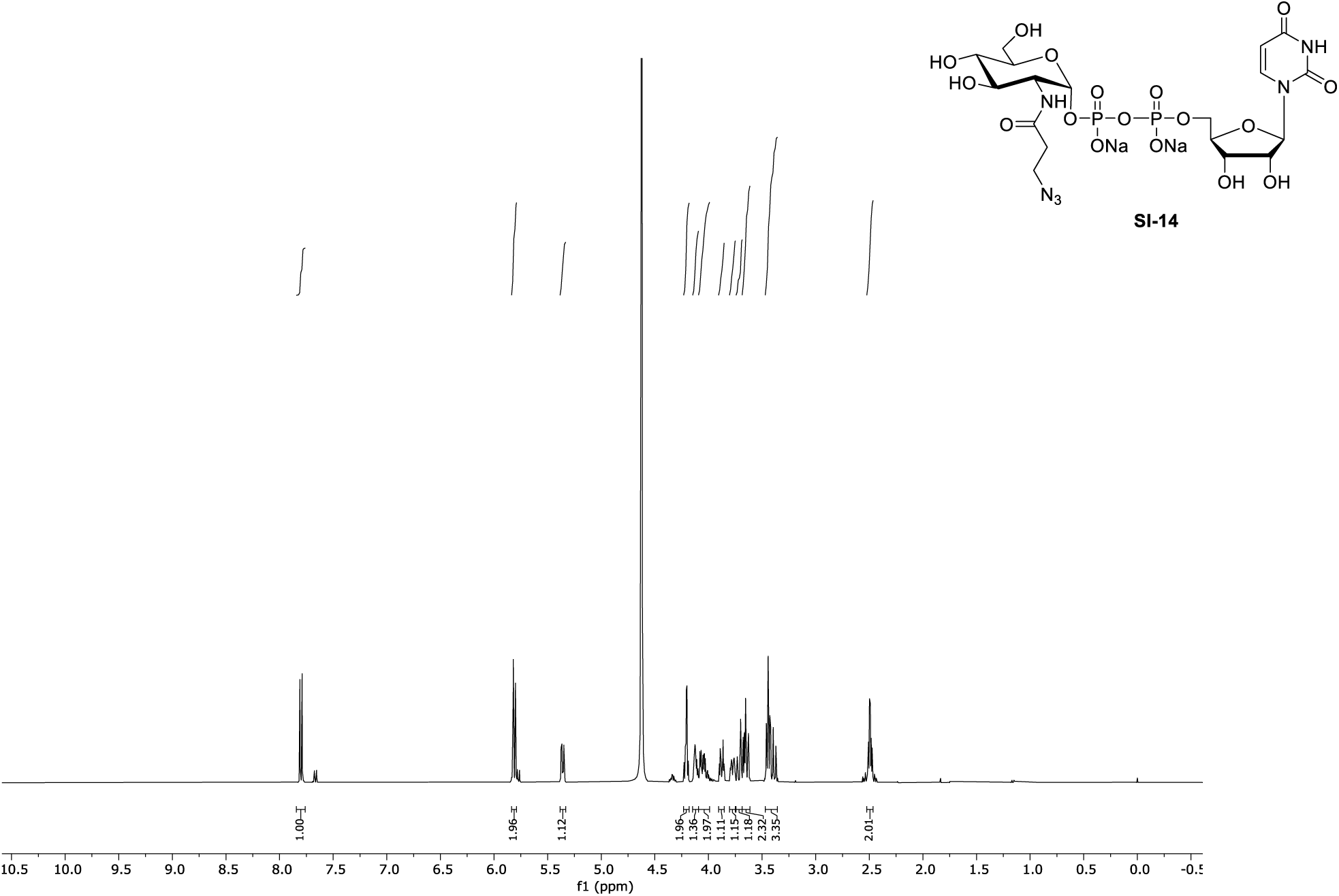

#### ^13^C NMR (101 MHz, CDCl_3_)

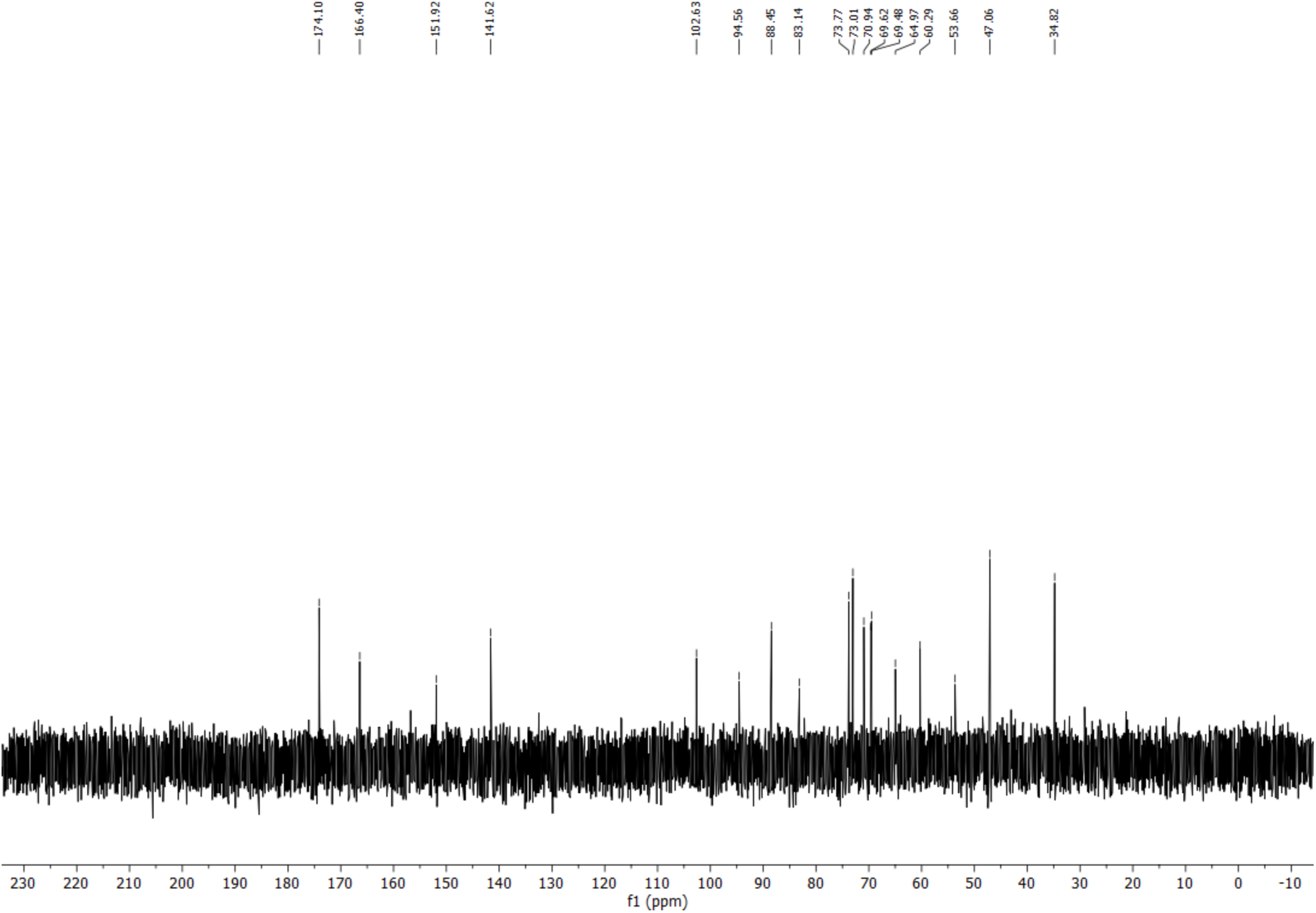

## Notes

### Competing Interest Statement

The authors have declared no competing interest.

